# Emergent Neuronal Mechanisms Mediating Covert Attention in Convolutional Neural Networks

**DOI:** 10.1101/2023.09.17.558171

**Authors:** Sudhanshu Srivastava, William Yang Wang, Miguel P. Eckstein

## Abstract

Covert visual attention allows the brain to select different regions of the visual world without eye movements. Predictive cues of a target location orient covert attention and improve perceptual performance. In most computational models, researchers explicitly incorporate an attentional mechanism that alters processing at the attended location (gain, noise reduction, divisive normalization, biased competition, Bayesian priors). Here, we assess the emergent neuronal mechanisms of Convolutional Neural Networks (CNNs) that exhibit behavioral signatures of covert attention, despite lacking a built-in attention mechanism. We use neuroscience-inspired approaches to analyze 1.8M units of CNNs trained on the cueing paradigm. Consistent with neurophysiology, we show early layers with retinotopic neurons separately tuned to the target or cue, and later layers with neurons with joint tuning and increased cue influence on target responses. CNN computational stages mirror a Bayesian ideal observer (BIO), but with more gradual transitions. The cue influences the target sensitivity through four mechanisms. A BIO-like cue-weighted location summation, and three mechanisms absent in the BIO: an opponency across locations, a summation/opponency location combination, and interaction with the thresholding Rectified Linear Unit (ReLU). Re-analyses of mice’s superior colliculus neuronal activity during a cueing task show CNN-predicted but previously unreported cue-inhibitory, location- summation, and location-opponent cells in addition to the commonly reported cue (attention) excitatory cells. The novel single-unit CNN analysis approach establishes a likely system-wide characterization mediating covert attention and a framework to identify new neuron types and emergent computational mechanisms contributing to perceptual behavior.

**Significance Statement:** Cues predictive of a target location orient covert attention and improve perceptual performance. Studies have focused on how attention influences neural activity, but how cues activate attention and how entire neuronal populations result in behavioral signatures of attention is not understood. Using neuroscience-inspired approaches to characterize properties of 1.8M CNN neurons, we predict new neuron types and emergent mechanisms. A re-analysis of mice’s superior colliculus activity shows new neuron types predicted by the CNN. The findings might explain how organisms, from primates and crows to insects, show behavioral signatures of covert attention using a variety of cue-target-location neuronal properties. The new CNN single-neuron analysis can guide neurophysiological studies to identify potential new neuron types and their computational contributions to perception.

## Introduction

Spatial covert visual attention refers to the process of selecting a location in an image or a scene without moving the eyes and can greatly benefit perceptual performance in detection, discrimination, and search. Both behavioral^1,2^ and electrophysiological studies^3,4^ often operationalize manipulations of covert attention to a location by using cues that co-occur with or point to a likely target location.

Subjects tend to be faster and more accurate when the target is at the cued location (valid cue trials) vs. when it appears at the uncued location (invalid cue trials).

There is a vast literature focused on understanding the influences of covert attention (cueing) using behavioral measures^2,5–7^. Early research focused on a descriptive understanding of attention in cueing tasks based on analogies such as a zoom or a spotlight^8,9^. Subsequent studies in the last three decades have used psychophysical results across stimulus conditions (e.g., varying target contrast^10,11^ and image noise^6,12^) and, by assuming an underlying quantitative model, made inferences about the mechanisms by which attention changes the processing of visual information. These studies have proposed that attention can exclude external noise^12^, result in signal enhancement^13^, internal noise reduction^13^, arise from interactions between a multiplicative attention field and a divisive suppression^10,11,14^, and optimize the weighting of cued and uncued sensory information^15^. Cell electrophysiological studies with spatial cues have been used to show that attention can increase the amplitude^4^, but not the width, of the orientation tuning curve ^4^, reduce inter-neuronal correlations^3^, and change the contrast response functions^16^ of neurons. Recently, studies have incorporated neuro-inspired attention mechanisms into convolutional neural networks^17,18^ (CNNs) to assess the impact on model perceptual accuracy.

These studies have been fundamental in characterizing different signatures of the effects of attention on visual processing, but they remain far from an entire hierarchical pipeline of computational stages that can be mapped to neural circuits and give rise to behavioral cueing effects. The models typically incorporate an attentional mechanism that alters the processing of information at the cued (attended) locations^6,14,17,18^ but omit the computational stages by which brains integrate the response to the cue with the responses to the target, nor do they predict the distribution of cell response properties that one might expect in the brain of biological organisms that manifest behavioral cue effects and covert attention. Even with the progress in multi-neuronal and multi-size recordings, physiological studies probe a small subset of neurons and areas, typically less than 1000 neurons^19,20^. How to map these measurements into computational stages is not straightforward^21,22^.

The purpose of this paper is to assess the emergent neuronal mechanisms in CNNs that show behavioral signatures of covert attention^23^. The CNN does not incorporate any attentional mechanisms into the model, but instead assesses the mechanisms that alter visual processing that result from task optimization (target detection maximization). Previous work^23^ showed that CNNs with no inbuilt attention mechanism, trained from images to optimize target detection, show human-like behavioral signatures of covert attention across various tasks. Such work did not investigate what computational properties of the network along the processing hierarchy give rise to these signatures of covert attention. In this paper, we characterize the entire multi-stage processing of the visual stimulus in these CNNs and understand how the hierarchy of computations across all networks’ units (neurons) of each layer can lead to the signature cueing behaviors shown by many organisms, from humans^1,2^ and monkeys^21^ to archerfish^24^, and bees^25^. Our analysis seeks to answer the following questions: At which stage are the cue, target, and location information integrated? When does the cueing effect arise in the network, and what are the emergent mediating mechanisms? Do the CNN predictions agree with reported neuronal measurements of the existing vast literature^19,26–29?^ And is there evidence for the CNN’s novel predictions about neuronal properties beyond the almost exclusive focus of the field on excitatory influences of attention (cue) on neuronal responses^19,26–29?^

We train a CNN on the Posner cueing task, shown to result in a behavior cueing effect^23^, and study the CNN’s neuronal properties that emerge. The network uses images embedded in luminance noise as inputs and is trained to detect whether a target (tilted line) is present at either location, or distractors with less tilt are presented at both locations. A cue (box around one of the two lines) indicates the likely target location when present. The network has no explicit knowledge about the cue or its predictive probability. We also do not explicitly incorporate any attention mechanism (e.g., gain^18^, prior^15,30^, feature similarity^17^) into the network to act differently at the cued location. We use classic neuroscience-inspired methods (single-neuron Receiver Operating Characteristic^31^ and reverse correlation) to characterize the tuning properties of each CNN unit (referred to as a neuron) to the target, cue, and locations, and the emergent mechanisms by which the cue influences target processing. We compare the properties of 180K neuronal units at the different layers of ten trained CNNs (1.8 M neurons) to the theoretical computational stages of an image-computable Bayesian ideal observer (BIO) that integrates visual information of the cue, the target, and across locations to maximize perceptual performance detecting the target. The BIO has been used as a normative model to assess how biological organisms process visual information^32–36^ and as a benchmark model of visual search^37–41^ and attention^7,15,39,42,43^. Understanding how a large population of network neurons represents task-relevant information relative to the computational stages of BIO might elucidate how probabilistic computations could be carried out in the brain. Finally, to illustrate the value of the neuroscience-inspired CNN analyses, network emergent attention mechanisms are compared to mice’s superior colliculus (SC) neuronal properties during a cueing paradigm by re-analyzing a previously published dataset.^19^

The results section is divided into four parts: 1) Behavioral predictions of the CNN for the cueing task; 2) Statistical characterization of CNN’s neuronal properties using neuroscience-inspired methods; 3) Emergent neuronal mechanisms mediating the cueing effect in CNNs; 4) Relating CNN predictions about neuron types to mice’s superior colliculus cell properties.

## RESULTS

### Behavioral predictions of the CNN for the cueing task

The task required that humans and models detect the presence of a target that appeared with a 50% probability. The target was a tilted line with a tilt angle sampled from a Gaussian distribution (mean 14 degrees (deg; Figure 1a), standard deviation 3.5 deg). Target-present trials contained a target at one location and a distractor at the other. Distractors were lines with orientations sampled from a Gaussian distribution (mean 7 deg, standard deviation 3.5 deg). Target-absent trials contained independently sampled distractors at both locations. A box cue appeared in all images and appeared with the target with 80% probability when the target was present. The cue had an equal probability of being present at either location and was presented simultaneously with the test image (target/distractor). White luminance noise was added to the images (see Methods).

**Figure 1.**
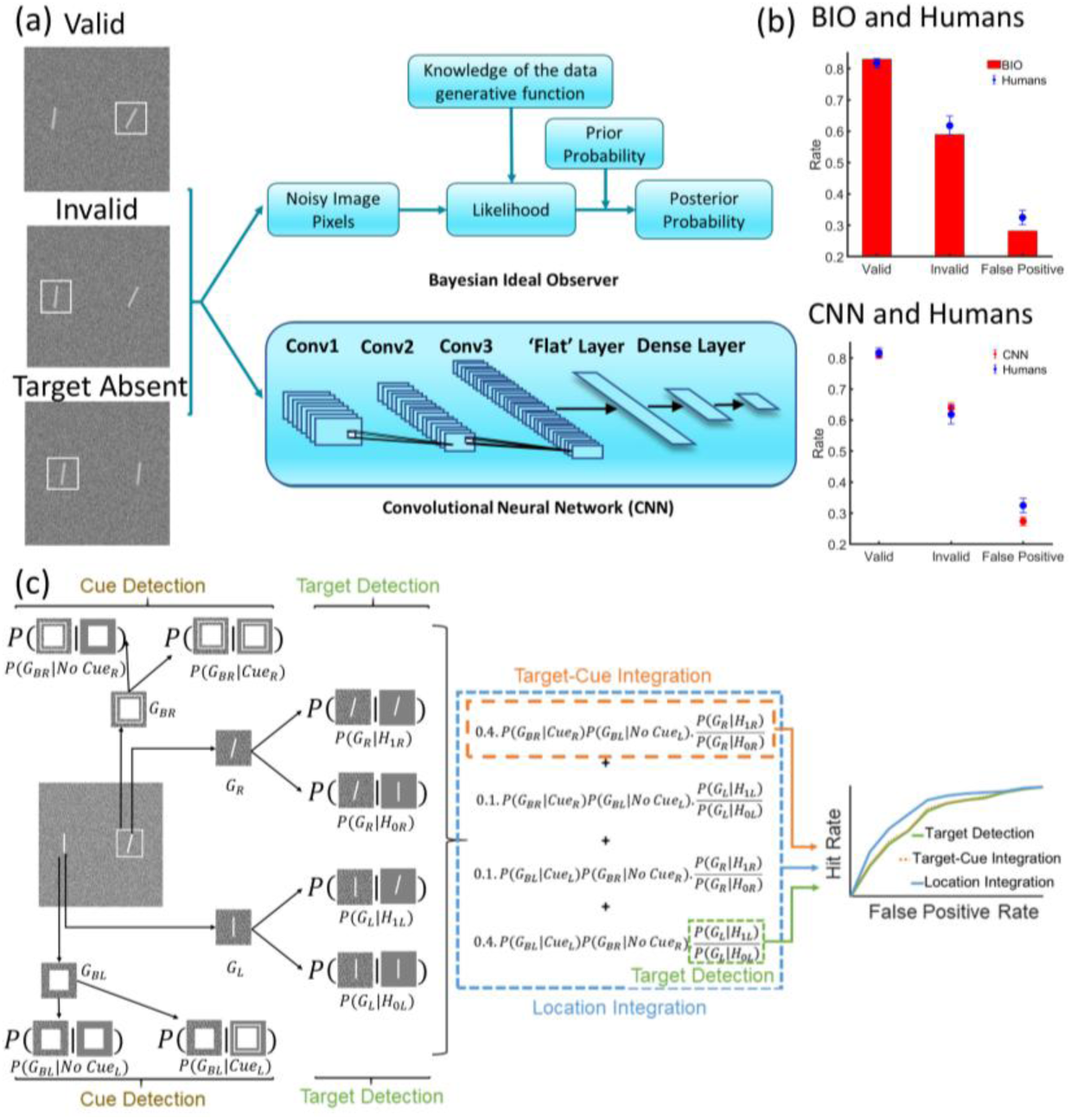
Example stimuli, model flowcharts, and decision accuracies of models and human observers. (a) Left: Example stimuli. Valid trials have the cue (box) appearing with the target (tilted line) and comprise 80% of the target-present trials; invalid trials have the cue appearing opposite to the target and comprise 20% of target-present trials, while target-absent trials comprise 50% of all trials. Target and distractor angles have variability (standard deviation 3.5 deg). The images contain additive independent Gaussian luminance noise. Right: Stimulus processed by an image-computable Bayesian ideal observer (BIO) and a Convolutional Neural Network (CNN). The flattened third convolution layer is referred to as the Flat layer. (b) Performance of eighteen human observers (blue) compared to the BIO (left) and CNN (right)(red) with adjusted RMS noise to match overall human performance. Blue error bars represent the standard error of eighteen human observers, red error bars represent the standard error across ten trained CNNs. (c) Three computational stages of the BIO: the cue/target detection stage (brown/green), where noisy stimulus pixels *G*_*Bi*_ (where subscript B refers to the surrounding background at location *i*) and *G*_*i*_ are compared to the template images of the cue and the target to compute the likelihood of cue (*P*(*G*_*Bi*_ |*Cue*_*i*_) or *P*(*G*_*Bi*_ |*No Cue*_*i*_)) or target presence and absence *P*(*G*_*i*_|*H*_*ji*_), given the noisy stimulus pixels at each location, after which the BIO computes the ratio of the target presence and absence likelihoods, LR= *P*(*G*_*i*_|*H*_1*i*_)/*P*(*G*_*i*_|*H*_0*i*_); the target-cue integration stage (orange), where the likelihoods for the cue and the target are multiplied with the prior probabilities *π*_*i*_ of target presence and absence given cue presence and absence: *π*_*i*_. *P*(*G*_*Bi*_ |*Cue*_*i*_). *P*(*G*_*B*!*i*_|*No Cue*_!*i*_). *P*(*G*_*i*_|*H*_1*i*_)/*P*(*G*_*i*_|*H*_0*i*_), where !i denotes the location other than the i^th^ location; and the location integration stage (blue), where the local target-cue integration stages are summed across both locations. The computations involved in each stage are shown in dotted boxes of corresponding colors, and the associated ROCs detecting the target for each stage (right panel). Note that the orange and green ROC curves overlap while the blue ROC curve lies above the other two ROCs showing that for the BIO the cueing arises after integration across locations. See Methods for more details.

The CNN was a feedforward convolutional neural network with five layers: three convolutional layers with a stride size of 3 with 8, 24, and 32 kernels, respectively, of size 3 by 3, followed by two fully-connected layers with 256 and 2 neurons, respectively (Figure 1a). The flattened third convolution layer is referred to as the flat layer, and the first fully-connected layer is referred to as the dense layer throughout the paper. The networks were trained via backpropagation to detect target presence/absence in noisy images until convergence. The CNNs did not use human trial responses, thus, the cueing does not emerge due to fitting the CNNs to the human data.

The image-computable BIO for the cueing paradigm^15,39^ (Figure 1a-c) is well-known and consists of weighting sensory evidence about the target (the likelihood) by the prior probability of the target appearing with the cue. The model often omits the process of cue detection and assumes detection with no error, but has been recently generalized to incorporate the detection of the cue, which is also presented in additive luminance noise^23^ (see Supplementary Section S.1a for details).

Both the CNN and the BIO attain a higher overall performance than humans when evaluated with the stimuli humans viewed (see Supplementary section S.1b-c), thus, we increased the external white noise of the images separately for each model to minimize the Akaike Information Criterion (AIC, Supplementary section S.1d), with respect to humans. The CNNs, trained to optimize overall accuracy across all trial types, and the BIO, show a cueing effect^15,43^ (Figure 1b) comparable to that of eighteen human observers in the two-location task (see methods), consistent with a recent publication reporting behavioral cueing effects of CNNs^23^. The CNN’s cueing effect is related to learning the statistical relationship between the predictive cue and the presence of the target. For example, a CNN trained on images with a non-predictive cue (50%) does not result in CNN behavioral cueing effects. Similarly, CNN training on an 80% predictive cue and testing on images that contain cues on both locations does not lead to accuracy improvements over images without cues (see Supplementary Section S.1.e.).

### Statistical characterization of CNNs’ neuronal properties using neuroscience-inspired methods

In the next sub-sections, we analyze the neurons’ responses to the target, cue, target-cue integration, integration across locations, and the influence of the cue on target sensitivity. All evaluations consist of neurophysiology-inspired methods to characterize the neuron’s responsiveness by generating samples of noisy images with the presence/absence of the target, cue or both at different locations, collecting response distributions for each neuron in the CNN across separate sets of sample images (target/cue present or absent), generating an ROC curve by varying the decision threshold, and estimating the area under the ROC curve (AUROC)^44^ to quantify the neuron’s sensitivity (Figure 2a). To ensure reproducibility, all evaluations included ten separate training repetitions of the networks, each with testing over 10,000 trials. Luminance noise was independent for each training and testing sample. Figure 2b summarizes all neuronal analysis, the specific conditions to generate the AUROCs, and the terminology adopted in the paper for the critical measures to be presented in the next sections.

**Figure 2.**
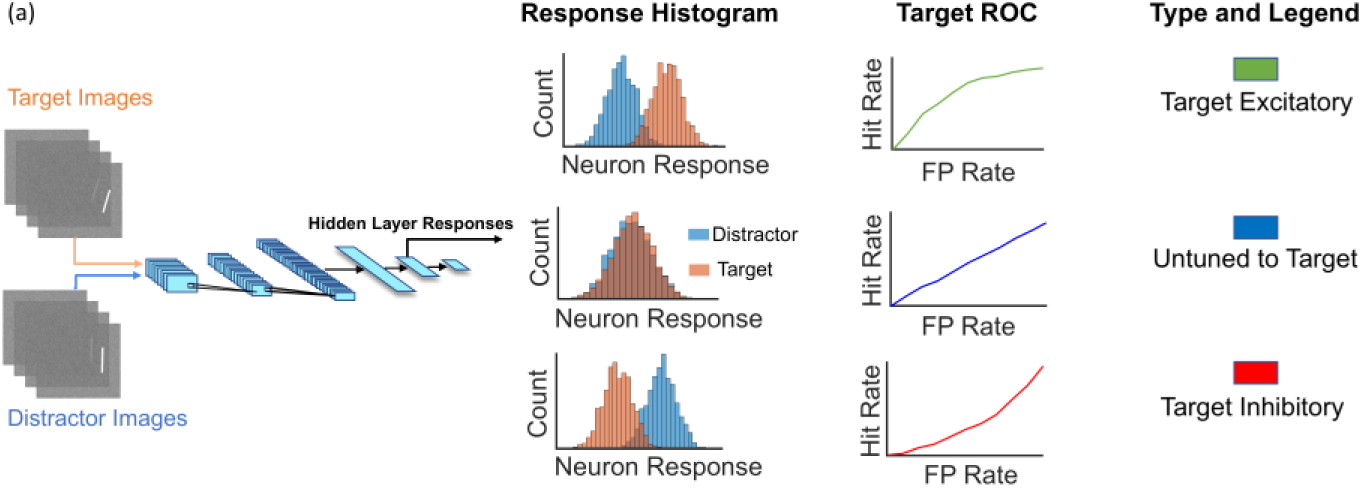
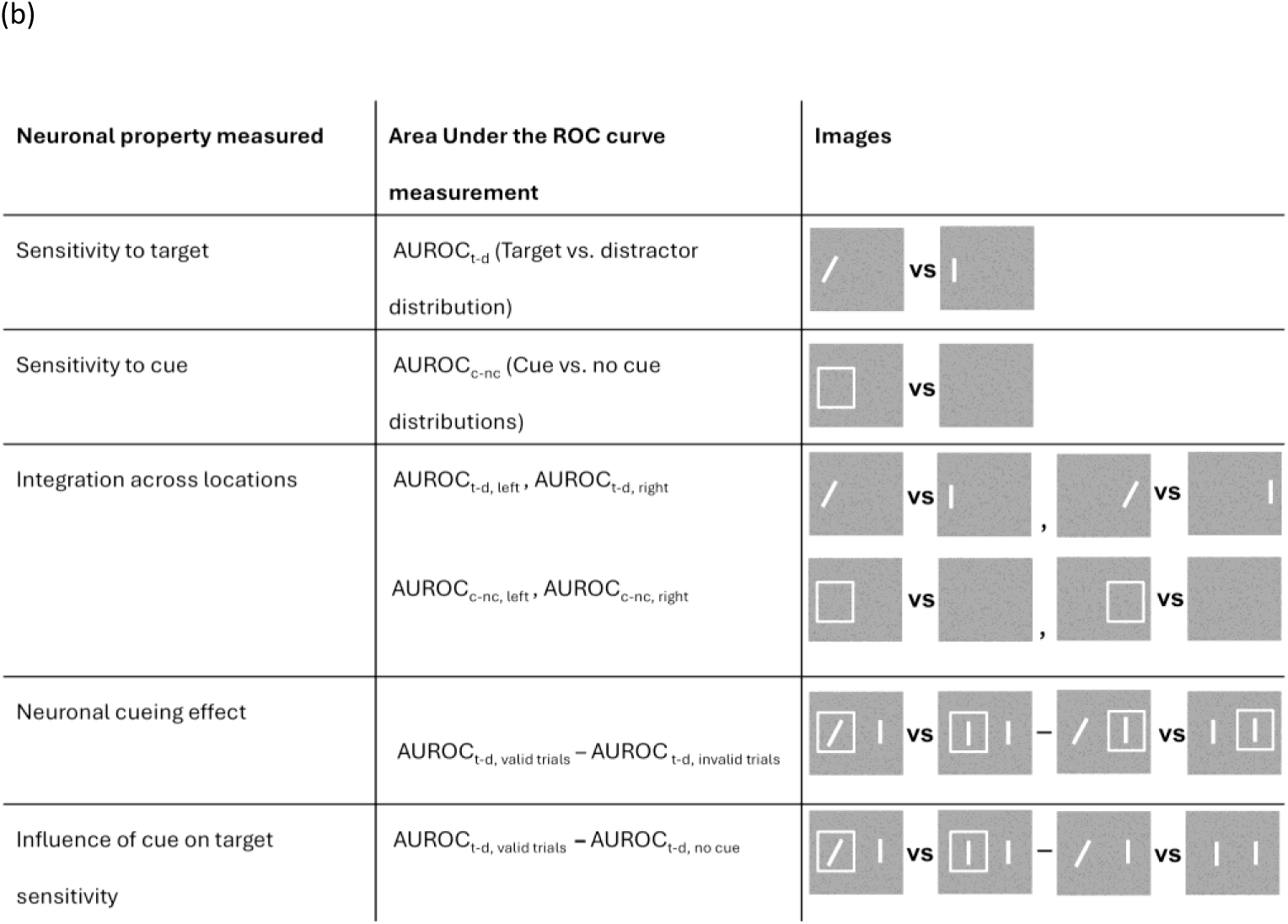
Receiver operating characteristic (ROC) curves, pipeline, and summary. (a) Flow diagram for calculating CNN unit (neuronal) sensitivity to target AUROC using 10K images with noise target present or absent (distractor present) and no cue. Neuronal response distributions for target-present trials and target-absent images. The ROC curve is traced by varying a decision threshold, and the Area Under the ROC (AUROC_t-d_) is estimated as a measure of target sensitivity. Right: Example of neuron types in the CNN, their response histograms to the target and distractor, and ROCs: target excitatory, T+ (AUROC_t-d_ > 0.5), Untuned (AUROC_t-d_ = 0.5); target inhibitory, T- (AUROC_t-d_ < 0.5) (b) A summary of all CNN neuronal properties evaluated, how these are quantified using AUROC measures and comparisons, and example images (right) for different conditions used to generate CNN neuronal response distributions. Images used in the evaluation had additive white noise.

#### Neuronal Target and Cue Sensitivity

We first quantified target sensitivity for each of the 1.8M units across the 10 networks. Figure 2a shows sample histograms of three neuron types encountered in the network in the dense layer. Some neurons responded more strongly when the target was present (target-excitatory, T+, AUROC_t-d_ > 0.5, p < 0.01, two-tailed t-test, with False Discovery Rate, FDR correction for every test). Other neurons consistently diminished their response when the target was present (target-inhibitory, T-, AUROC_t-d_ < 0.5), a third type was insensitive to the target presence (Figure 2a, middle histogram, AUROC_t-d_ = 0.5), and a fourth type was unresponsive altogether. Figure 3a shows how the percentages of T+ and T- neurons increase as the processing progresses through the network layers (see Figure 3e for percentages. Supplementary Figure S2 shows layerwise histograms of AUROC_t-d_). Figure 3b shows how the CNN’s neuronal target sensitivity (absolute difference between AUROC_t-d_ and 0.5) increases with processing across layers (see Supplementary section S.2 for stats).

**Figure 3.**
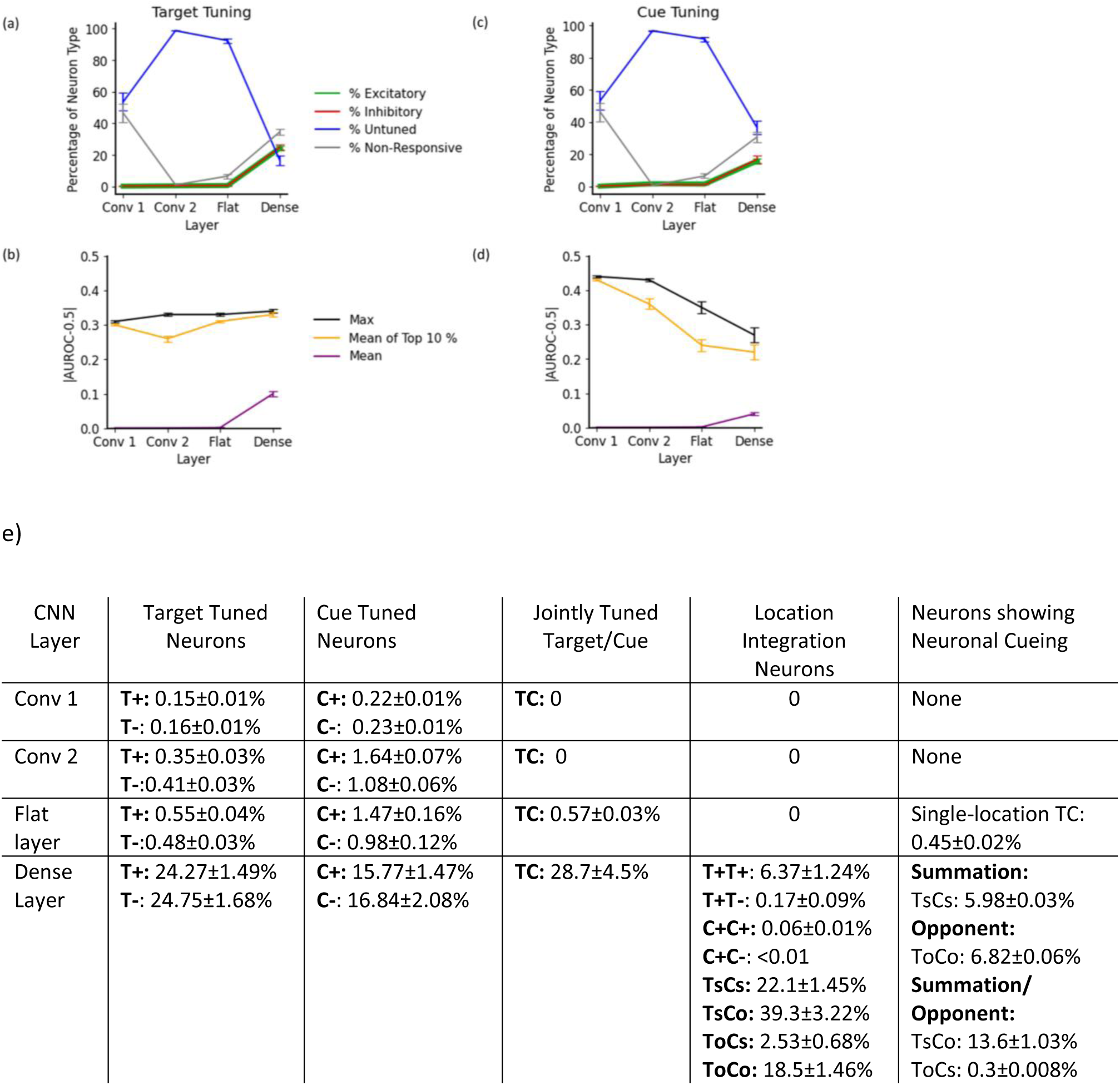
Summary statistics for neuronal tuning to target and cue. First row: Percentages of different neuron types by layer based on (a) target tuning and (c) cue tuning. Second row: Layerwise mean, max, and mean of the top 10% of the absolute difference between target AUROC_t-d_ and 0.5 (b) and cue AUROCc-nc and 0.5 (d) for ten CNN models trained on 80% valid cue. Error bars are standard errors over ten networks. (e) Table showing the exact percentage of all neurons in each CNN layer for different neuronal types (standard error across ten networks): T+, target excitatory; T-, target inhibitory; C+, cue excitatory; C-, cue inhibitory. Neurons with two letters (T+T-) indicate their response property across the two locations; TsCs is a target and cue location-summation unit (T+C+T+C+); ToCo is a location opponent unit (T+C+T-C-), TsCo and ToCs are units with mixed location-summation and opponency (opponent for cue and summation for target or vice versa).

We conducted a similar analysis to quantify the neurons’ cue sensitivities by computing the AUROCc-nc with the cue present or absent in noise (50% probability, see 2^nd^ row Figure 2b). The evaluation images did not contain targets or distractors. We classified each neuron based on the AUROCc-nc as cue- excitatory (C+, AUROCc-nc > 0.5), cue-inhibitory (C-, AUROCc-nc < 0.5), cue-untuned (AUROC c-nc = 0.5), and unresponsive (Figure 3c). Figure 3c shows that the percentage of neurons tuned to the cue increased as processing progressed through the network (Figure 3e, second column, see Supplementary Figure S3 for layerwise histograms of cue AUROC). However, unlike neuronal target sensitivities, the neurons’ max and mean top 10% cue sensitivities (|AUROC- 0.5|, Figure 3d) diminished greatly from Conv1 to the dense layer (p < 0.05 for all comparisons except for max sensitivity between Conv1 and Conv2). This result shows that the network’s representations of the target and cue are different along the processing hierarchy.

#### Target-Cue Integration Neurons

A fundamental step to give rise to a cueing effect must involve units that integrate cue information with sensory information related to the target. We assessed how neurons at different layers are jointly tuned to target and cue. Figures 4a-d show layerwise neuronal AUROCs for the target plotted against those for the cue. Neurons falling along the two AUROC = 0.5 lines (vertical and horizontal perpendicular lines) correspond to neurons separately tuned to only the target or the cue. Neurons that fall off the two perpendicular lines correspond to neurons that are jointly tuned and integrate target and cue information (TC neurons which depending on whether the AUROC is greater or smaller than 0.5 for the target and cue can be: T+C+, excitatory to both target and cue, in the upper right quadrant; T-C+ target-inhibitory and cue-excitatory, in the bottom right quadrant; and so on T+C- and T-C-). Only the neurons in the deeper layers show an integration of the cue and target detection (flat layer onwards, Figure 4c-d).

**Figure 4.**
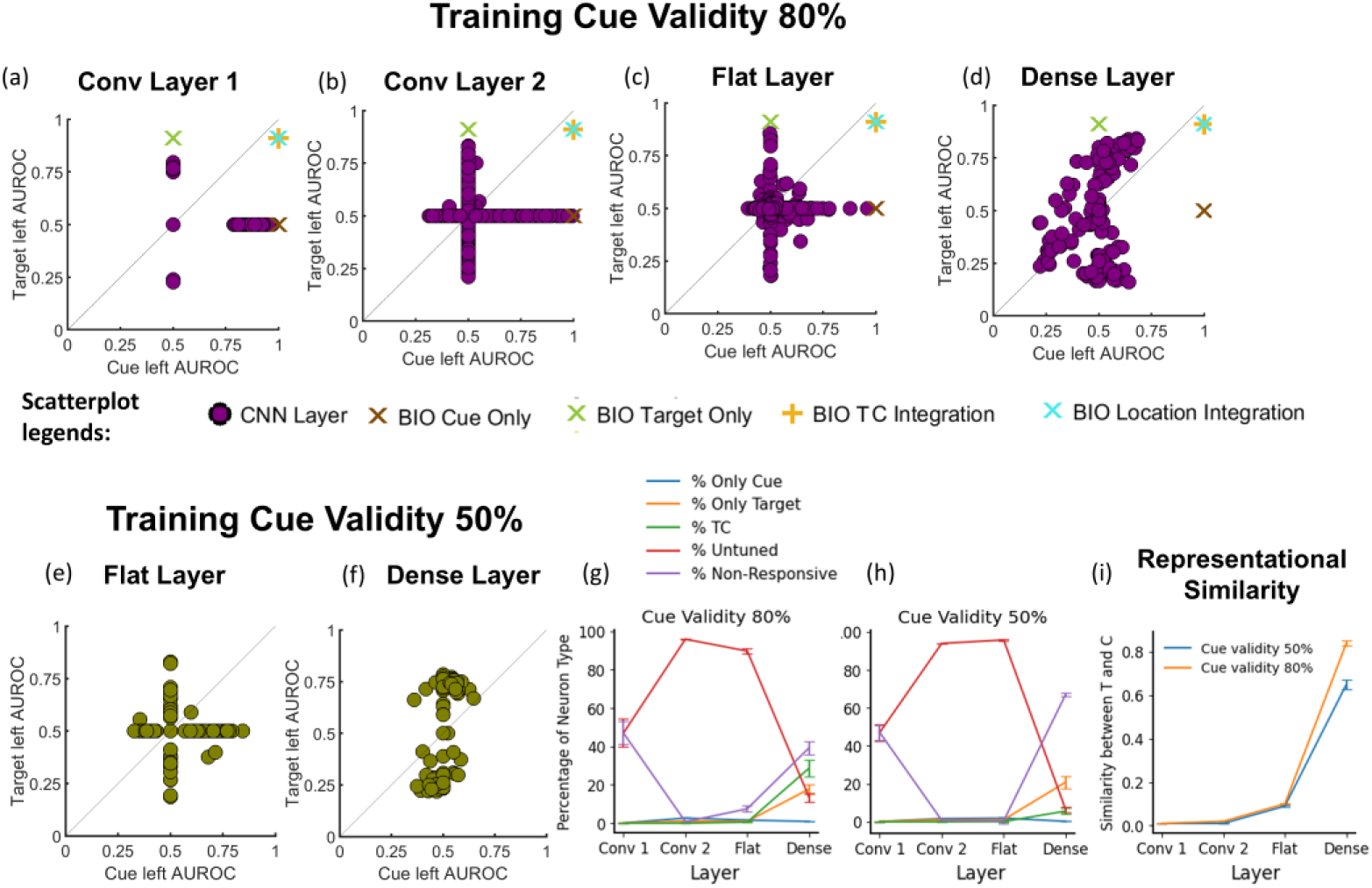
Target-cue integration at the same location. Top row: Scatterplots of target AUROCs vs. cue AUROCs at the left location for neurons in the convolution layers (a-b), flat layer (c), and dense layer (d) of CNNs with 80% training cue validity. The ROCs obtained from the corresponding stage of the BIO are also marked on the scatter plot. (e-f) Scatterplots of target AUROCs vs. cue AUROCs at the left location for neurons in the flat layer (e), and dense layer (f) of CNNs with 50% training cue validity. All results are for the target and cue on the left location, but similar results are obtained for the right location. (g)-(h) Relative percentages of the various neuron types for 80% (g) and 50% (h) valid cues. (i) Layerwise representational similarity between population responses to the target and the cue for 80% and 50% valid cues.

As a control, an analysis for the flat and dense layers of a network trained on a 50% predictive cue (Figure 4e-f; see Supplementary Figure S4 for convolution layers) shows similar separate subpopulations of target- and cue-tuned neurons in the flat layer (Figure 4e) but a significantly lower number of TC neurons in the dense layer compared to the CNN trained with 80% predictive cue (Figure 4f vs. 4d; Figures 4g-h for a summary of percentage of neuron types for 50% vs. 80% cue validity). To complement the single-neuron level analyses, we investigated population-level representational similarity analysis (RSA)^45^ of the layerwise population activity to target-only and cue-only images (averaged across trials; see methods) for networks trained with cue validities of 80% and 50%. We find that the target-cue population activity similarity is the lowest and does not vary with cue validity in the early convolution layers. In contrast, consistent with our single-neuron analyses, it is highest and varies with cue validity in the dense layer (Figure 4i).

#### Integration across spatial locations

Detecting a target across two possible locations in a cueing task requires the organism to integrate information across the two locations. We evaluated the target and cue sensitivity (AUROC) of each neuron for each of the two locations (see Figure 2b). Neurons falling along the cardinal axes are tuned to one location, while neurons off the cardinal axes are tuned to both locations. Conv1, Conv2 (Supplementary Figure S5), and flat layers (Figure 5a) have target and cue neurons tuned to single locations (no integration across locations). In contrast, the dense layer (Figure 5b) showed neurons integrating across both locations (AUROC ≠ 0.5 for both locations). We found distinct types of TC neurons that integrated visual information across locations: location summation neurons (AUROC > 0.5 for both locations, upper right quadrants in Figure 5b, labelled TsCs units), location opponent (excitatory for one location and inhibitory for the other, ToCo units), and combination of location summation and opponent cells (TsCo and ToCs, see Figure 3e for exact % of each type).

**Figure 5.**
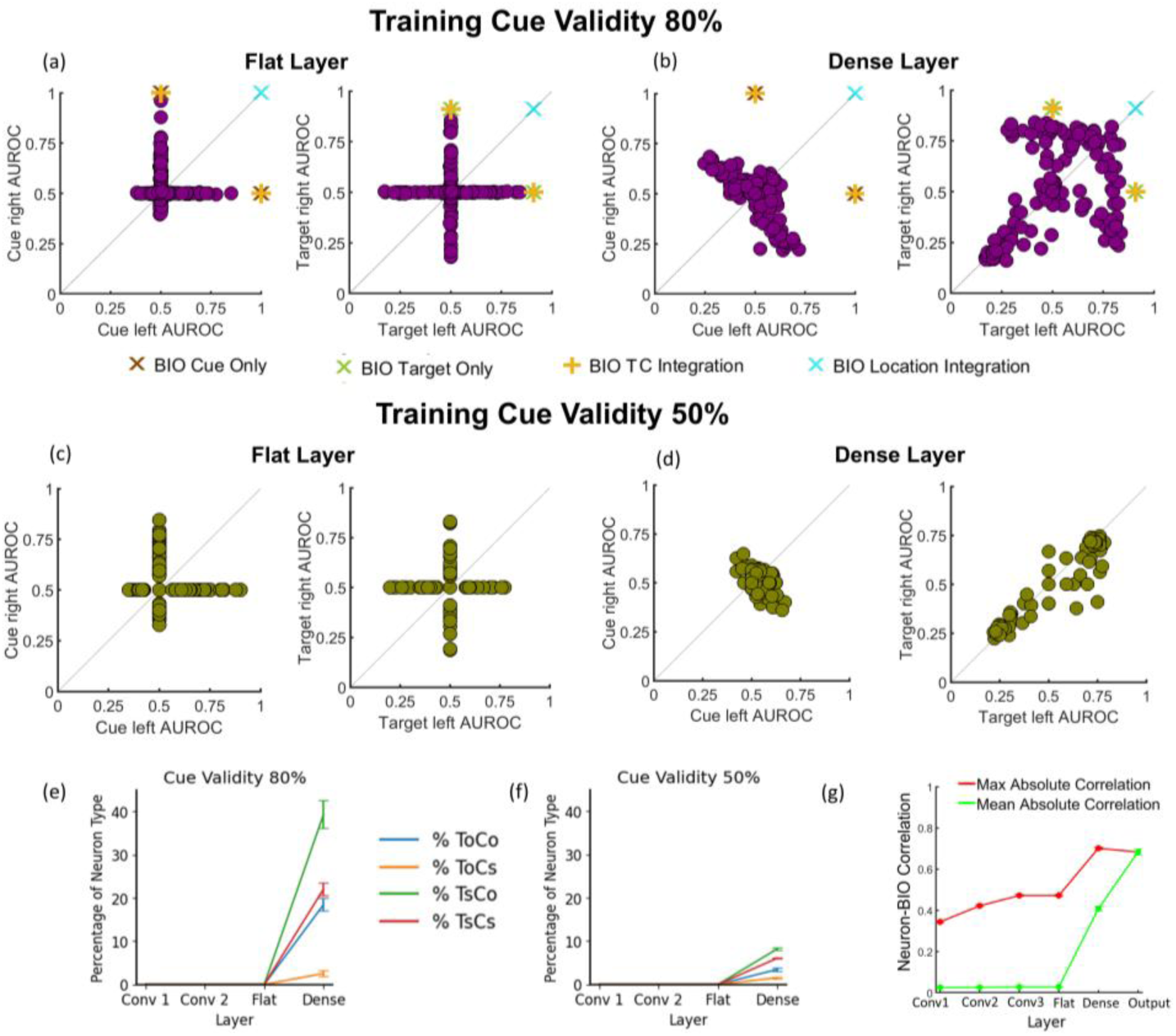
Integration across locations. (a-b): Scatterplots of neuronal AUROCs for cue and target detection at the right and left locations for neurons in the flat and dense layers for the CNN trained with 80% cue validity. The AUROCs obtained from the corresponding stage of the BIO are also marked on the scatter plot. (c-d): Scatterplots for neuronal AUROCs for cue and target detection at the right and left locations for neurons in the flat and dense layer for CNN trained with 50% cue validity. Third row: Layerwise percentage of neurons for various location integration types (see Figure 3e caption for definition of neuron types) for 80% (e) and 50% valid (f) cue. (g) Layerwise mean and max neuron-BIO decision variable correlations (absolute value) averaged across ten CNNs.

As a control, analysis of a network trained on a 50% predictive cue (Figure 5c) shows fewer TC neurons in the dense layer (Figure 5b vs. 5d, 5e vs. 5f), with the TsCo units resulting in the greatest decrease (see Supplementary Figure S4-5 for changes in the convolution layers).

#### CNN and optimal computations

Figures 4a-d and 5a-b plot the AUROC for a BIO’s decision variables at different stages of its computational hierarchy (Figure 1c): cue detection, target detection, target-cue integration, and location integration. There is a correspondence between the types of neurons that emerge in the network and the stages of processing in the BIO (see Supplementary Figure S23 for a CNN visualization and comparison to BIO stages). The neurons in the early layers map to the separate cue and target detection stages of the BIO (Figure 4a-b). Neurons in the dense layer (Figures 4d, 5b) show properties similar to the BIO decision variable at the target-cue integration stage and the final BIO location integration stage. The neurons at the flat layer (Figures 4c, 5a) show an intermediate stage with more neurons involved in separate cue and target detection and a small proportion of TC neurons. There are some important differences. The BIO shows very defined properties with higher AUROCs and abrupt transitions to cue/target integration, while the CNN shows a larger range of AUROCs across its populations and a more nuanced transition toward target-cue integration across layers. To obtain a quantitative measure of the correspondence between neurons’ responses and optimal computations, we correlated the BIO decision variable (log-likelihood, Supplement Section S.22) and each neuron’s response across trials. Figure 5g shows the layerwise increases in the mean and maximum (absolute value) neuron-BIO correlations (see Supplementary Figure S24 for a layerwise visualization).

### Emergent neuronal mechanisms mediating the cueing effect in CNNs

Next, we seek to understand the emergent mechanisms in the network responsible for the CNN’s behavioral cueing effect. We first identified neurons that showed a neuronal cueing effect defined as a higher AUROC_t-d_ discriminating the target vs. distractor in valid cue trials relative to invalid cue trials (see neuronal cueing effect, Figure 2b). We found an increasing number of neurons showing cueing effects as the processing progressed across the layers (Figure 6a; also higher influence of the cue on the mean neuronal responses and AUROC_t-d_ across layers - see Supplementary Section S.6 for layerwise distributions of neuronal change in target sensitivity, mean responses, and standard deviations due to the cue). Of the 11% of the flat-layer neurons showing neuronal cueing, all were single-location TC neurons. Of the 56% of the dense layer neurons that show neuronal cueing, we identified three major types (see Supplementary Section S.7 for ∼7% of residual types): 24.29% location summation, 22.61% location-opponent, and 46.33% location summation-opponency combination (e.g., target summation and cue opponency).

**Figure 6.**
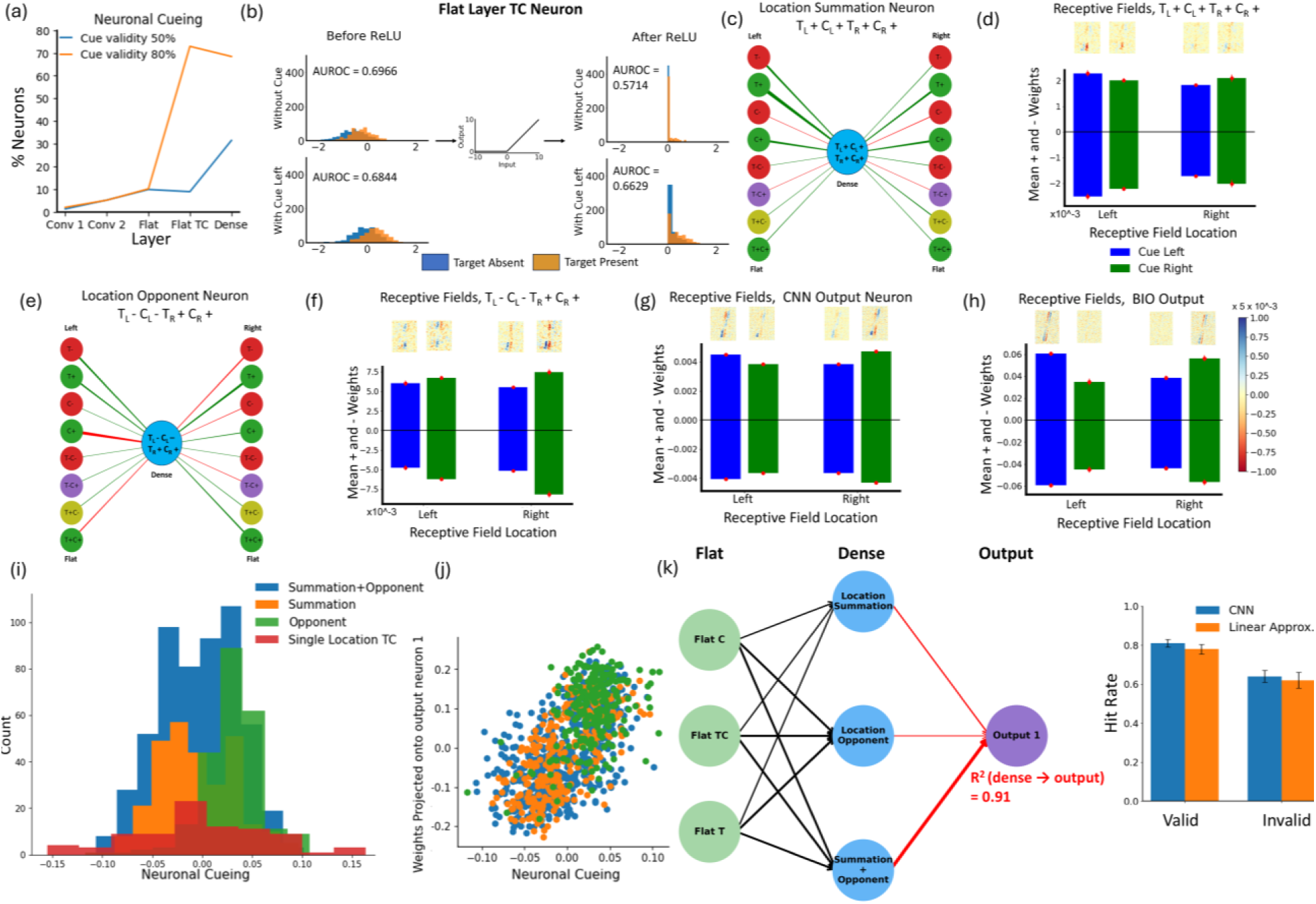
Neuronal Cueing across layers and cueing mechanisms. (a) Layerwise neuronal cueing for 80% (orange) and 50% (blue) cue validity. (b) Response distributions before and after ReLU of an example flat-layer TC neuron when the target is present at the left location(orange) and absent (blue). The top row shows responses when the cue is absent, the bottom row shows responses when the cue is present at the left location. (c) Weights projected by various flat layer neurons from the two locations into a sample location-summation neuron in the dense layer. The color of each connecting line (green/red) denotes whether the sum of weights from that neuron type to the dense neuron is positive/negative. The thickness of each connecting line corresponds to the magnitude of the sum of weights between that neuron type and the shown dense neuron. (d) Linear receptive fields (RF) obtained via linear regression predicting neuronal response for the neuron shown in (c) with cue on the left (above blue columns) and the right (above green columns). Bar plots depict mean positive and negative RF weights for each location and condition. (e): Same as (c) but for a location opponent neuron. (f) Linear RFs predicting neuronal response for the neuron shown in (e) with cue on the left (above blue columns) and the right (above green columns). Note that the RF for the left location of the location opponent neuron (TL-) is excitatory to the vertical distractor and inhibitory to the tilted target. (g) Linear RFs and associated weights for the network’s output neuron representing target presence. Error bars indicate standard errors over ten reruns. (h) Linear RFs and associated weights for BIO model (log of weighted likelihood ratio). (i) Histograms of neuronal cueing by mechanism types. (j) Scatterplots depicting the relationship between the dense layer units’ neuronal cueing and their weights projected onto the output layer for each mechanism type in the dense layer. The different colors correspond to different mechanism types consistent with the legend of graph (i). (k) Left: Simplified circuit diagram depicting how the three dense-layer attention mechanisms (location summation, opponency, and summation/opponency) receive input from the three flat neuron types, and feed into the CNN output neuron.

Arrow widths between dense and output layers depict the mean absolute regression weight for each dense neuron mechanism type for predicting the output responses. Arrow widths between the flat and dense layer depict the mean absolute regression weights for each flat neuron type for predicting the average response of each dense mechanism type (see Supplementary section S.25 for details). Right: valid and invalid hit rates of the CNNs (blue) and linear regression (orange).

Below, we describe three of the main emergent mechanisms in detail (due to space constraints, the fourth mechanism, summation + opponency, is discussed in Supplementary Section S.8)

#### Single-Location TC Neurons with ReLU interaction

The first mechanism involves TC neurons in the flat layer tuned to a single location. The single location- TC neurons show cueing effects before the integration across locations. What mediates the change in target sensitivity (AUROC valid cue) in the flat layer? The effect arises from the interaction of the influence of the valid cue on the neurons’ mean response with the non-linear ReLU thresholding. Figure 6b illustrates a sample neuron’s response distributions to the target and distractor before and after the ReLU. The top left graph shows a flat-layer neuron with pre-ReLU AUROC_t-d_ of 0.697±0.007 without the cue. The ReLU zeroes all negative values, effectively cutting off part of the tails of the target and distractor distributions (but more of the lower end of the distractor distribution). This thresholding reduces the AUROC of the shown neuron when the cue is absent. Figure 6d (bottom left) shows the same neuron’s response distributions but with the valid cue, which has a pre-ReLU AUROC of 0.684±0.006. The valid cue increases the neuron’s mean response (and variance) to the target and distractor, and a larger part of both response distributions shifts to the right above zero. Thus, less from both distributions are truncated, resulting in a lower decrease in AUROC_t-d_ for valid cue (Pre-ReLU: AUROC_t-d_ valid cue = 0.684±0.006, AUROC_t-d_ no cue = 0.697±0.007, p = 0.0753; Post-ReLU: AUROC_t-d_ valid cue = 0.663±0.005, AUROC_t-d_ no cue = 0.571±0.004, p < 10^-5^; see Supplementary Figure S7f for dense layer, and Supplementary section S.8 for estimated receptive fields for the single-location TC neuron).

#### Location Summation TC Neurons

The second mechanism involves neurons with summation across both locations for the target or cue (TL+CL+TR+CR+). We analyzed the response properties of the input neurons and their weights to characterize such neurons’ connectivity, represented in Figure 6c. Reverse correlation estimation of the linear kernels (see methods) at the cued and uncued location shows that the mechanism’s location- integrated output response is driven more strongly by the visual information at the cued location (mean positive and negative weights of the estimated RF) than the uncued location (Figure 6d; p < 0.01 for each location with and without the cue), a mechanism similar to the CNN’s output neurons (Figure 6g) and BIO’s location integration stage (Figure 6h).

#### Location Opponent TC Neurons

The third mechanism is the location-opponent TC units (see Figure 3e). Figure 6e shows an example of one type of location-opponent units (TL-CL-TR+CR+) and the input neurons from the flat layer. Reverse correlation of the linear kernels shows how the opponent unit increased the weighting of the target- excitatory RF at the cued location and also increased the weighting for the target-inhibitory RF at the location opposite to the cue (Figure 6f), a distinct computation from any stage in the BIO.

#### Mechanism Contributions to CNN Cueing Effects

Figure 6i shows the distribution of neuronal cueing effects for each of the four neuron categories. We found that dense layer units with higher neuronal cueing magnitude were weighted more heavily as inputs to the target presence CNN output neuron (r = 0.635; p < 10^-50^; Figure 6j, see Supplementary Section S.7 for mechanism-specific correlations). To conceptualize how the distinct CNN emergent mechanisms contribute to the CNN’s overall cueing effect, Figure 6k (left) shows a simplified schematic (see Supplementary Section S.25 for details). The first stage shows how the dense layer mechanisms receive inputs from flat layer C, T, and TC neurons. A second stage shows how different dense layer mechanisms combine to result in the CNN’s output response, based on a linear approximation model that captures 91% of the variance of the CNN’s output unit response and generates similar cueing effects as the CNN (Figure 6k, right, with higher weights given to the summation + opponent units followed by the location-summation and then the location-opponent units)).

#### Generality of results across CNN architectures and tasks

Supplementary results (Sections S.5, S.10-12, S.14-16, S.20; Figures S6, S11-S14, S16-17, S22) show that even though some of the percentages of different neuron types vary across conditions, the main results (hierarchy of computations across the network, presence of cue inhibitory neurons, location opponent, summation, and single-location ReLU interaction neurons) generalize to many variations of the task and network properties such as a task with mixed-in neutral cues (half the images from the 80% predictive cue experiment intermixed with 25% of images with no cues and 25% of images with cues on both locations), network dense layer size, number of layers, kernel size, length of training, external noise levels, some levels of sparsity (through L1 regularization) and central cues^1,2^.

Increasing the cue validity from 80% to 100% increased the number of cue-tuned neurons and cue sensitivity (S.9; Figure S10). Similar neuron types emerge in a network pretrained on ImageNet (VGG-16) with the dense and output layers retrained on our task (Supplementary Figures S18-S19), but TC integration emerges earlier in that network. Finally, changing the stride of the convolution kernels (from 3 to 5) resulted in poorer convergence and spatial asymmetries in the cue sensitivity (Supplementary Figure S15).

### Relating CNN predictions about neuron types to mice’s superior colliculus cell properties

In this section, we illustrate how the neuroscience-inspired methods of analyzing emergent neuronal units of a CNN and attention mechanisms can help further understand the response properties of actual neurons during a cueing task. We focused on assessing whether, aside from the typically reported excitatory effects of a predictive cue (attention) on firing rate^19,21,46–48^, there is evidence for some of the neuronal properties as predicted by our CNN theoretical results, which are not typically reported: cue- inhibitory neurons, location-summation, or opponency neurons.

We re-analyzed the response properties of 109 neurons from the superior colliculus (SC) of four mice executing an adapted cueing task. Head-fixed mice were trained to detect the orientation change (50% probability) of a Gabor patch in one of two locations (left or right visual fields). In one condition, a 100% valid pre-cue indicated the possible target location (orientation change). In a control condition, there was no pre-cue. The cue and location vary across locations in blocks. More details about the task can be found in the methods section and the previous publication^19^. Figure 7a lists all the specific AUROC comparisons to assess individual neuron sensitivity to target, cue, and across spatial locations (left and right visual fields for ease of comparison with the CNN; see Figure S20 for ipsilateral vs. contralateral visual fields to recorded cells). We compared the SC results (Figure 7b-d) to theoretical CNN predictions (Figure 7e-g) for ten models trained on the stimuli and task that the mice were trained on (see methods). Figure 7b shows that 30.42% of SC neurons are jointly tuned to the target and cue as compared to 79.28% (out of 127 responsive units) for the CNN’s dense layer (Figure 7e). In addition, both the CNN (70.66%) and SC (27.59%) neurons showed cells tuned to both locations (Figure 7c, f). We further classified the neurons in terms of whether they were excitatory or inhibitory, tuned to a single location or incorporated a location summation mechanism (Ts or Cs) or opponency for the cue or target (see Supplementary Figure S21 for a breakdown of all neuron types found in the SC and CNN). We found SC cells with the typical target- and cue-excitatory responses (T+: 50.0%; C+: 32.9%) but also SC cells with the dense layer CNN-predicted inhibitory response (T-: 39.5%; C-: 30.3% for SC), location- opponency (cue or target, SC: 15.8%), location-summation cells (SC: 11.8%), and SC single-location TC cells (SC: 11.84%; see Supplementary Fig. S21 for complete table). The SC cells also showed single-location T-only (SC: 31.6%), and C-only cells (SC: 19.7%) that are present in the flat layer of the CNN (see Supplementary Fig. S21). Thus, the SC cells showed cell types present in the flat and dense layers of the CNN.

**Figure 7.**
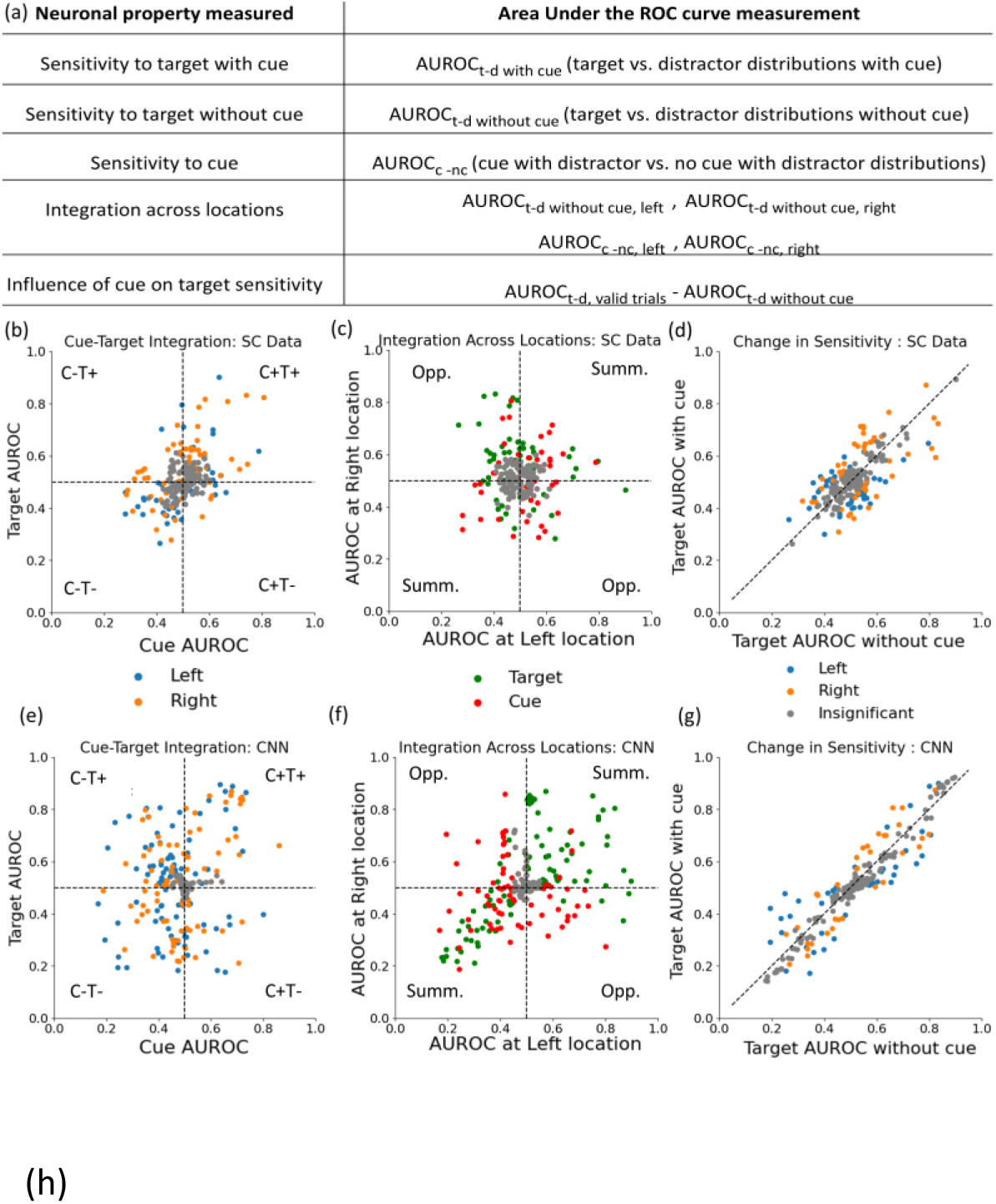
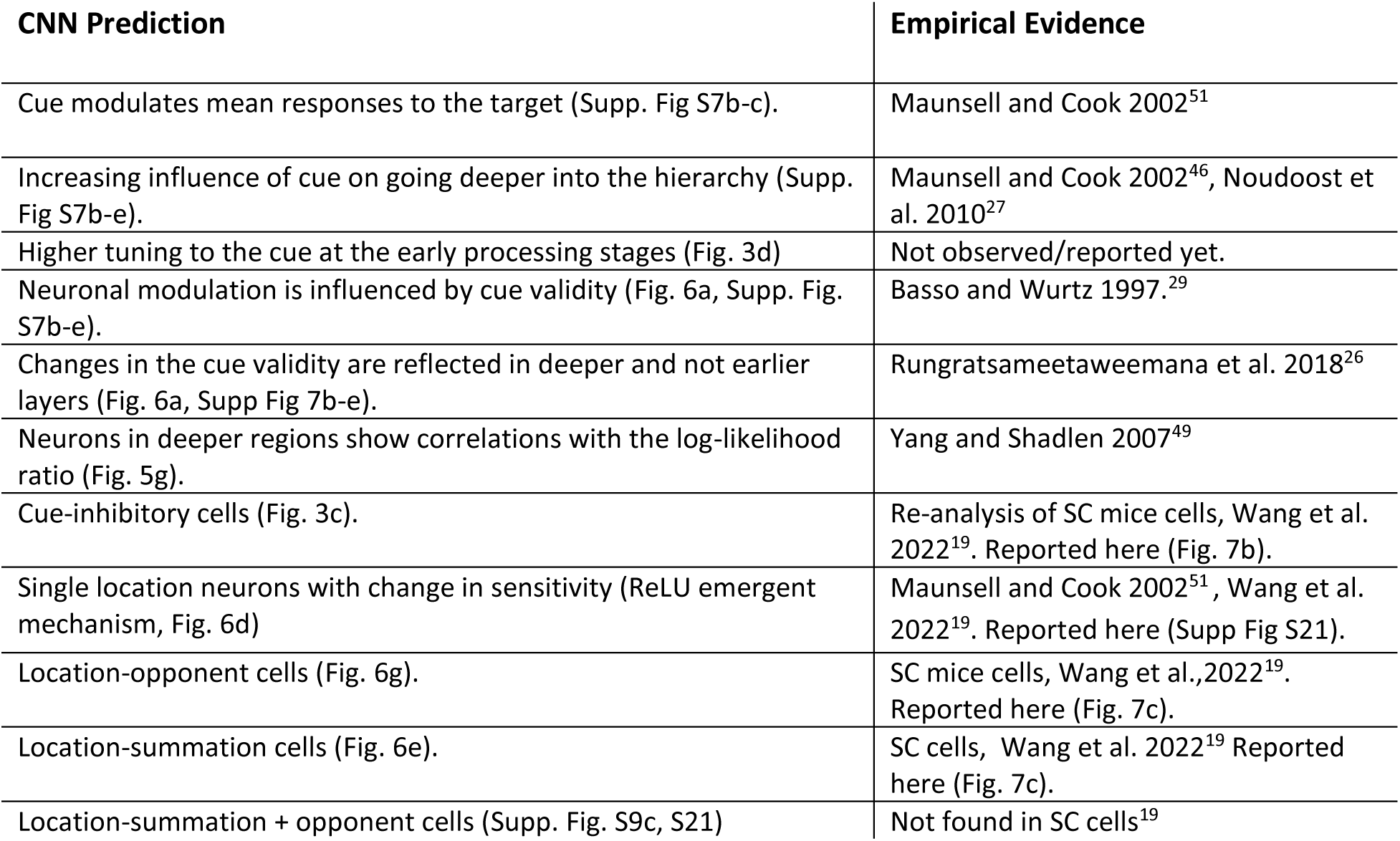
SC neuronal data and model predictions. (a) A glossary of the AUROCs used in the re-analyses of the SC cells. Note that cue sensitivity (AUROCc-nc) is estimated differently here, constrained by the neurophysiological data available, in comparison to the approach used in Figure 2b. (b) Cue and target sensitivity of 109 mice SC neurons during an orientation change detection task with a 100% valid cue and another condition with no cue (Wang et al., 2022). Gray points indicate neurons with statistically non-significant AUROCs for both cue and target. Quadrants are labeled with the neuron types in each quadrant. c) Cue and target sensitivity for each location for SC neurons; Gray points indicate neurons with statistically non-significant AUROCs for both locations. Quadrants are labeled with the neuron types in each quadrant. d) Target sensitivity (AUROC_t-d_) with and without cue for SC cells; e) Cue and target sensitivity for CNN dense layer units trained on the same 100% valid pre-cue task of the mice. f) Cue and target sensitivity for each location for CNN dense layer units. g) Target sensitivity (AUROC_t-d_) with and without cue for CNN dense layer units. (h) A summary of predictions made by the CNN and neuroscience evidence from previous studies.

However, there were some important differences between CNN predictions and SC cells. CNNs showed higher target and cue sensitivities than the SC cells (Figure 7b-c vs. 7e-f). The CNNs had a larger percentage of cue-inhibitory, location-summation cells, and target-inhibitory cells than the recorded sample of SC neurons, which were more excitatory. The CNN also showed ∼10% of cells with a combination of summation and opponency (target-summation, cue-opponency) while the SC showed none. Given this finding, we assessed whether a CNN using only location-opponent and location- summation units (without summation + opponent units) would give rise to cueing effects. A linear regression trained on just location-summation and location-opponent neurons to predict the CNN output neuron responses explained 86% of the variability and showed comparable cueing effects to the original CNN (see Supplementary Section S.26).

In addition, some SC cells showed asymmetries (tuned to one location for the target and both locations for the cue, T1C1+C2-), which were less frequent in the CNNs.

Finally, we calculated the percentage of each SC cell type that showed a significant influence of the cue on target sensitivity (see Figure 7a for definition). We found that 55.6% of the SC single-location TC neurons had a significant change in target sensitivity with the presence of the cue (Figure 7d). Critically, the newly identified location-opponent and summation SC neurons also showed a significant influence of the cue on target sensitivity (33.3% and 44.4%, respectively). These were comparable to the percentage of CNN dense layer neurons (Figure 7g) showing target sensitivity change with the cue (65.2% of single-location TC, 49.7% of location-summation neurons). However, only 12.9% of location- opponent CNN units showed sensitivity change with the cue (vs. SC: 33.3%). Furthermore, averaged across all units, both SC and CNN (dense layer) neurons do not show any target sensitivity change with the cue, consistent with that reported by Wang et al., 2022 for the mice.

## DISCUSSION

For over forty years, researchers have studied visual attention by characterizing the influences of cues on target detection. Although there are numerous theories explaining the cue facilitatory effects in terms of psychological concepts^1,2,50,51^, model-based mechanisms (signal enhancement^6,52^, noise exclusion^12^, gain change of various types^16,53,54^, Bayesian decision weighting by priors^7,15,30,42,43,55,56^, probabilistic population codes^57^, divisive normalization^14,58–60^, neural networks^17^), or neuronal response properties (correlations^3^, contrast response function^16^, tuning curve amplitude^4^), there are no endeavors that start with the image as input, and characterize a whole system-wide visual processing and computations of the target, cue, integration across locations, and how these lead to behavioral performance and cueing effects. Here, we used a five-layer CNN whose representations can be more easily related to actual neural systems than other modeling approaches^30^. We trained the networks to optimize target detection with predictive cues. A critical aspect of our approach is that we did not incorporate into the architecture any specialized attention mechanism^17,60^ that operates at the cued location, nor any explicit resource limitation^61^ shared across locations. The CNN has been shown to result in emergent behavioral cueing effects similar to those of human observers^23^. We used single-cell neuroscience-inspired methods to characterize the tuning properties of every unit in the CNN and understand the hierarchy of processing and emergent neuronal mechanisms that mediate the cueing effects. Below, we discuss our main findings, their relationship to existing findings in neurophysiology (see Figure 7h for a summary) and previous computational approaches, new predictions from the CNN endeavor, and their impact on the field.

### Hierarchical Processing of Cues, Targets, and Locations

Early layers in the CNN are retinotopic, tuned to individual locations, and do not integrate target and cue information. As we move deeper into the network, the integration of cue and target information at each location emerges first (flat layer), followed by the integration of information across the two locations (dense layer). There is an increasing effect of the cue on the CNN’s neural population responses to the target and sensitivity as processing progresses through the layers of the network, consistent with that reported in neurophysiological studies^28,47^. The convolution layers show a small effect, akin to that reported in V1 primary visual cortex single-cell measurements.^4,27^ The dense layer shows strong modulation by the cues more similar to the superior colliculus (SC)^27,62^ and lateral intraparietal areas (LIP)^27^. The flat layer neurons show intermediate effects of the cue, perhaps more similar to those reported in area V4^27^. The increased influence of the cue on the responses to the target occurs even though the sensitivity to the cue itself diminishes with processing across the network layers (Figure 3d). Cue-induced influences on target responses appeared in later CNN layers and are causally related to the cue validity (Figure 6a), consistent with EEG results^26^.

We also show a correspondence between neuronal properties of the CNN and the stages of the BIO (Figure 1c and Supplementary Figure S23) and a higher correlation with the BIO decision variable as processing progresses through the network (Figure 5g), mirroring correlates of BIO log-likelihood ratio in the lateral intraparietal (LIP) region^49^. In terms of the CNN’s emergent mechanisms, the single-location TC neurons correspond more closely to findings of cells in V4^51^ showing cue-related changes in target sensitivity prior to integration of information across spatial locations. There is no previous neurophysiological evidence corresponding to the other three emergent mechanisms (location summation, opponency, and their combination). These and other novel CNN neuronal property predictions are discussed in the next section.

### Novel CNN Predictions about Neuron-Types and Emergent Neural Mechanisms Mediating Covert Attention

The long tradition of neurophysiology of attention has focused mostly on the excitatory influence of attention (cue) on neuronal activity ^19,28,62,63^. Aside from cue-excitatory neurons, our CNN analyses identified different emergent neuron subtypes not discussed in the single-cell electrophysiology literature.^19,28,62,63^. A re-analysis of mouse SC neurons during a cueing task identified, in addition to the common cue/target excitatory neurons, subpopulations of cue and target inhibitory neurons as predicted by the CNN (Figure 7). Besides the more traditional single-location target/cue neurons^46^, the SC also had cells paralleling the CNN location-summation and opponency units. Thus, the SC showed cells present in the flat and dense layers of the CNN. Importantly, a sizable percentage (33-44%) of the newly-reported summation and opponency SC units showed a significant change in target sensitivity with the cue.

In contrast, the opponent/summation combination neurons predicted by the CNN were not found among the re-analyzed SC neurons, but analyses (Supplement section S.26) showed that the CNN’s output and cueing effects could be approximated even if only using CNN location-opposition and summation mechanisms.

There were also some discrepancies between the SC neurons and the CNN in the percentages of each neuron type (e.g., location-summation units, excitatory vs. inhibitory). In addition, CNN AUROCs were higher than SC neurons, which is not unexpected since the CNN (but not mice SC neurons) is optimized for a single task. Despite these differences, the CNN approach highlighted cell types often overlooked in the neurophysiology of attention literature and explains how a visual system with neuronal subpopulations with a multiplicity of target, cue, and location properties can be computationally compatible with a behavioral cueing effect.

### Relationship to Covert Attention Mechanisms Inferred from Human Psychophysics and Computational Models

In general, our CNN results show emergent mechanisms that have some commonalities with previously proposed models applied to visual psychophysics to make inferences of how attention alters visual processing^2,7,13,25,43,58,64^. The single-location TC neurons are similar to the idea of sensitivity changes prior to integration across locations in some of these models^2,13,64^, however, the underlying mechanism is different, arising through an interaction with the ReLU activation (rather than noise exclusion, internal noise reduction, or a pre-noise gain, etc.). The location-summation CNN units are more similar to the BIO prior weighting models^15,39,56^, showing that the location-integrated variables are more driven by sensory information from the cued location. Lastly, the location-opponent neurons are not typically explicit components of covert attention computational models, although opponency mechanisms are pervasive in visual processing of color^65^ and motion^66^.

At the system level, the results showed a correspondence between neuronal properties of the CNN and the stages of the BIO. However, the BIO shows defined transitions to target-cue integration and location integration stages, while the CNN had a distribution of neuronal properties showing different degrees and types of integration, varying sensitivities, and more nuanced transitions across layers.

### Comparing mechanisms for mice, non-human primates, humans, and CNNs

Covert attention in mice and monkeys is thought to be similarly mediated by excitatory influences of the cue on neuronal responses to a target^67^. In contrast, there is some indication of differing temporal dynamics, with mice having cue-related modulation more restricted to the presentation of the visual stimulus^19^ while monkeys have modulations that extend to post-cue, pre-stimulus periods^68^. In terms of mechanisms in the current paper, the location opponent mechanism might be implemented at different stages along the processing pathway. While we find opponent cells in mice SC, Herman et al., 2018^68^ did not explicitly report location-opponent cells in monkey SC. However, they successfully modeled the animals’ behavior in terms of a difference operation (opponency) between left and right SC activity, suggesting that a location-opponency operation might be implemented, not in SC as for mice, but downstream of SC for monkeys.

Finally, a hallmark finding in human covert attention is cueing effects with partially valid pre-cues followed by 100% valid post-cues^2,6,64^. Neither a BIO nor a CNN would predict these cueing effects without building some bottleneck in temporal processing, and thus these effects are unaccounted for by the current approach. Similarly, CNNs do not capture the time course of attentional effects^69^ and human cueing effects with non-predictive exogenous cues^2^. Finally, an assessment of how the results would be influenced by the addition of divisive normalization^14,58^, explicit biological attention^17,18^, or feedback connections^27,70^ into the networks is important.

### Broader implications

Many neuronal mechanisms of vision have been traditionally inspired by psychophysical results and/or hypothesized by researchers to account for behavioral findings. However, these proposed mechanisms often end up being a stylized version of what is observed in neuronal recordings (e.g., measured color tuning in cells depart from the prototypical theoretical R-G, Y-B opponency^71^). The proposed single-unit neuroscience-inspired CNN approach results in a distribution of cells that vary in their tuning and identifies novel mechanisms. Thus, it may provide neuronal predictions closer to those observed in the brain, including novel neuron types, and complement the stylized mechanisms and theories conceived by neuroscientists.

## Methods and Materials

### Human Data

We used the human subjects’ behavioral data from the Posner task in Srivastava et al., 2024^23^. The demographics and experiment design are described in their methods and repeated in Supplementary Section S.1b.

### CNN Architecture

We used a feedforward CNN with three convolutional layers with 8, 24, and 32 kernels, respectively, and kernel strides of size 3 by 3; and two fully-connected layers of size 256 and 2 (output layer). ReLU activation function was used for all layers except the output layer, where SoftMax activation was used.

### Training details

The training was done via gradient descent to minimize cross-entropy loss, with batch sizes of 32, until convergence, which was typically around ten epochs. We used 10000 training images, with half of the images containing no target, 4000 target-present images with valid cues, and 1000 target-present images with invalid cues for the main result with 80% cue validity. We tested on ten sets of 1000 images with the same relative proportion of the various stimuli types. For the models trained with 50% predictive cues, the number of target-absent images was kept the same (50%).

Ten different models were trained from scratch with random weight initializations for 80% predictive cues, and another ten for 50% predictive cues. No training history or weights were shared between any two models.

### ROC Analysis

#### Target vs. distractor at each location

The CNN was shown ten sets of 500 noisy target-present and 500 target-absent (distractor-present) images with no cue. We separately evaluated the left and right locations. The response of each neuron in the network to these stimuli categories was recorded. For each neuron’s pair of response distributions (target-present and absent), we systematically varied a decision threshold to generate a hit rate (proportion of target-present distribution to the right of the decision threshold), a false positive rate (proportion of target-absent distribution to the right of the decision threshold) and traced a ROC curve for detecting whether the target or the distractor was present at each location. We obtained a non-parametric Area Under the ROC (AUROC_t-d_) from the traced ROC curve.

Unresponsive neurons have only one point on their ROC curve since their response range is just one point. We assigned an AUROC of 0.5 to non-responsive neurons.

#### Cue presence vs. absence at each location

The CNN was shown ten sets of 500 noisy cue-present and 500 noisy cue-absent images, each with no target or distractor. We separately evaluated left and right locations. The response of each neuron in the network to the cue-present and cue-absent images was recorded. Using these responses, we constructed ROC curves for detecting cue presence/absence at each location and calculated non-parametric area under the ROC curve (AUROCc-nc)

#### Obtaining p-values of the neuronal AUROC

Each AUROC for each neuron was computed on ten sets of 1000 images each (500 for present, 500 for absent). We used the mean and standard error over these ten iterations to perform one-sided t-tests of being different from 0.5. Multiple comparison correction was performed, correcting for the False Discovery rate at α = 0.01.

#### Estimating CNN Neuronal Receptive Fields

We used ten sets of 40k noisy images with the target (14 degrees) and distractor (7 degrees) present at both locations with no angle noise to fit a linear regression model between input image pixels and neuronal responses. The superposition of the target and the distractor was done to ensure that all relevant pixels in the image had a baseline input required to cause a variation in the neuron output. The predictor variables were noisy pixels of cropped regions of size 100 by 170 from the input images that contained both locations. The noise level was the same as that used in training. Separate regressions were run for the cue-left and cue-right conditions for each neuron or BIO output depicted in Figure 6d-h.

### Mice SC neuronal data details

#### Neurophysiology experiment details

We reanalyzed data from Wang et al., 2022^19^ (Figure 7), which comprises the neuronal responses of four mice to a cued orientation change detection task with a 100% valid pre-cue. The pre-cue and the distractors were vertical Gabor patches, while the target was a Gabor patch tilted either 9 or 12 degrees. Cued trials started with a pre-cue appearing on one of two locations, followed by another distractor appearing at the uncued location, followed by the target at the cued location (for target-present trials). A control condition without the cue followed the same procedure, but without the appearance of the pre-cue. Only a subset of the neurons (n = 109) was recorded in both cued and uncued conditions. Out of these 109 neurons, 61 neurons were from sessions where the target orientation was 9 degrees, and 48 neurons were from sessions where the target orientation was 12 degrees. Our reanalysis focused on neurons that were recorded in both the cued and uncued conditions, because the computation of the cue AUROC was possible for these neurons. After 300 ms from the orientation change, a 500 ms response window started where mice were rewarded for correct decisions (using a licking response) on whether an orientation change was present or absent in the trial. The trials in which the mice responded before the response window started were discarded.

#### Neuronal AUROC analysis of the SC neuron data

We computed the AUROC for target detection by analyzing the distributions of spike counts in a 150 ms interval that started 60 ms after the orientation change and ended 210 ms after the orientation change and comparing it to the distribution of spike counts in a matched 150 ms interval in the no-change trials. Separate target detection AUROCs were calculated for trials with and without pre-cues (Fig 7).

The cue detection AUROC was computed by comparing the distribution of spike counts in the 150 ms interval beginning 60 seconds after the orientation change time point for cued vs uncued conditions. The cue detection AUROCs were computed on trials without an orientation change (target-absent).

The AUROCs and their p-values were computed using bootstrapping (consistent with Wang, 2022). For each neuron, we sampled responses (spike counts in the relevant interval) with replacement, from each response class (e.g., cue-present left responses vs. cue-absent responses for cue AUROC) to match the size (40) for each class in the original data. This was repeated 10,000 times. Each neuron’s AUROCs were tagged as significant if the AUROC was different from 0.5 on more than 9500 bootstrap samples (p< 0.05). To correct for multiple comparisons, we used false discovery rate (FDR) correction.

Supplementary section S.24 provides more details and example bootstrapped distributions of AUROC for example SC cells.

#### 3D CNN Model for the mouse task

Each stimulus was represented by a 3D array with 6 images of size 224 by 224. The first two of these images contained the pre-cue at one location (for cued condition), followed by two images with the distractor present on both locations (for target-present stimuli), followed by a target presented at the cued location (for target-present stimuli) in the next two images. Target absent stimuli contained four images with distractors on both sides on the third to sixth images and a cue on the first two images. Consistent with Wang et al.’s design, the cue and distractors were vertically oriented Gabor patches while the target was a Gabor patch oriented 9 degrees.

The model was a 3D CNN with 3 convolution layers with 8, 16, and 24 kernels, respectively. The kernel sizes and strides were kept the same in the spatial dimensions, while in the temporal dimension, the first two kernels had size 2 with a stride size of 1 pixel, while the third kernel had size and stride size of 1 in the temporal dimension.

The noise contrast was 0.14 while the contrast of the target, distractor, and cues was 0.18. The networks were trained to detect whether a target appeared in the stimulus. Other training details remained the same as the main networks used in the paper.

## Data and Materials Availability

All data gathered from this study and images used in the experiments are available upon request from the corresponding author and will be available publicly upon publication. The raw files for the mice SC data from Wang et al. 2022 can be obtained by request from the authors of that paper, but the processed files from our reanalysis of their data will be made available along with the rest of the data.

## Code Availability

All scripts are available upon request from the corresponding author and will be available publicly upon publication.

## Acknowledgments

The Institute for Collaborative Biotechnologies through US Army Research Office Contract W911NF-19- D-001.

## Author Contributions

S.S. and M.P.E. conceived the study, designed experiments, and interpreted the results. S.S. implemented the models. W.Y.W. provided expertise in CNN architecture selection. All authors wrote and edited the manuscript.

## Supplementary Text

The Bayesian ideal observer (BIO) derivation shown in this section is the same as used in Srivastava et al., 2024^1^. We reproduce it here for the sake of completeness.

### S.1a Bayesian Ideal Observer Models

General developments of the Bayesian ideal observers for the Posner cueing task can be found in many publications ^2–9^. The current treatment differs from previous Bayesian ideal observer treatments ^10–17^ in incorporating the process of the detection of the cue. Previous model treatments incorporate a prior reflecting the conditional probability of the presence of the target given the cue but implicitly assume the perfect detection of the cue. Here, we develop a theoretically more general model that explicitly calculates the likelihoods of the data given the joint presence of the cue and target, target alone, and cue alone (Figure S1.a). This is an image-computable Bayesian ideal observer^3,9,18,19^. Luminance noise is added to the image perturbing the luminance values of the target, distractor, and cue. When the cue visibility is high, the model makes identical performance predictions to the BIO model that only uses the priors and assumes perfect detection of the cues.

In the BIO framework, a decision across hypotheses ***s***_***i***_ is arrived at by calculating a posterior probability of each hypothesis given the sensory data, *P*(***s***_***i***_|***g***)^4,20^. Bayes’ theorem relates the posterior probability of a hypothesis to the likelihood of the observed sensory data given the hypothesis, *P*(***g***|***s***_***i***_) and the prior probability of the hypothesis, *P*(***s***_***i***_):

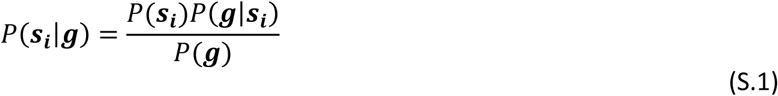

Typically, the denominator, *P*(***g***), which is the probability of the data over all possible hypotheses, is a constant across the various hypotheses. Thus, the BIO chooses the hypothesis with the highest product of likelihood and prior.

#### Calculating Likelihoods

Throughout the following sections, we will make references to the likelihood terms defined here. Suppose we have an n by m image (***g***) which contains either a luminance-defined element, a tilted line with orientation 1 (***s***_**1**_), or a tilted line with orientation 0 (***s***_**0**_) or a cue with additive Gaussian independent noise (with mean µ and standard deviation σ). Thus, each pixel luminance, *g*_*p*_ of ***g*** is a corresponding luminance pixel of *s*_*i*,*p*_ (where the first subscript refers to the hypothesis and the second subscript the location in the image) and some additive noise.

Because the noise added is Gaussian, the likelihood of observing a pixel value *g*_*p*_ given that the i^th^ signal is present is given by the normal probability density function:

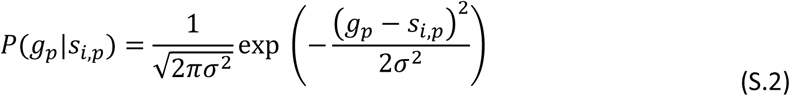

where σ is the pixel standard deviation of the luminance noise. We note that the likelihoods are defined with respect to pixel luminance values rather than hypothetical internal responses, feature values, or neural responses.

Because the noise added to each pixel is independent and identical, the joint likelihood of all pixels given the presence of the i^th^ signal, *P*(***g***|***s***_*i*_), reduces to the product of individual pixel likelihoods:

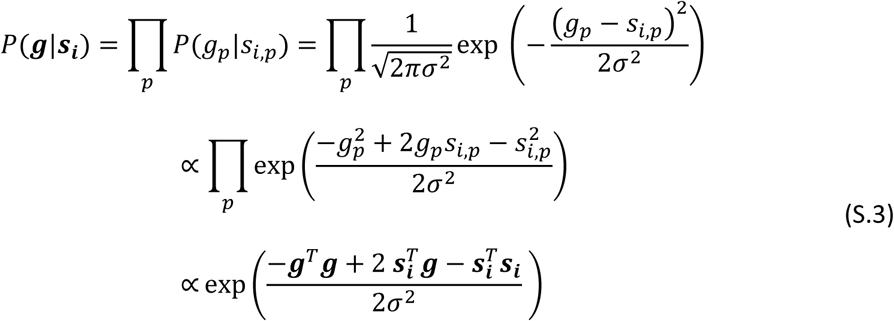

where ***g*** is a 1-D vectorized version of the image and ***s***_***i***_ is a vectorized version of the i^th^ signal. Eq (S.3) is a basic likelihood calculation and contains a template matching term ( ***s***_***i***_^𝑇^***g***).

#### Combining likelihoods across within hypothesis sub-events

Another important property that we will use repeatedly in our development of the BIO for the various tasks is that of summing likelihoods. In basic probability, the probability of an event A involves summing probabilities across all mutually exclusive events that qualify as A: P(A) = P(A1)+ P(A2)+…+P(An). Similarly, in many tasks, calculating the likelihood of the data given the presence of the target involves considering different locations, orientations, etc., for the target and distractors. The likelihood is calculated for each of the possible events and then summed to obtain a sum of likelihoods of the data given the presence of the target.

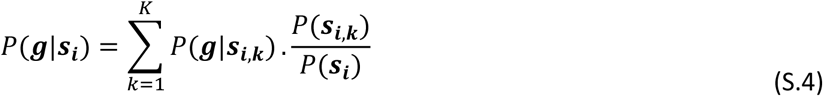

where 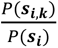 is the prior probability of the k^th^ event or target parameter value. Evaluating Equation S.4 requires the BIO to know apriori all possible values or locations attained by the target and distractors.

#### Posner Paradigm

In the Posner cueing task (Supplementary Figure S1a), the hypothesis ***s***^+^is that the target is present (target line with orientation 1 is at one location and a line with a smaller tilt, orientation 0, is at a second location), and the hypothesis ***s***^−^ is that the target is absent (both elements have lines with orientation 0). The box cue is present at either location with equal probability. The target appears with the cue with a probability of 80%. The likelihood of the sensory data, ***g***, given the presence of the target has to be summed across the mutually exclusive locations of the target.

**Supplementary Figure S1.**
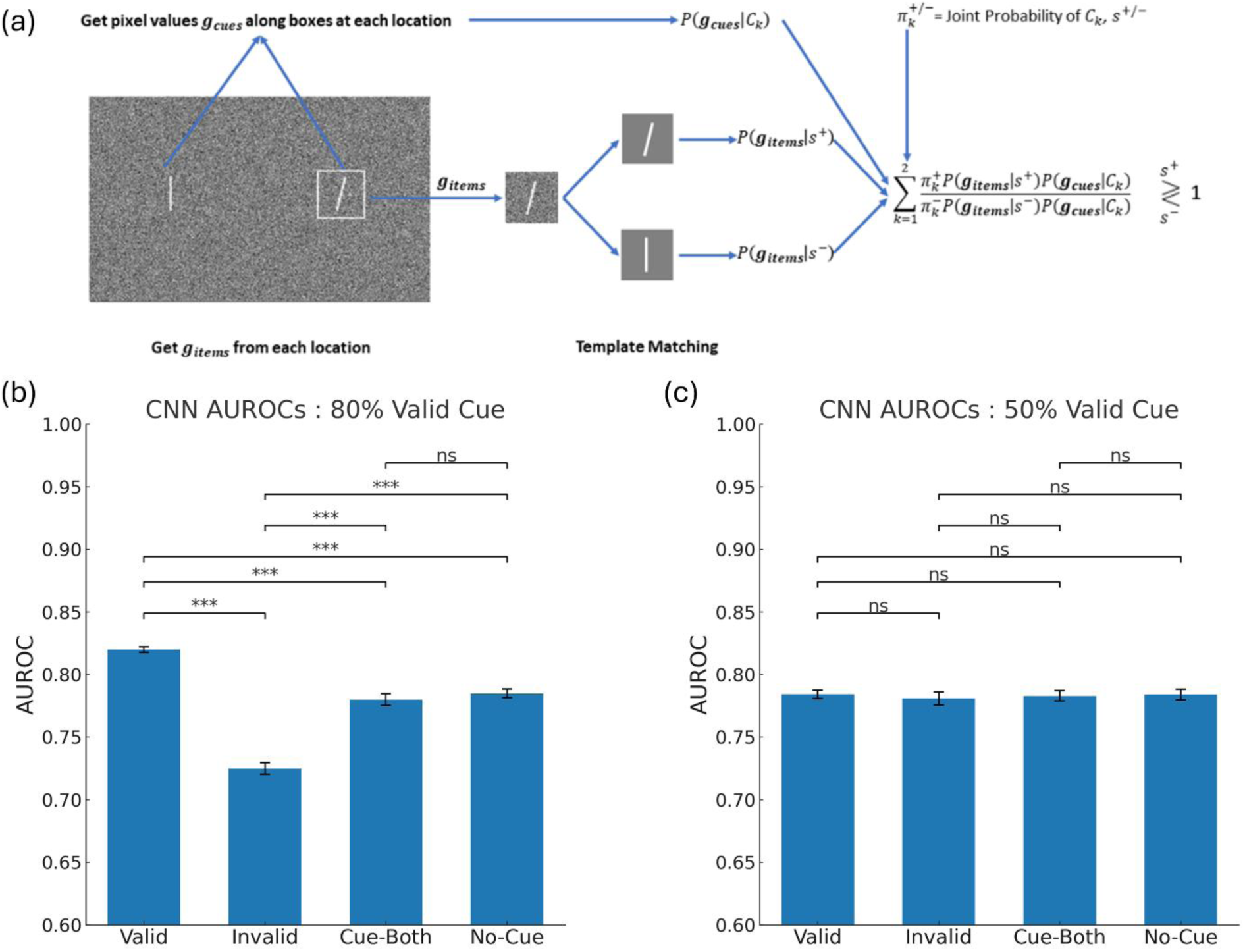
(a) Flow Chart for the Ideal Observer for Posner Paradigm. (b-c) CNN Output AUROCs for Valid, Invalid, Cue-Both, and No-Cue trials for networks trained on (b) 80% and (c) 50% valid cues. AUROC is used as a measure of target detection accuracy because the false positive rate of the CNN varies for the Cue-Both, No-Cue, and single-cue (valid and invalid) conditions. The AUROC combines hit rate and false positive rate into a single accuracy measure.

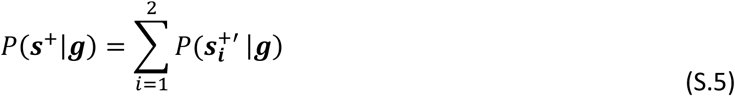

Where ***s***^+′^ stands for the target being present at location i, and the target being absent at the other location.

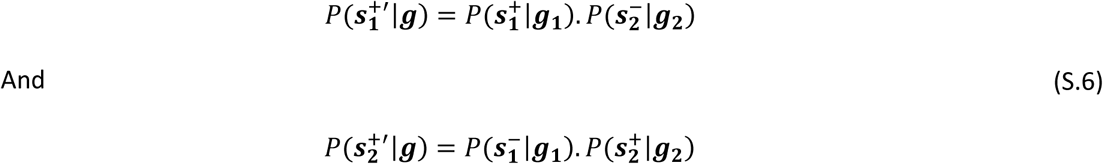

where ***g***_**1**_ and ***g***_𝟐_ are pixel values from the first and second locations, respectively.

Marginalizing over the two configurations corresponding to the cue appearing at the right or left location, we have

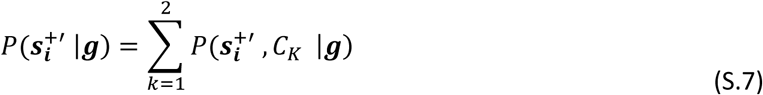

Where 𝐶_1_ refers to cue on the Left Location, no box on the right location and 𝐶_2_ refers to the cue on the right location, no cue on the left location.

Now,

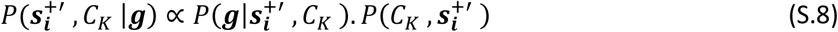

The likelihood can be separated into the independent contributions of the items (target/distractor) and the cues:

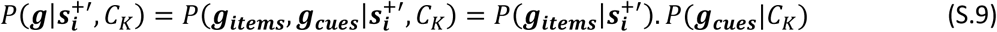

And

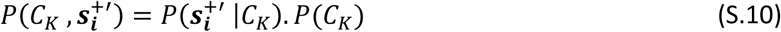

Thus, (S.9) reduces to:

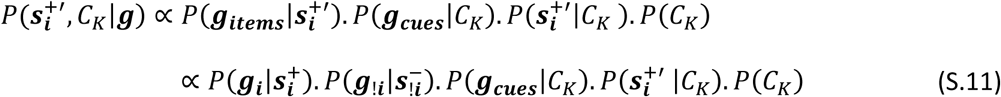

Where ***g***_***i***_ indicates pixels from location i, and ***g***_!***i***_ indicates pixels from the location other than i. ***s_i_***^+^ indicates target presence at location i, while ***s*_!*i*_**^−^ indicates distractor presence at the location other than i.

Now, in our implementation of the Posner Paradigm, the target angle is sampled from a Gaussian distribution with a mean of 15 degrees and standard deviation of 3.5 degrees, while the distractor angle is sampled from a Gaussian distribution with a mean of 7 degrees and a standard deviation of 3.5 degrees. However, since the number of pixels is finite, only a finite number of angles can be drawn on the actual images of the target and distractors. While calculating the likelihood of a target being present, we marginalize across all possible angles along with their prior probabilities.

Thus, using equation (S.4), it follows that:

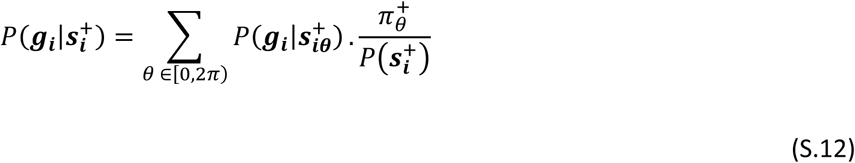

And

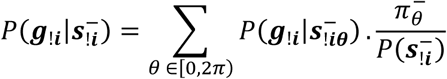

Where ***s***_***i***𝜽_^+^ denotes the presence of the target with an angle 𝜃 at location i, ***s***_!***i***𝜽_^−^ denotes the presence of the distractor with an angle 𝜃 at the location other than i, *π*^+^ is the prior probability of sampling a template with angle 𝜃 from the target angle distribution and *π*^−^ is the prior probability of sampling a template with angle 𝜃 from the distractor angle distribution.

Thus, combining equations (S.5), (S.7), (S.9), and (S.12), we get:

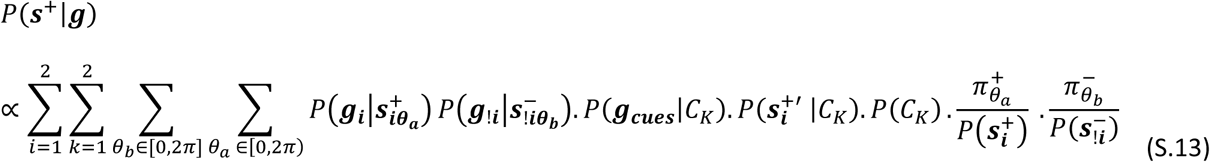

The first three likelihood terms in the summand here are obtained using (eq (S.3)), and the last four terms are known. The four summations are across target location (i), cue location (k), target angle (𝜃, and distractor angle (𝜃).

Likewise, to calculate the posterior probability for the target absence

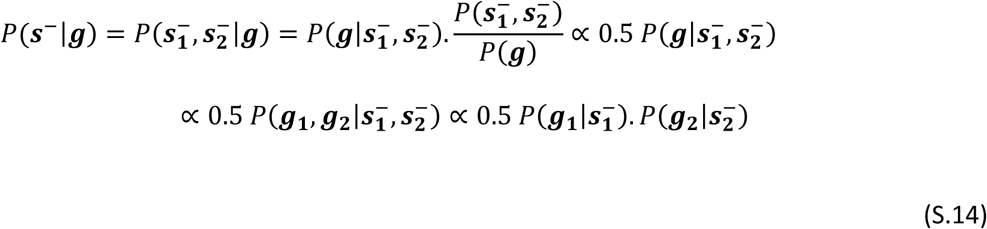

where ***s***_***i***_^−^ denotes the distractor being present at location i. *P*(***g***_**1**_|***s***_**1**_^−^) and *P*(***g***_𝟐_|***s***_𝟐_^−^) are again marginalized over all possible distractor angles similar to (S.12).

Finally, the BIO makes the decision: target absent, ***s***^−^, if *P*(***s***^−^|***g***) > *P*(***s***^+^|***g***) and target present, ***s***^+^, if *P*(***s***^−^|***g***) < *P*(***s***^+^|***g***).

#### Relation with the computational stages of the BIO in the main text (Figure 1c)

In the main text, we describe the computational stages of the BIO (Figure 1c). Here, we relate these stages to the formulation described in the sections above. In the first stage, the BIO detects the presence of the cue and target at each of the two locations by computing the likelihood of the data (noisy luminance values *G*_*Bi*_ at the pixels along a box at the i^th^ location) given the presence/absence of cue *P*(*G*_*Bi*_ |*Cue*_*i*_) and the likelihood of the data (noisy luminance pixels *G*_*i*_ from an i^th^ location) given the presence of the target or a distractor *P*(*G*_*i*_|*H*_*ji*_), where i denotes the i^th^ location and j denotes the j^th^ hypothesis (j=1, target present, and j=0 target absent, distractor present). This stage is denoted as cue detection and target detection in Figure 1c and is calculated using equation (S.3).

In the second stage, the BIO integrates target and cue information by multiplying the cue and target likelihoods and the conditional probability *π*_*i*_ of the target appearing with the cue at that location:

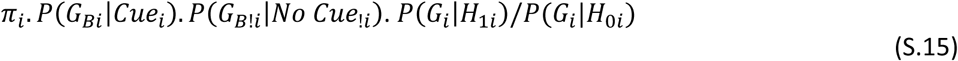

Here, !i denotes the location other than the i^th^ location. This calculation is separately computed for each location and denoted by the target-cue integration stage in Figure 1c and is equivalent to equation (S.11).

The third stage integrates the evidence across spatial locations by summing joint probabilities across possible target/cue and location configurations and is denoted as location integration in Figure 1c and is equivalent to equations (S.13) and (S.14).

### S.1b Psychophysics Task details

The task is the same as used in Srivastava et al., 2024. We reproduce it here for the sake of completeness.

#### Subjects

Eighteen undergraduate students from the University of California at Santa Barbara were recruited as subjects. The subjects all had normal or corrected-to-normal vision. Eleven subjects were women, and seven were men, all in the age range of 19-22. The subjects were naïve to the hypotheses of the study. Each participant provided informed consent to a protocol approved by the Institutional Review Board at the University of California, Santa Barbara.

#### Eye Tracking and Experiment Setup

The left eye of each participant was tracked using an SR Research Eyelink 1000 Tower Mount eye tracker sampling at 1000 Hz. Before each 100-trial session, a 9-point calibration and validation were done, with a mean error of no more than 0.5 degrees of visual angle. In each experiment, participants were asked to fixate at a central cross. If they moved their eyes more than 1 degree from the fixation cross, an error message would appear, and the trial would abort and restart with a new stimulus.

#### Stimuli and Psychophysics

Stimuli for all experiments were generated using MATLAB. The mean pixel value of the image backgrounds was 128. The minimum luminance was 0.08 cd/m2 and the maximum 114.7 Cd/m2. The luminance was linear with greyscale. The display had a resolution of 1920 by 1080 pixels. The viewing distance was 75 cm. The psychophysics experiment was written in MATLAB using the Psychophysics Toolbox.

#### Task

We implemented a forced-fixation, yes-no version of the classic Posner cueing task. Before each stimulus appeared on the screen, a blank screen with a fixation cross appeared on the screen for 250 ms. Participants were asked to fixate at the fixation cross at the center of the screen. Next, the stimulus was presented on the screen for 250 ms, with the fixation cross at the same location. An error message appeared if participants moved their fixation more than 1 degree away from the fixation cross, either on the pre-stimulus screen or while a stimulus was presented. When broken fixations were detected, the trial was restarted with a different stimulus. Following the stimulus presentation, a response screen was shown, asking participants to press the right arrow key if the target was present in the stimulus and the left arrow key otherwise. The response was recorded, and participants were asked to press the enter key to move on to the next stimulus.

Participants were shown 400 stimuli in total over four blocks of 100 stimuli each. The stimulus contained a target with a 50% probability. When a target was present, it was present with the box with 80% probability and on the other side of the box with 20% probability. These trials are called valid cue trials and invalid cue trials, respectively. The box cue appeared on either side with equal probability. The target angle was sampled from a Gaussian distribution with a mean of 15 degrees and a standard deviation of 3.5 degrees. The distractor angle was sampled from a Gaussian distribution with a mean of 7 degrees and a standard deviation of 3.5 degrees. The noise contrast was 0.14, while the target and the distractors had a contrast of 0.18. The cue contrast was 0.40.

### S.1c Statistical Efficiency

The external Gaussian noise added to the stimuli shown to the BIO and the CNN was varied to obtain the best fit to human observers. The goodness of fit was determined using the Akaike Information Criterion (AIC, see next section for details). The statistical efficiency^2^ (SE) is the ratio of the variance (RMS^2^) of the stimulus noise shown to the CNNs or humans compared to the BIO. The SE of humans was 0.39 (averaged across 18 observers) and that of the CNNs was 0.90, as reported in Srivastava et al., 2024^1^.

### S.1d Goodness of Fit Measure for the Models

The negative log-likelihood of the model predictions given human data was minimized to determine the best-fit model, while the external noise was varied using grid search. The Akaike Information Criterion was calculated using the log-likelihoods of the best-fit models using:

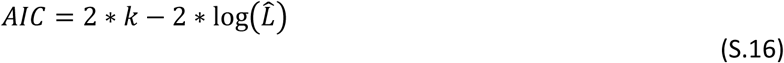

Where 𝐿^ denotes the likelihood of getting values equal to the model predictions if they were sampled from Gaussian distributions with means equal to the human performances and standard deviation equal to the human standard errors, and 𝑘 denotes the number of parameters varied (𝑘 = 1).

### S1e Testing cue-target learning in networks on cue presence on both locations (Cue-Both) vs. No-Cue trials

To further test whether the models learn the combination of target and cue as a consequence of the predictive nature of the cue, we tested the main networks on additional conditions. The network was trained on trials with a single cue present in all trials (as the CNNs in the main text) but tested on images with cues on both locations (Cue-Both) and images without a cue (No-Cue). We found that networks trained on the predictive cue (80% valid, Figure S1b) show no significant difference between the Cue-Both and No-Cue conditions. This result suggests that the networks’ improvement with the cue presence is a result of reducing the spatial uncertainty about which of the two possible locations contains the target. The presence of the cue at both locations does not improve the accuracy over no cues present. In addition, the presence of a single valid cue results in higher accuracy than the Cue-Both, No-Cue, and the invalid cue condition (see Figure S1b). Further support for the interpretation that the CNNs’ cueing effect arises from learning the predictive relationship between cue and target comes from training the network with a non-predictive cue. A non-predictive (50 % valid) cue results in no accuracy differences across all four conditions (valid cues, Cue-Both, No-Cue, and invalid cues; Figure S1c).

### S.2 Layerwise Distributions of Cue and Target AUROCs

Figures S.2. a-d show histograms of target AUROC across neurons and pie charts of the percentage of the different neuron types for the four layers. The results show a greater percentage of T+ and T- neurons as the processing progresses through the layers of the network (dense layer: 24.3 ± 1.29% T+ and 24.3 ± 1.68% T- neurons). Figure S.3.a-d shows histograms of cue AUROC across neurons and pie charts with the percentage of different neuron types for the four layers. The percentage of neurons tuned to the cue (C+ and C-) increased as processing progressed through the network. The target sensitivities increase as the processing progresses through the layers of the network: mean (p < 10^-5^ for all layers vs. dense), maximum (p < 0.01 for Conv1 vs. all layers) and mean of top 10% neuronal (p = 0.0022 for Conv2 vs. flat, < 10^-5^ for Conv2 vs. dense and 0.0005 for flat vs. dense).

**Supplementary Figure S2.**
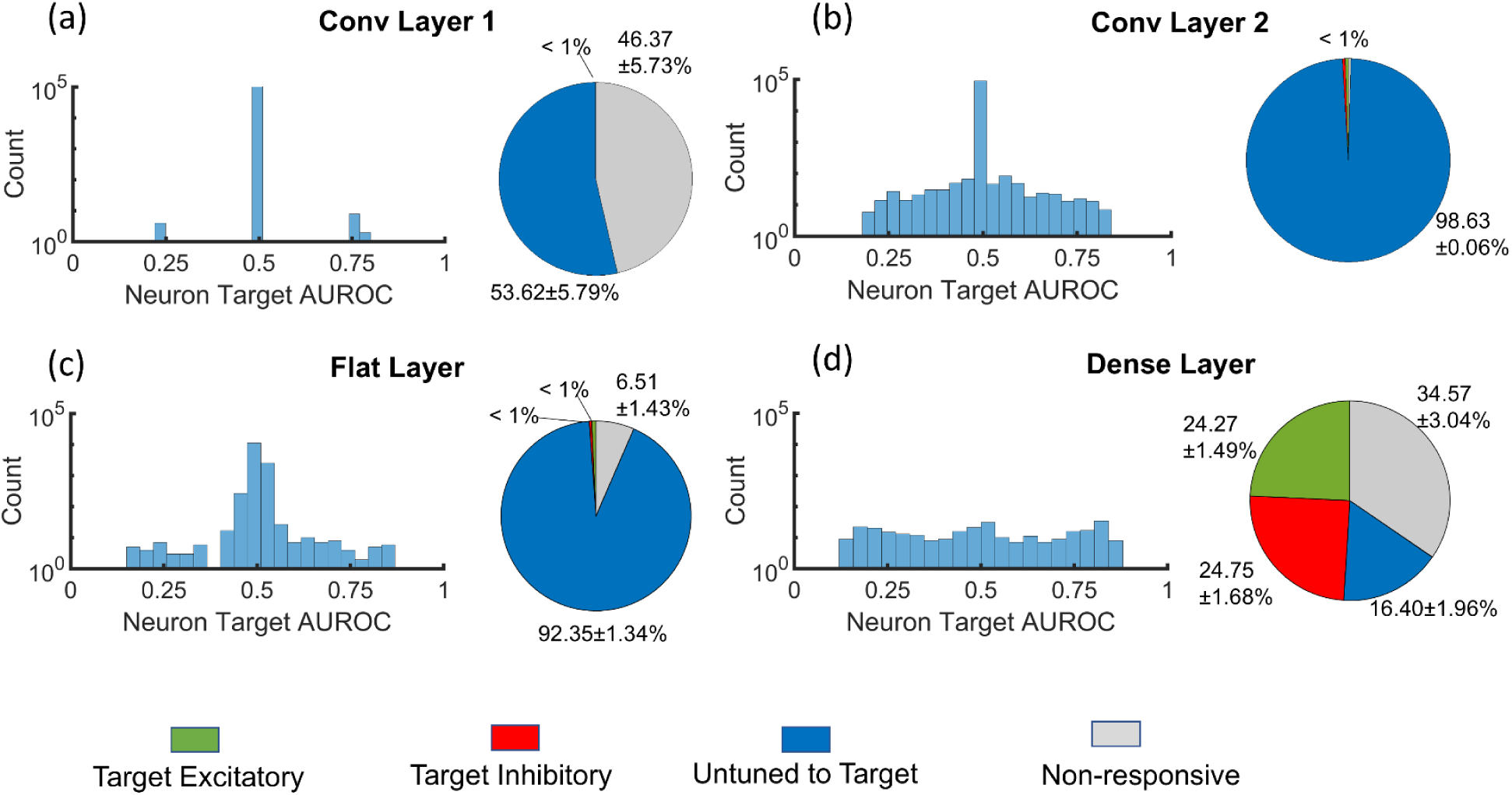
Target AUROC pie charts and histograms. First row (a-b): Histograms of target AUROC for the first two convolution layer neurons (the y-axis is on a logarithmic scale) and pie charts for various types of neurons based on their target tuning. Error bars denote the standard error of the mean over ten trained networks. Second row (c-d): Same as the middle row but for the flat and the dense layers.

**Supplementary Figure S3.**
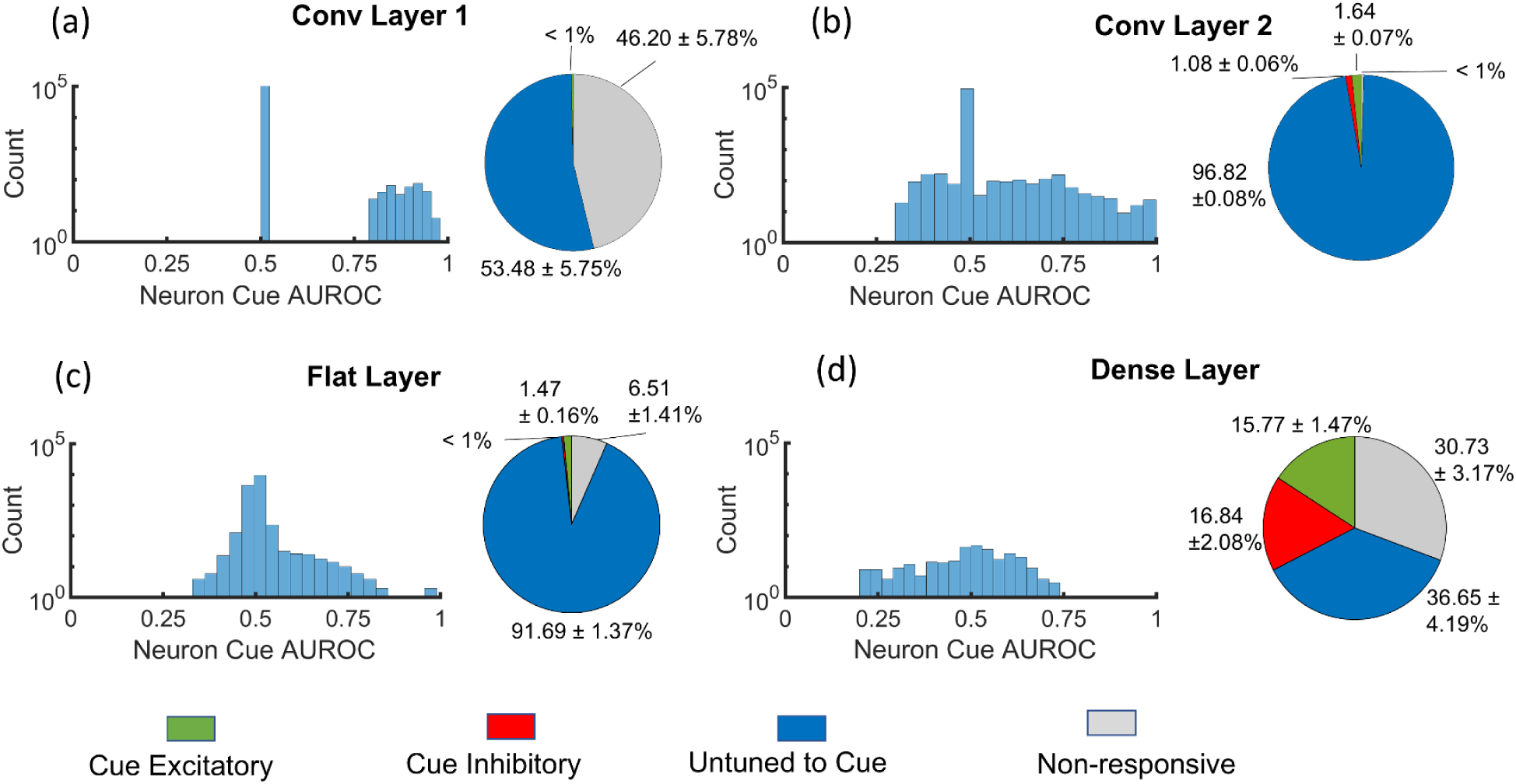
Cue AUROC pie charts and histograms. First row (a-b): Histograms of cue AUROC for the first two convolution layer neurons (the y-axis is on a logarithmic scale) and pie charts for various types of neurons based on their target tuning. Error bars denote the standard error of the mean over ten trained networks. Second row (c-d): Same as the middle row but for the flat and the dense layers.

### S.3 Early Layer Scatterplots for Models Trained on 50% Predictive Cue

The main text presents the scatterplots for target-cue integration (Figure 4) and location integration (Figure 5) only for the flat and dense layers for models trained on 50% predictive cues. Supplementary Figure S4 presents these scatterplots for the first two convolution layers. We find that, similar to the models trained on the 80% predictive cues, the first two convolution layers show no target-cue integration or location integration.

**Supplementary Figure S4.**
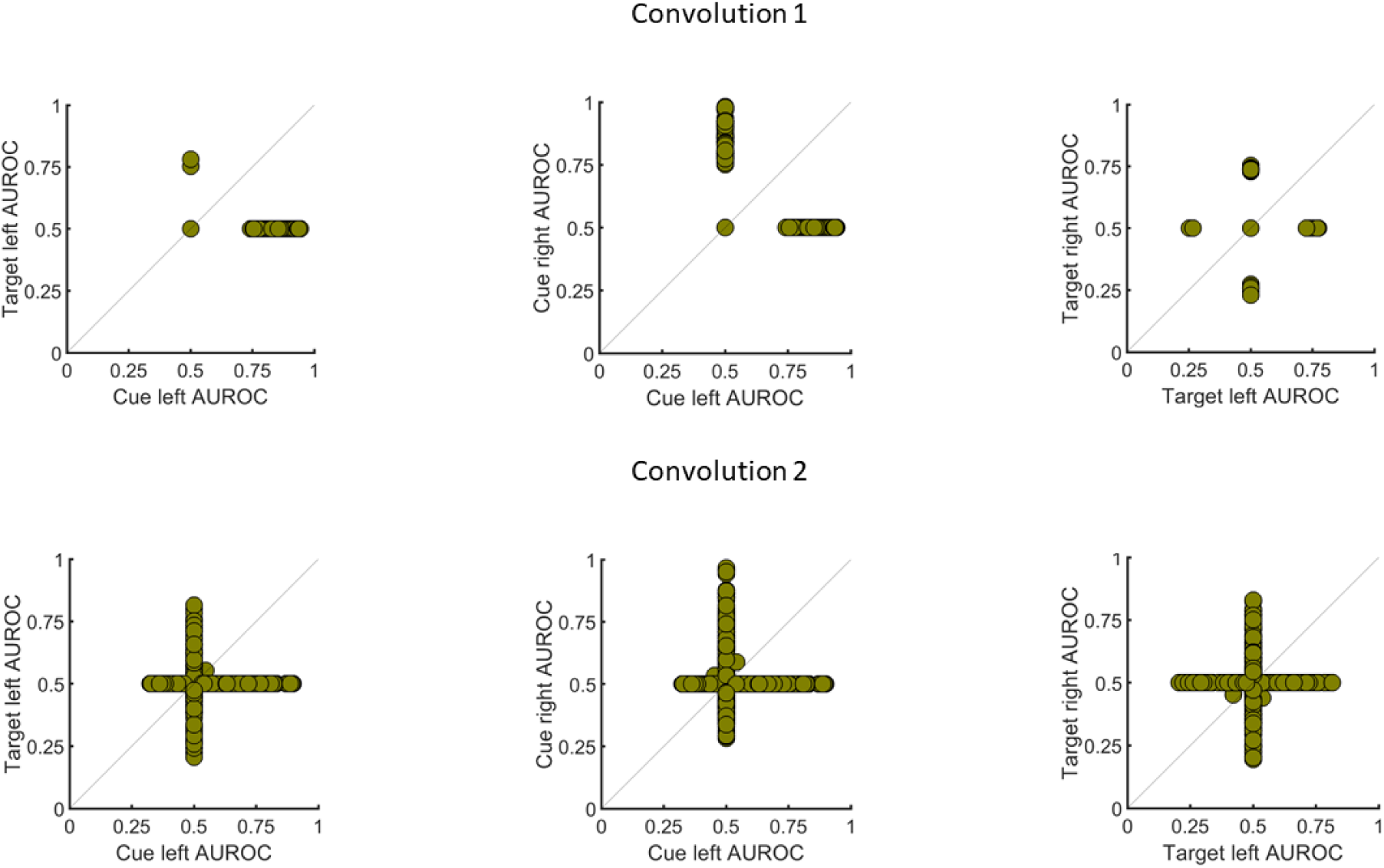
Cue-target and location integration in early layers for models trained on 50% predictive cue. Top row: Convolution layer 1, Bottom row: Convolution layer 2. No evidence for TC-integration (left graphs in both rows) or location integration in the early layers (center and right graphs in both rows), similar to networks trained on 80% predictive cue (main text Figures 4-5). The flat and the dense layers for models trained on 50% predictive cue are plotted in the main text, Figures 4-5.

### S.4 Early Layer Location Integration Scatterplots for Models Trained on 80% Predictive Cue

The main text presents the scatterplots for location integration only for the flat and dense layers for the models trained on 80% predictive cue (Figure 5). Supplementary Figure S5 shows the scatterplots for location integration in the first and second convolution layers. We find that the first two convolution layers do not exhibit any neurons jointly tuned to the two locations.

**Supplementary Figure S5.**
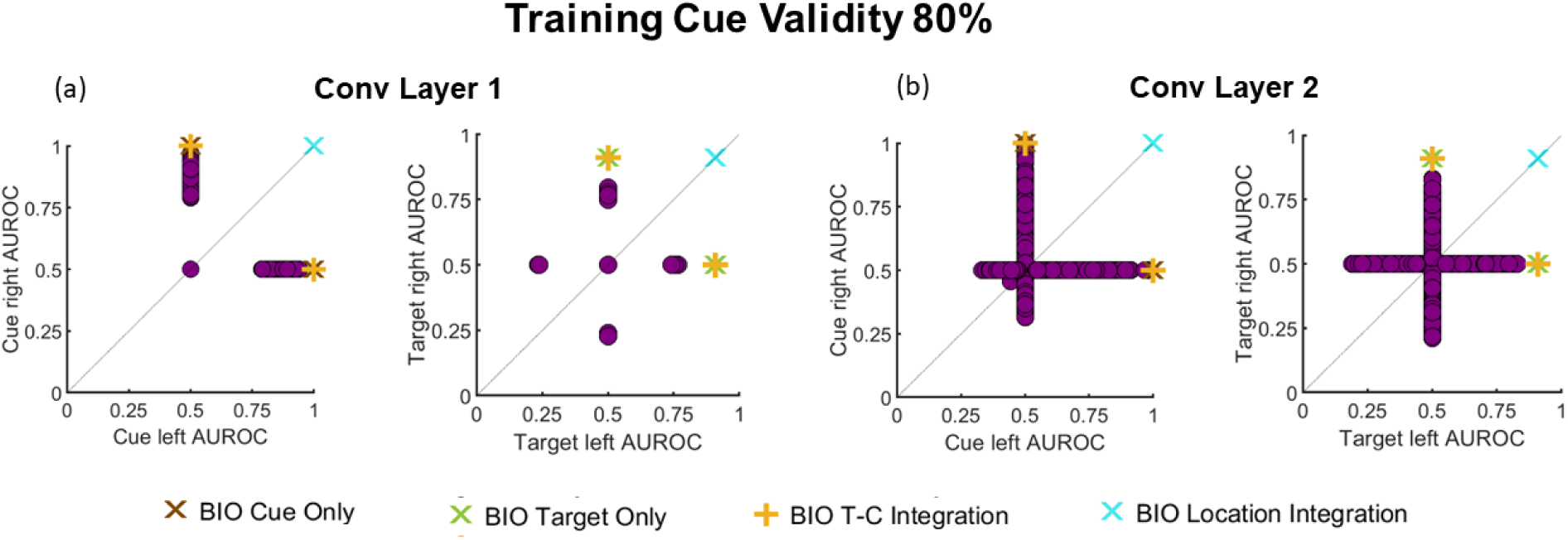
Location integration in early layers for models trained on 80% predictive cue. Scatter plots between neuronal AUROCs for cue detection and target detection at the right and left locations for neurons in the first two convolution layers for CNN with training cue validity of 80%. The AUROCs obtained from the corresponding stage of the BIO are also marked on the scatter plot.

### S.5 CNN Predictions for Mixed Trials: Valid/Invalid Cue, Neutral Cue, and No Cue

For the task with mixed-in neutral cues and no cues, we added two kinds of neutral cue trials: trials without any cue on either side (25%) and trials with a cue on both sides (25%). When the target was present in these trials (50% probability), it was equally likely to be present on either side. In the remaining 50% of the trials, the split between valid, invalid, and target-absent trials was the same as the main task (40%, 10%, 50%).

We trained a network with neutral stimuli, with cues at both locations, or no cues at either location, besides valid and invalid stimuli. The network’s valid hit rate was 0.78, invalid hit rate was 0.66, and false positive rate was 0.26. Additionally, the network’s accuracy on trials with cues on both locations was 0.71, and on trials with no cue on either side was 0.73.

**Supplementary Figure S6.**
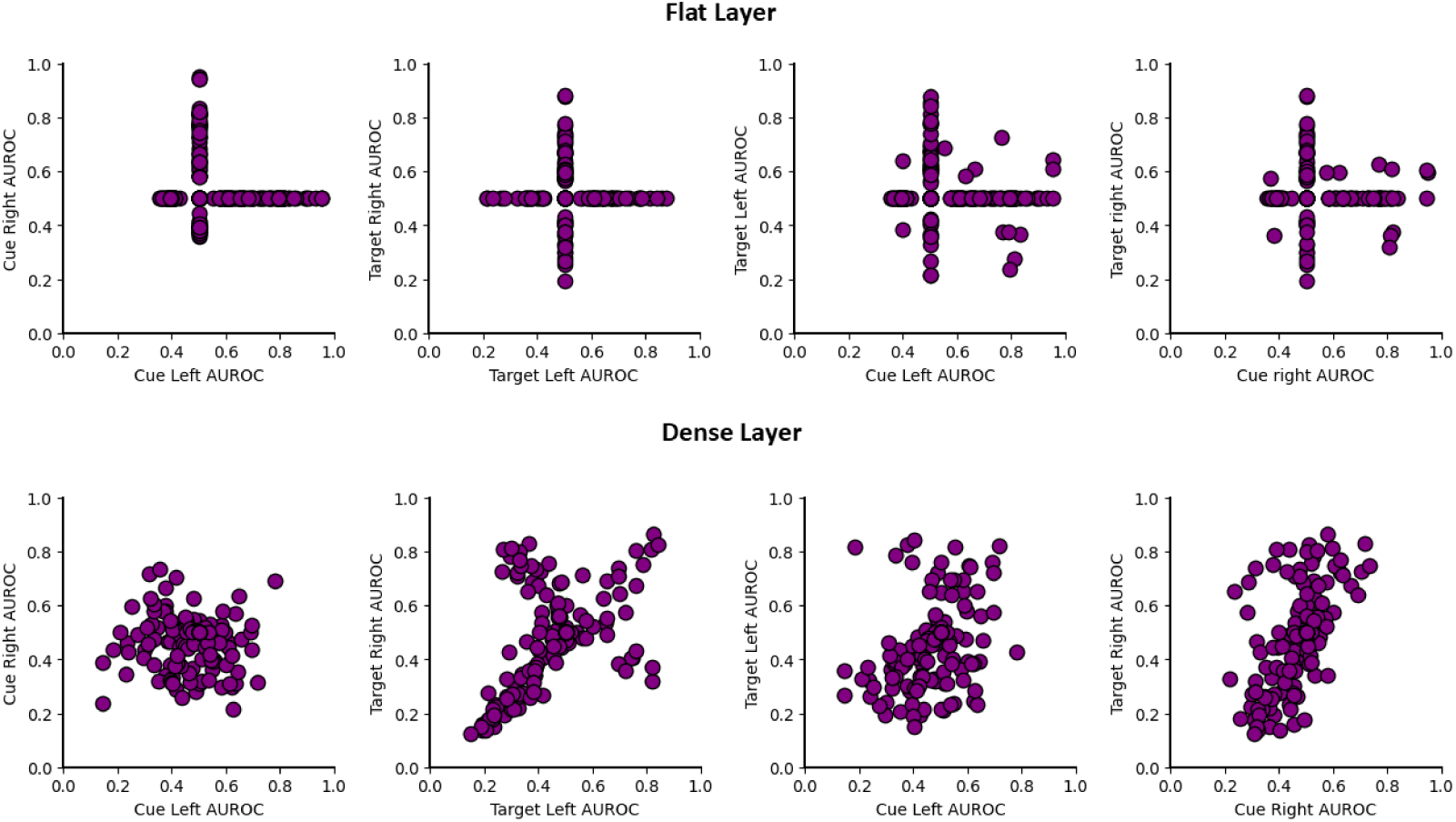
The effect of adding mixed-in neutral cues Scatter plots between neuronal AUROCs for cue detection and target detection at the right and left locations for neurons for the flat and dense layer of a CNN with training with mixed trials (25 % no cues, 25 % cues in both locations, and remaining other trials with 80 % valid cues)

### S.6 Violin plots for Layerwise Effect of Cue on AUROCs, Means, and Standard Deviations

In Supplementary Figure S7 we present the effect of the cue on the mean neuronal responses (S7 (b)-(c)) when the target is present, and on the AUROC for target detection for valid cue vs no cue (i.e., S7(d)) and no cue vs invalid cue conditions (S7(e)).

We also demonstrate (S7f) that the interaction with the ReLU thresholding is not essential for cueing in the dense layer neurons (compare with Figure 6d in the main text). Finally, we show that the ReLU is required for flat layer neurons to show cueing, but not for dense layers (S7(g)-(h)). Instead, the dense layer neurons require integration across the locations to show cueing (S7(h)).

Figure S7f (right) shows pre- and post-ReLU response distributions for a sample dense layer neuron. Dense layer neurons have higher means relative to the flat layer TC neurons. The distributions have a very small fraction of responses below zero, and thus, they are not affected as much by the interaction of the cue’s influence on the mean response and the ReLU thresholding. While pre-ReLU, the difference in AUROC without and with cue was 0.0418 in the example shown, post-ReLU, this difference is 0.0472.

The ReLU non-linearity has a larger influence at the flat layer TC neurons where the mean absolute neuronal cueing decreased from 0.034 to 0.010 (KS test: p < 10^-5^; Figure S7g) while the mean absolute neuronal cueing in the dense layer TC neurons decreased from 0.032 to 0.028 (KS test: p = 0.2219) after removing the ReLU (Figure S7h, left)).

We also assessed the functional importance of location integration in these units by removing the weights coming into the dense layer from the flat layer neurons tuned to the uncued location and found that the mean absolute neuronal cueing decreased from 0.032 to 0.011 (KS test: p < 10^-5^; Figure S7h, right), indicating that the integration across locations is required to give rise to the neuronal cueing effects in the location summation units of the dense layer.

In Supplementary Figure S8, we present the effect of the cue on the mean of neuronal responses when the target is absent. We also show the effect of the cue on the standard deviations of neuronal responses. On comparing the distributions of differences in the mean neuronal responses to the cue, we find that Flat TC and dense layers show a significant difference between networks trained with 80% and 50% predictive cues (KS-test: p < 10^-5^) for cue left vs. no cue and dense layer for cue right vs. no cue (KS-test: p < 10^-5^). On comparing the distributions of differences in the standard deviation of neuronal responses to the cue, we find that Flat TC and dense layers show a significant difference between networks trained with 80% and 50% predictive cues (KS-test: p < 10^-5^) for cue left vs. no cue and dense layer for cue right vs. no cue (KS-test: p < 10^-5^) both when the target is present and absent.

**Supplementary Figure S7.**
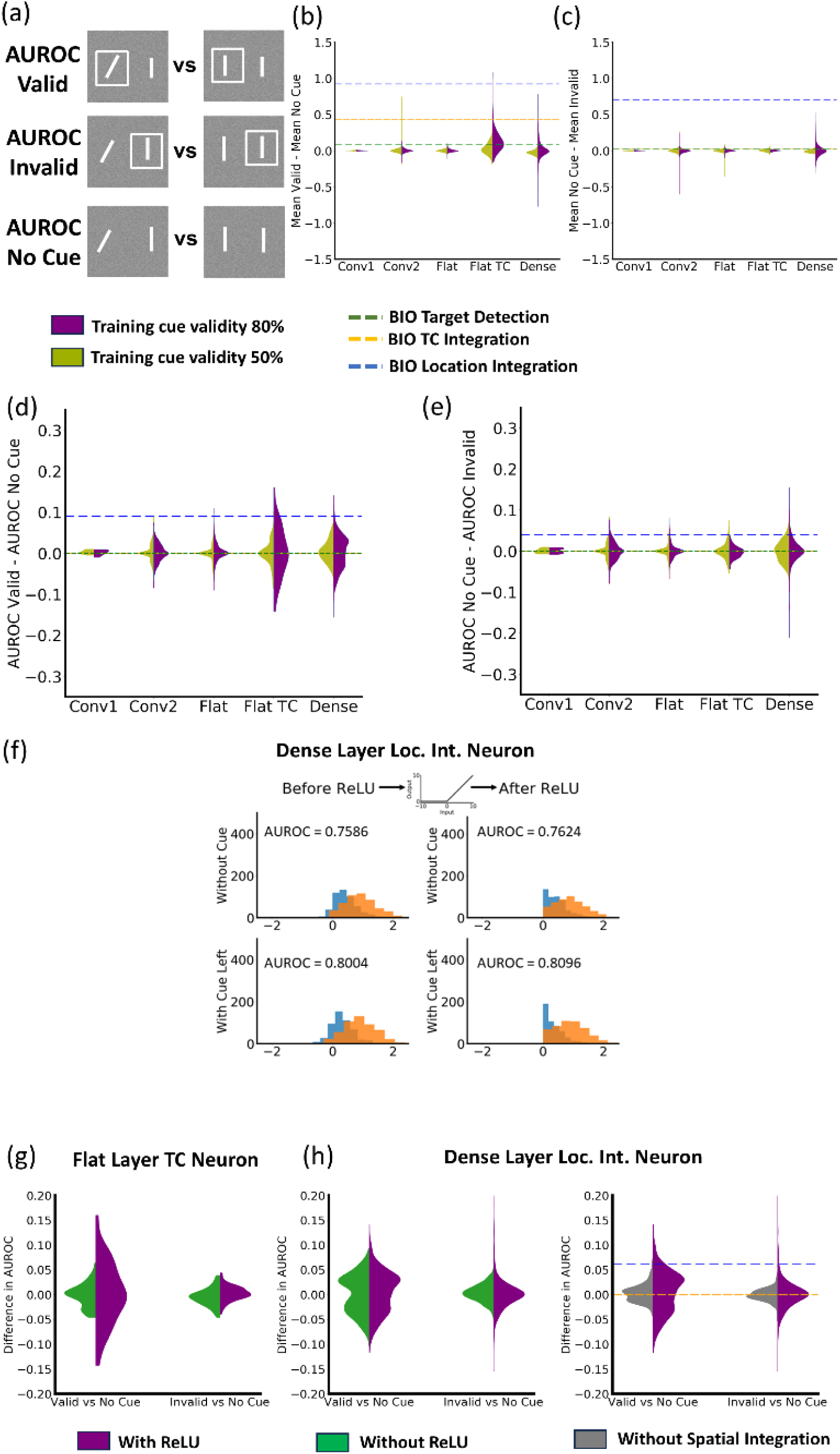
Distributions of the effect of the cue on mean responses and AUROC for target detection. (a) Example figures depicting stimuli for which the AUROCs are computed. (b)-(e) Layerwise distributions of (b) differences in mean neuronal responses between images with valid cue at the left location vs. no cue for networks trained with 50% (green) and 80% valid cues (blue), (c) differences in mean neuronal responses between images with invalid cue at the right location vs. no cue for networks trained with 50% (green) and 80% valid cues (blue), (d) differences in neuronal target presence AUROC between images with valid cue at the left location vs. no cue for networks trained with 50% (green) and 80% valid cues (blue), (e) differences in neuronal target presence AUROC between images with invalid cue at the right location vs. no cue for networks trained with 50% (green) and 80% valid cues (blue). Dotted lines represent the BIO. (f) Example responses before and after ReLU for an example dense location integration neuron (right half) when the target is present at the left location(orange) and absent (blue). The top row shows responses when the cue is absent, and the bottom row shows responses when the cue is present at the left location. There is less interaction between the cue and the ReLU for dense layer neurons because the mean responses of the distributions are higher relative to the threshold of the ReLU, thus, the cue has less of an effect. (g) Effect of removing ReLU (green) on the cueing of flat layer TC neurons relative to with ReLU (purple). Removing the ReLU for flat layer TC neurons has an important effect on the difference target sensitivity (AUROC). (h) Left: Effect of removing ReLU (green) on the cueing of dense layer location integration neurons relative to a condition with ReLU (purple). Removing the ReLU for dense layer TC neurons has small effect on the difference in target sensitivity (AUROC, left graph of (h)); Right: Effect of removing weights from the uncued location (gray, ‘without spatial integration’) on the cueing of dense layer location integration neurons relative to with ReLU and spatial integration. Analyses are presented for neurons tuned to the left location (convolution layers and flat layer), but similar results are obtained for neurons tuned to the right spatial location.

**Supplementary Figure S8.**
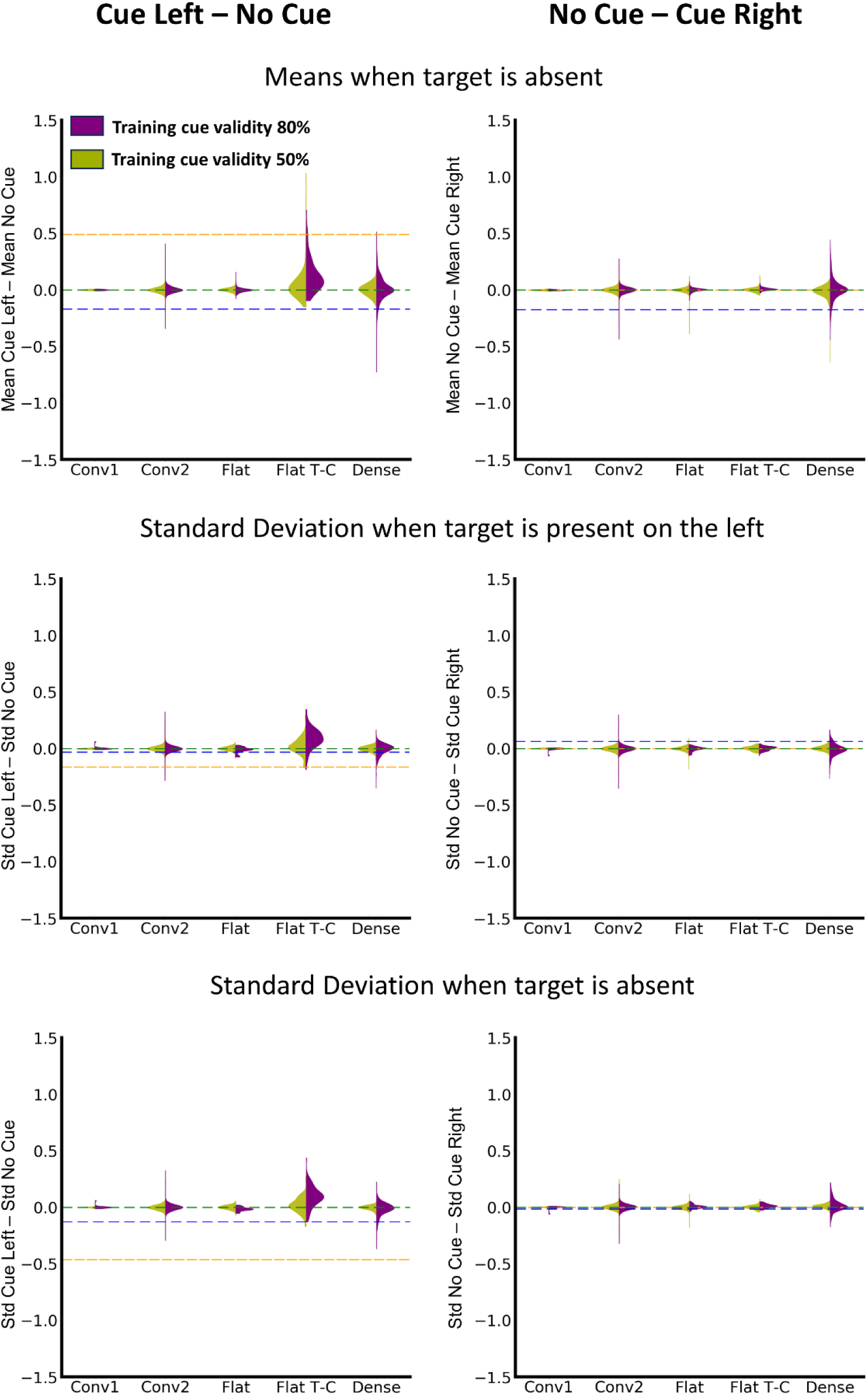
Distributions of the effect of the cue on means and standard deviations of responses. Left Column: Comparison between CNN neuronal responses per layer for cue left trials and no cue trials. Right Column: Comparison between CNN neuronal responses per layer for no cue trials and cue right trials. Top Row: Layerwise comparison between means of CNN unit responses with and without cue when the target is absent. Middle Row: Layerwise comparison between standard deviations of CNN unit responses with and without the cue when the target is present. Bottom Row: Layerwise comparison between standard deviations of CNN unit responses with and without the cue when the target is absent.

### S.7 The Relation Between Neuronal Cueing and Weights, and Other Dense Layer Mechanisms That Show Cueing

We computed the correlations between neuronal cueing of dense-layer TC neurons and the weights between these neurons and the target presence CNN output neuron. Across ten networks, neuronal cueing has a high positive correlation (r = 0.635) with the weights projected onto the target presence output neuron. For summation neurons, the correlation is 0.6624, for location summation + opponency neurons, the correlation is 0.5458, and for location opponency neurons, the correlation is 0.3977.

Besides the three dense layer mechanisms discussed in the paper, we find that among the dense layer neurons that show neuronal cueing, 4.28% are tuned to the target on one location and are cue opponent (T1Co), 2.24% are tuned to the target on one location and cue summation (T1Cs) and 0.25% are single location TC neurons.

### S.8 Receptive Fields Estimation for Flat Layer Single-location TC neuron and Dense Layer Summation + Opponent Neuron

In the main text, we presented the estimated receptive fields for a location integration neuron (Fig 6f), a location opponent neuron (Fig 6h), the CNN output (Fig 6i), and the BIO output (Fig 6j). In this section, we present the estimated receptive fields for a flat-layer single-location TC neuron (Supp Fig S9a-b), and a dense layer summation + opponency neuron (Supp Fig S9c).

#### S.8a. Flat Layer Single-location TC neuron

Unlike the receptive fields shown in the main text, the flat TC neuron weights only the left location. When the cue is present at the left location, the unit’s output is more driven by the image information because the response distributions are above the ReLU threshold. This results in a higher amplitude RF.

#### S.8b. Dense Layer Summation+Opponent Neuron

The dense layer summation + opponency neuron (Fig. S9c) weights both locations, but has a higher weight for the right location than the left. However, it is cue inhibitory on the left and cue excitatory on the right, and its weight of both locations increases with the cue on the right than on the left. However, irrespective of the placement of the cue, the right location is weighted higher than the left.

**Supplementary Figure S9.**
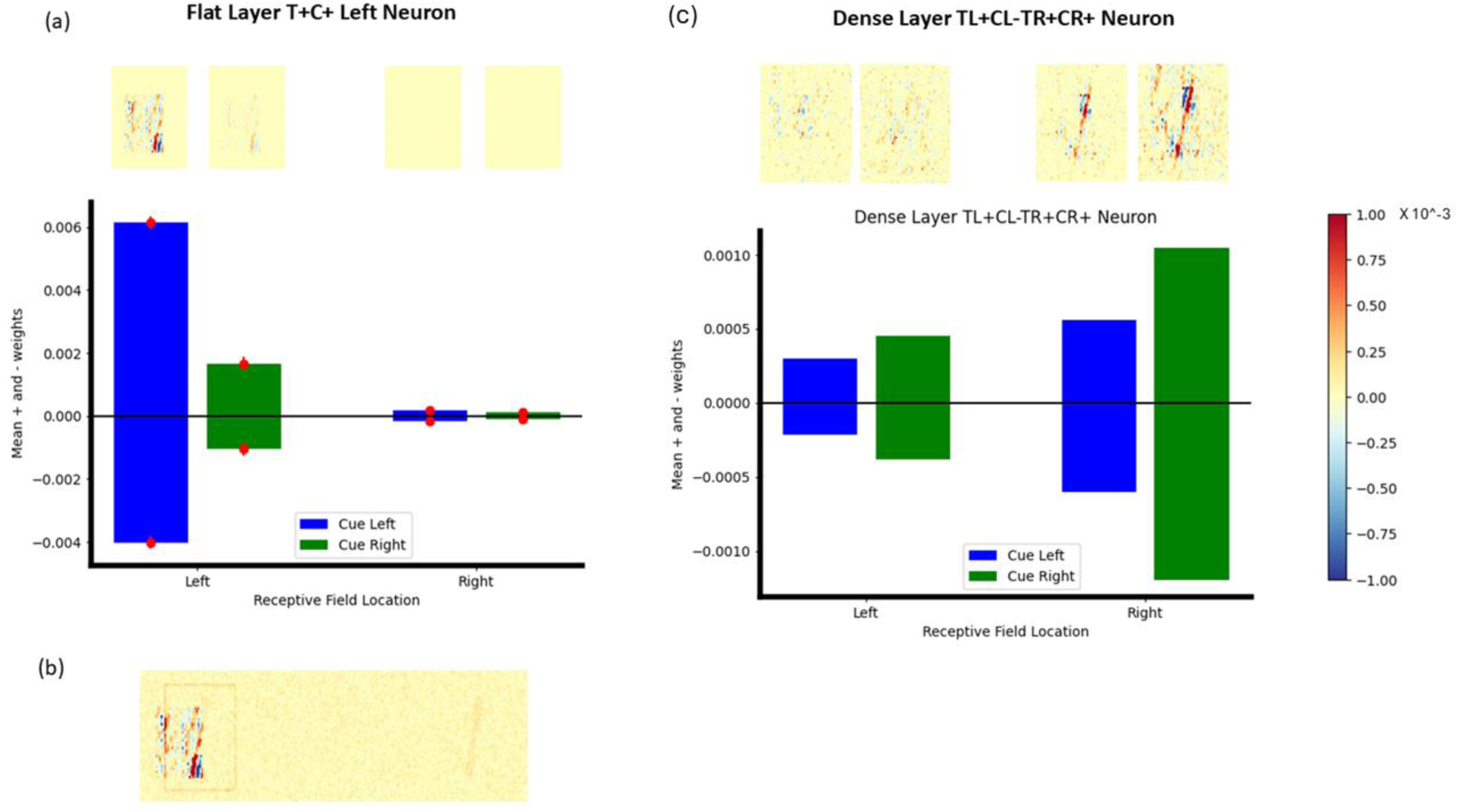
(a) Estimated Receptive Field of a flat layer T+Left C+Left neuron at the two locations when the cue is on the left (blue bars, and the RFs shown above them) vs. when the cue is on the right (green bars, and the RFs above them). (b) The RF overlaid on a sample target left trial. The box cue (orange box outline on the left) and the distractor on the right location (orange shade) depicted in this figure are not a part of the RF and only serve the purpose of indicating where the neuron’s RF is relative to the stimuli. The shown RF overlaps with both the cue region and the target region. (c) Estimated Receptive Field of a dense layer TsCo neuron at the two locations when the cue is on the left (blue bars, and the RFs shown above them) vs. when the cue is on the right (green bars, and the RFs above them).

### S.9 Effect of Changing the Cue Validity

Changing the cue validity was reflected in changes in the dense layer neuron AUROCs, particularly the neurons that are tuned to both the target and the cue (TC neurons). When the cue validity is increased, the dense layer neurons that integrate cue and target information show a larger range of cue AUROC while the range of target AUROCs does not vary significantly (Supplementary Figure S10, first two rows).

**Supplementary Figure S10.**
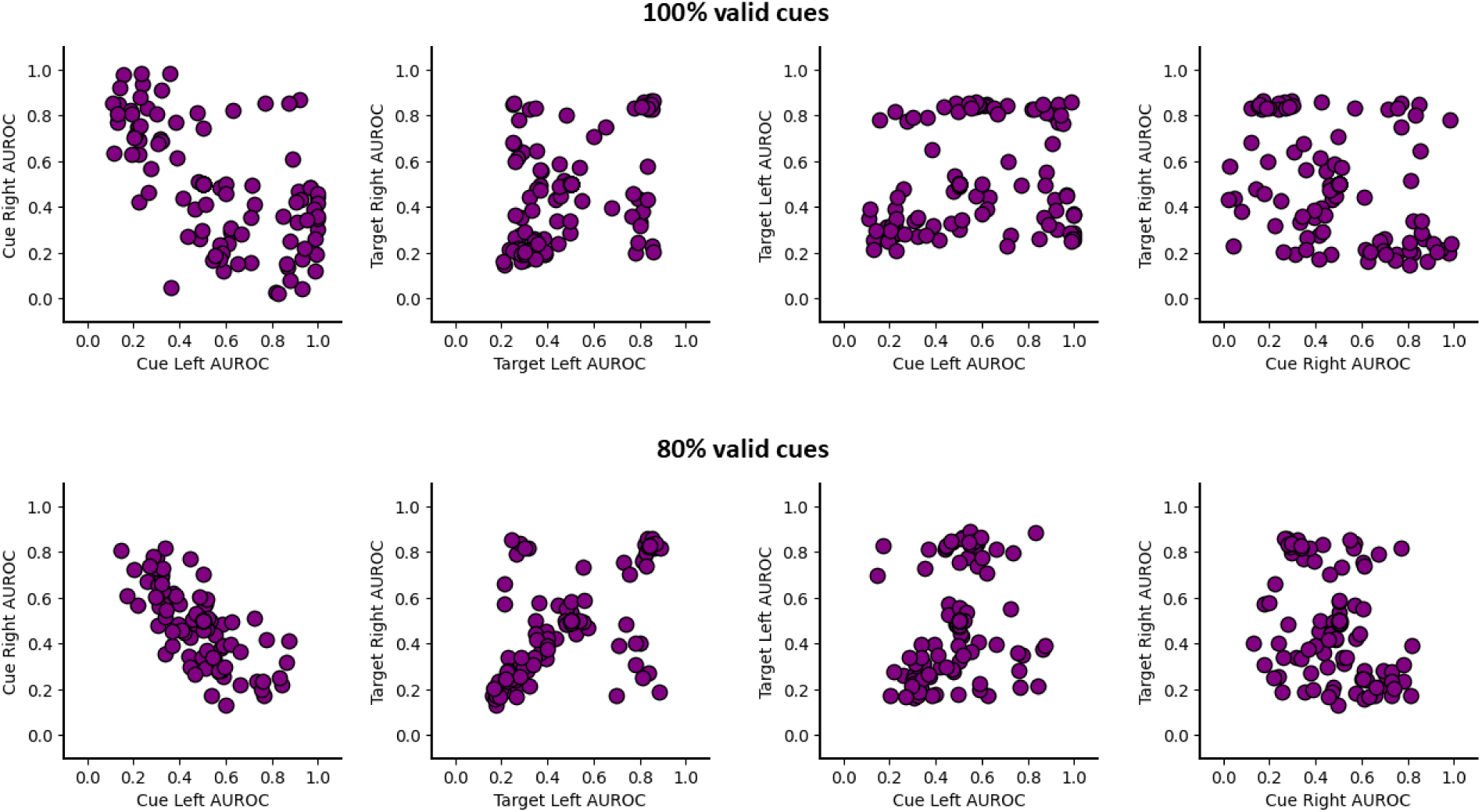
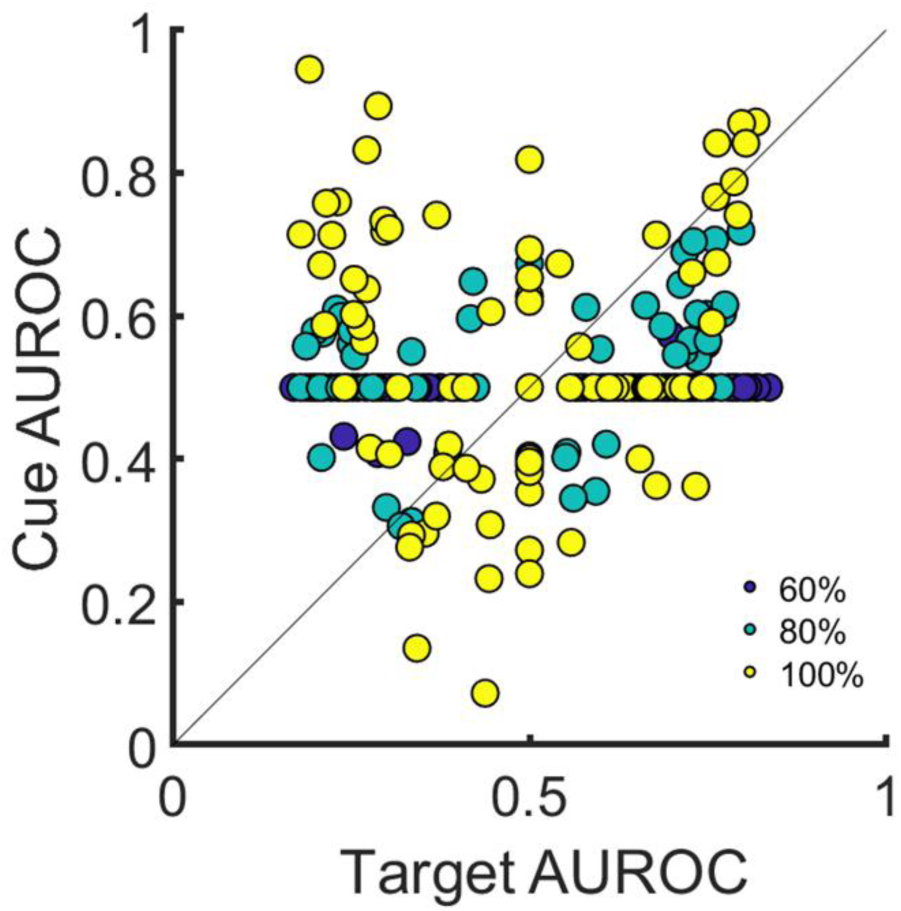
The effect of changing cue validity on the dense layer neurons. Top row: Scatter plots for cue right AUROC vs cue left AUROC, Target right AUROC vs target left AUROC, Target left AUROC vs Cue left AUROC, and Target right vs cue right AUROC for the dense layer of the model trained with 100% valid cues. Second row: Scatter plots for cue right AUROC vs cue left AUROC, target right AUROC vs target left AUROC, target left AUROC vs cue left AUROC, and target right vs cue right AUROC for the dense layer for the model trained with 80% valid cues. Third row: The AUROC scatter plots for cue-target integration in the dense layer are shown for three different cue validities (60%, 80%, and 100%). The bluer colors correspond to lower validity. The range of the cue AUROCs increases while that of the target AUROCs is nearly the same. The results shown are for the right location but generalize to the left location.

### S.10 The Effect of Regularization

Regularization is a machine learning technique that forces the network to optimize under constraints on total weights, biases, or activations. We assessed the influence of the L1 weight regularizations on the resulting emergent neuronal properties. For each of five regularization parameter values, we measured the accuracy (Figure S11a), cueing (Figure S11a), lifetime sparsity^21,22^, excess population kurtosis^23^, and excess weight kurtosis^24^.

The lifetime sparsity^21^ (Figure S11b) is calculated as:

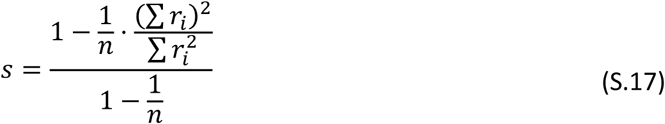

Here, 𝑟_*i*_ is the response of a neuron to image *i*, and n is the number of images. We calculate *s* for each neuron in the dense layer and report the mean across all neurons.

The excess kurtosis of a distribution measures the difference between the kurtosis of the distribution and a normal distribution. The excess population kurtosis (Figure S11c) of a distribution of neuronal responses to a stimulus is calculated as:

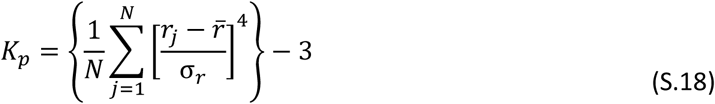

Here, 𝑁 is the number of neurons, 𝑟_*j*_ is the response of the j^th^ neuron to a given stimulus, 𝑟̅ is the mean and 𝜎_𝑟_ is the standard deviation of the distribution of neuronal responses to the stimulus. We calculate the kurtosis for each test stimulus and report the mean across stimuli in Figure S11. For excess population kurtosis, we considered the pre-ReLU activations of the neurons because the measure is usually not preferred for nonnegative distributions^23^. Figure S11c shows how excess population kurtosis increases with regularization parameters.

The excess weight kurtosis is calculated over the distribution of weights from neurons in the dense layer to output neuron 1 instead of the distribution of neuronal responses (population kurtosis). Sparser distributions have a higher kurtosis. Figure S11d shows how excess weight kurtosis increases with the regularization parameter.

To summarize, L1 regularization resulted in more sparse representations as measured by the lifetime sparsity of the dense layer neurons, excess population sparsity, and the excess kurtosis of the weights projected by them into the output layer. Importantly, we obtained levels of sparsity that are comparable to those reported by Vinje and Gallant (2000)^21^ that resulted in high levels of target detection accuracy and cueing effects (Figure S11a) and the dense layer mechanisms we report for the CNN without regularization. For regularized networks at regularization parameters 0.0025 and 0.005, we find all four mechanisms, although fewer as expected from regularization (Figure S11e; Figure S11f). More extreme L1 regularization (regularization parameters 0.0075 and 0.01) resulted in lower cueing, fewer cue-tuned neurons, and a lower extent of cue-tuning in those neurons, indicating that the network started ignoring the cue instead of the uncued location when trying to minimize resource usage.

**Supplementary Figure S11.**
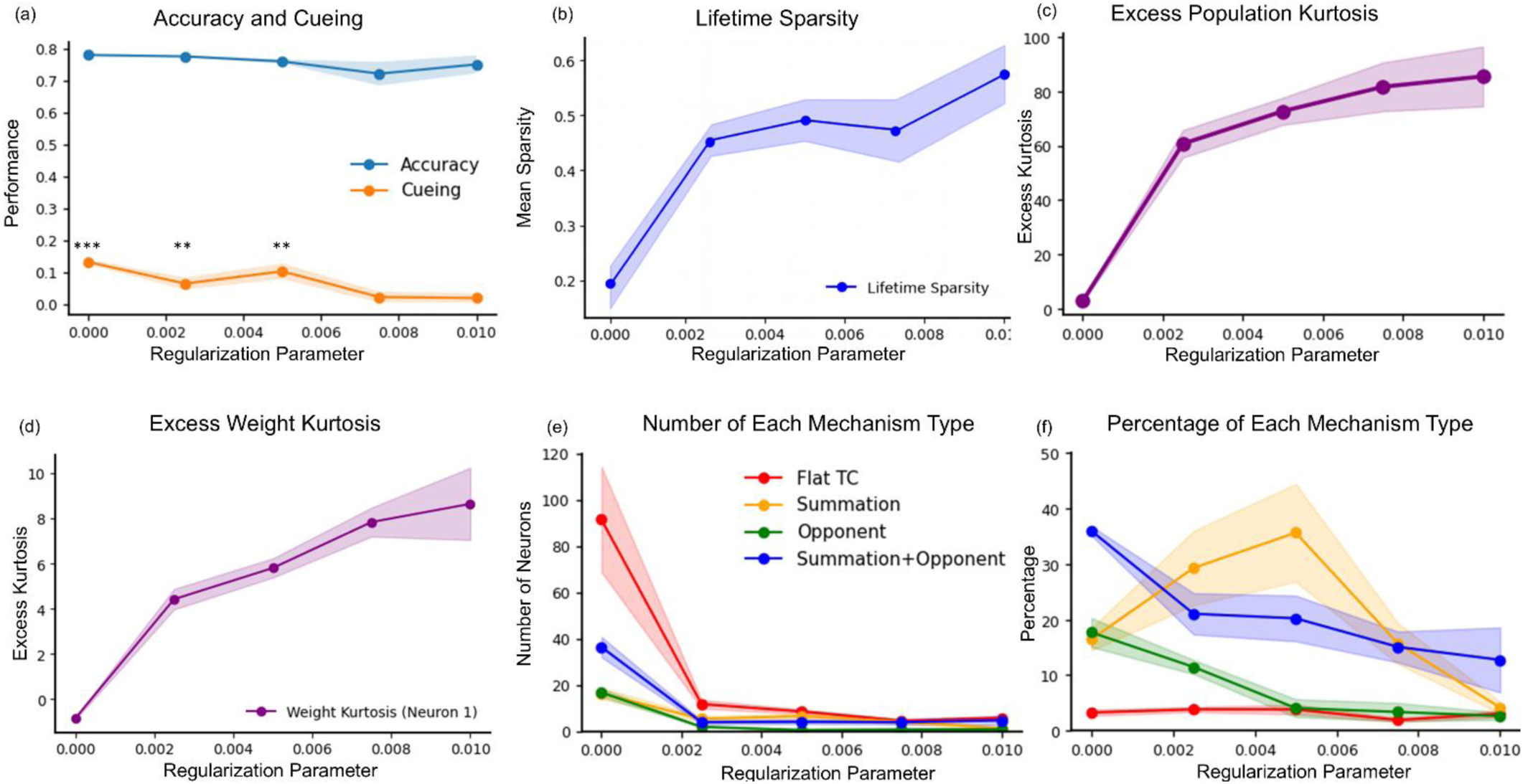
The effect of weight L1 regularization on the dense layer neurons. (a) Accuracy (proportion correct), cueing (hit rate valid cue trials – hit rate invalid cue trials, asterisks denote significant difference from zero), (b) lifetime sparsity, (c) excess population kurtosis of the dense layer neurons, (d) excess weight kurtosis of the weights from the dense layer neurons to the output layer, (e) number, and (f) percentage (as a fraction of total number of active neurons in that layer) of each mechanism type found in the CNNs as a function of the regularization parameter for weight L1 regularization applied to the dense layer.

### S.11 Effect of Changing the Size of the Dense Layer

We assessed the generality of results with variations in the size of the dense layer. We evaluated five different sizes of the dense layer, including 32, 128, 256, 512, and 1024. On increasing the size of the dense layer, more untuned neurons emerge but the cue-target scatter plots, the cue on both sides scatter plots, and the target on both sides scatter plots continue to show qualitative consistency with the results in the main text with a dense layer size of 256 (See Supplementary Figure S12; the results shown here for a dense layer size of 256 are from a different network and thus show small differences from the graph in the main text).

**Supplementary Figure S12.**
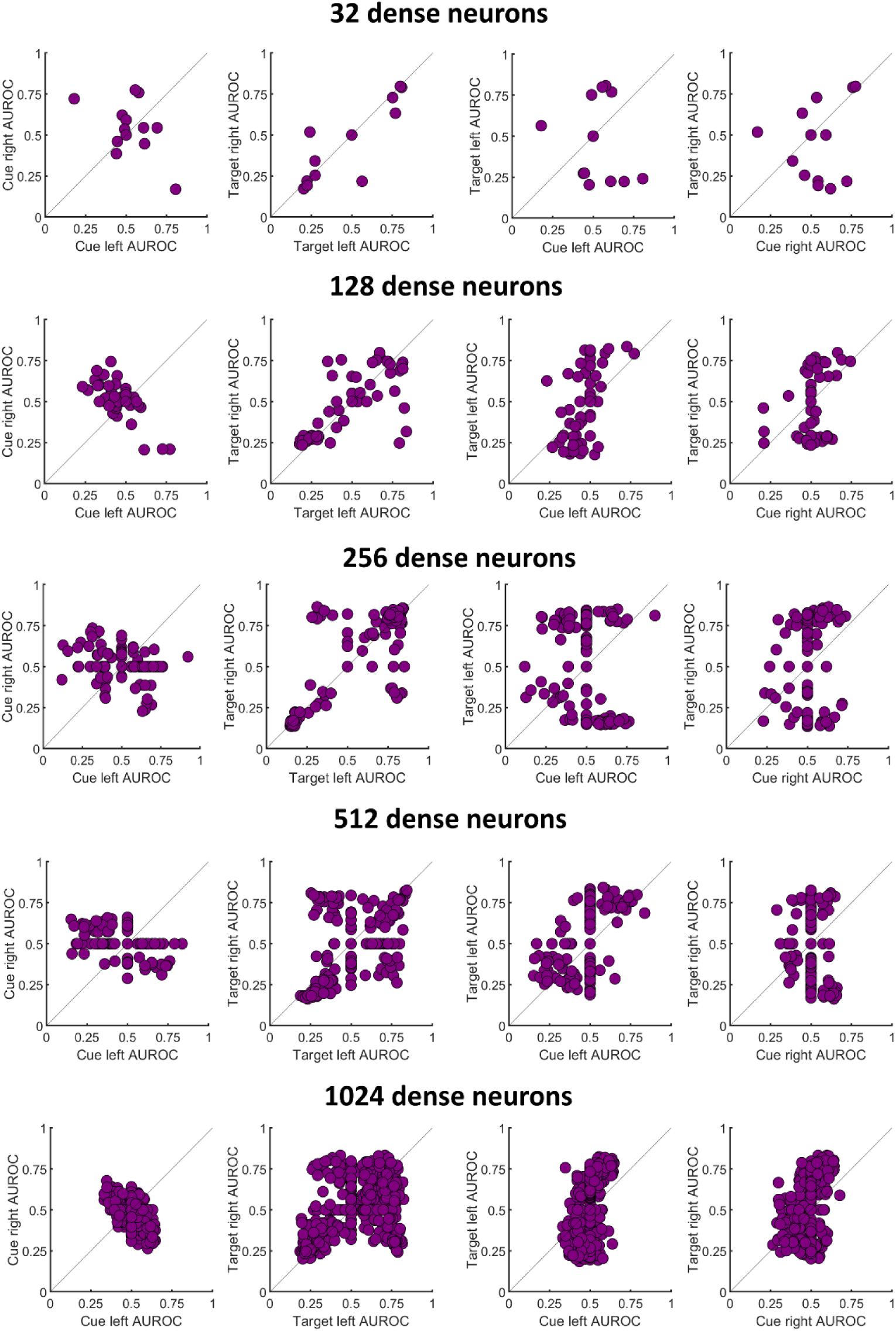
The effect of changing the size of the dense layer. The rest of the network architecture was kept the same as the network in the main text. Both target-cue integration and location integration are seen across all sizes.

### S.12 Effect of Increasing the Number of Layers

We tried two ways of increasing network depths: adding another dense layer or adding one or two more convolutional layers. The neuronal properties depend on which integration stage the neuron is in, and not the depth of the network. The network performance stays in the same range in all cases (between 0.73 to 0.75).

#### S.12(a) The Effect of Depth of the Network: Adding Another Dense Layer

When an additional dense layer is added, the CNN shows similar integration across locations and of cue and target as the first dense layer (Supplementary Figure S13). Adding the dense layer did not greatly change the output correlation with the BIO (0.6612 vs 0.6832 noted for the ten networks above).

**Supplementary Figure S13.**
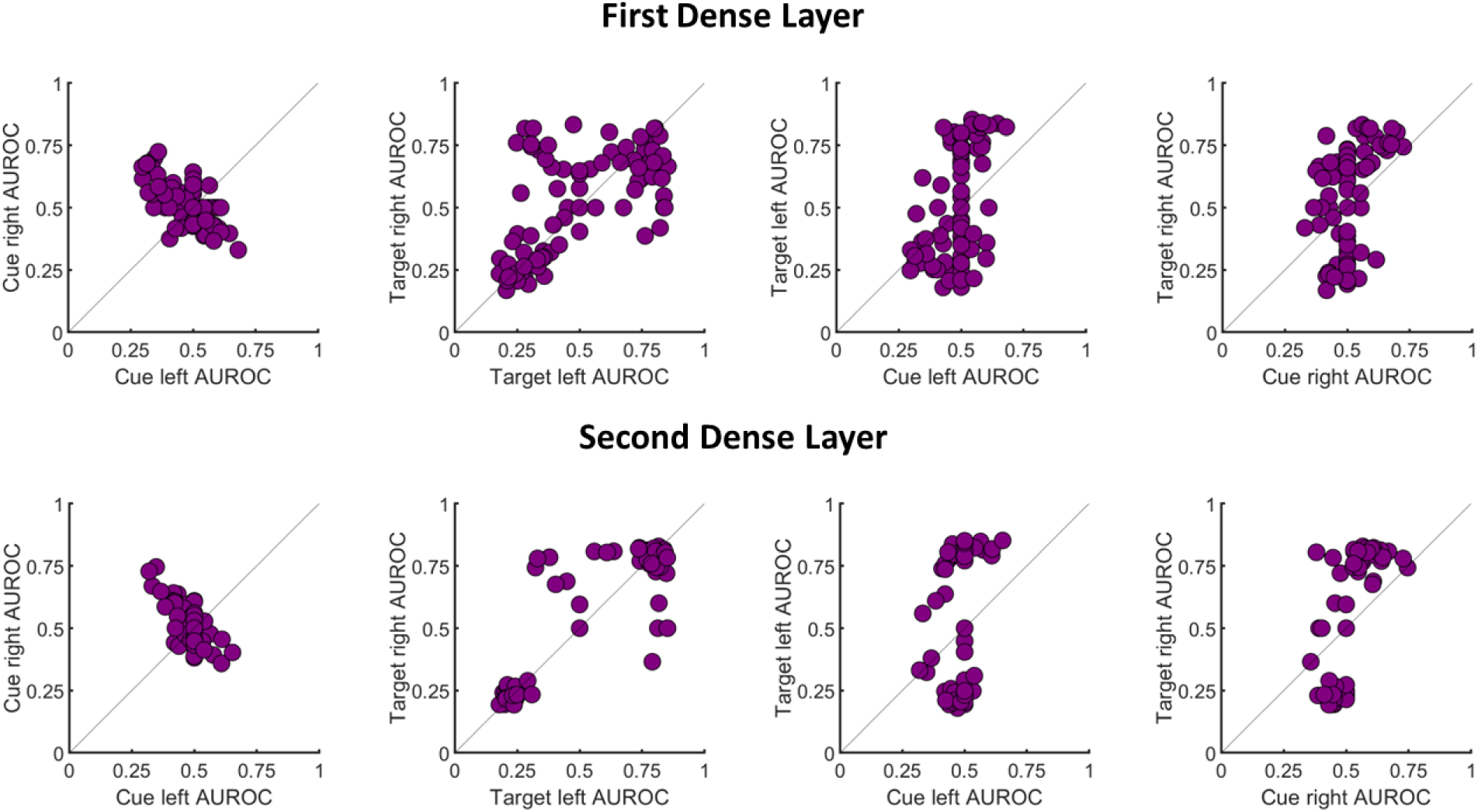
The effect of adding another dense layer. The rest of the network architecture was kept the same as the network in the main text. The first dense layer has 256 neurons while the second dense layer has 100 neurons. The second dense layer doesn’t look qualitatively different than the first dense layer or most other dense layers reported throughout the main text and the supplement.

#### S.12(b) The Effect of Depth of the Network: Adding Another Convolution Layer

Increasing depth via convolutions does increase the extent of local target-cue integration in the flat layer but not across locations (See Supplementary Figure S14). Adding another convolution layer changes the output correlation with the BIO (0.732 for four convolution layers).

**Supplementary Figure S14.**
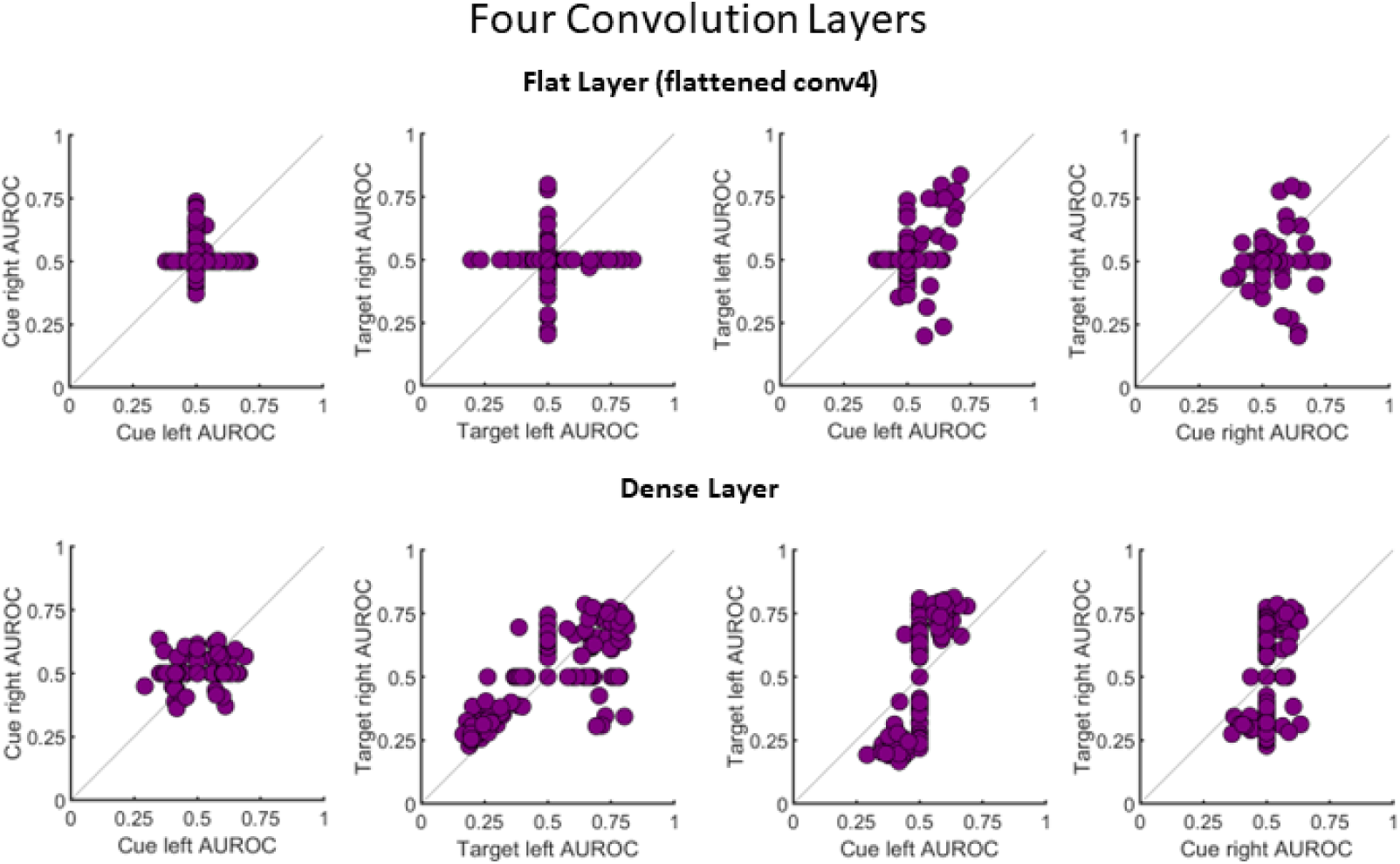
The effect of adding another convolution layer. The rest of the network architecture was kept the same as the network in the main text. The flat layer refers to the flattened last convolution layer. Adding another convolution layer increases the target and cue sensitivity of the TC neurons in the flat layer (last two columns; contrast with Figure 4 in the main text), but not the integration across location (first two columns; compare with Figure 5 in the main text).

### S.13 Effect of Increasing Convolution Stride Size

Increasing the stride size of all convolutions from 3 to 5 reduced the network performance to 0.71 while increasing the local target-cue integration on the left location but not the right in the flat layer (Supplementary Figure S15). The output correlations with the BIO were also reduced to 0.625. These results are likely due to the loss of resolution due to the strides.

**Supplementary Figure S15.**
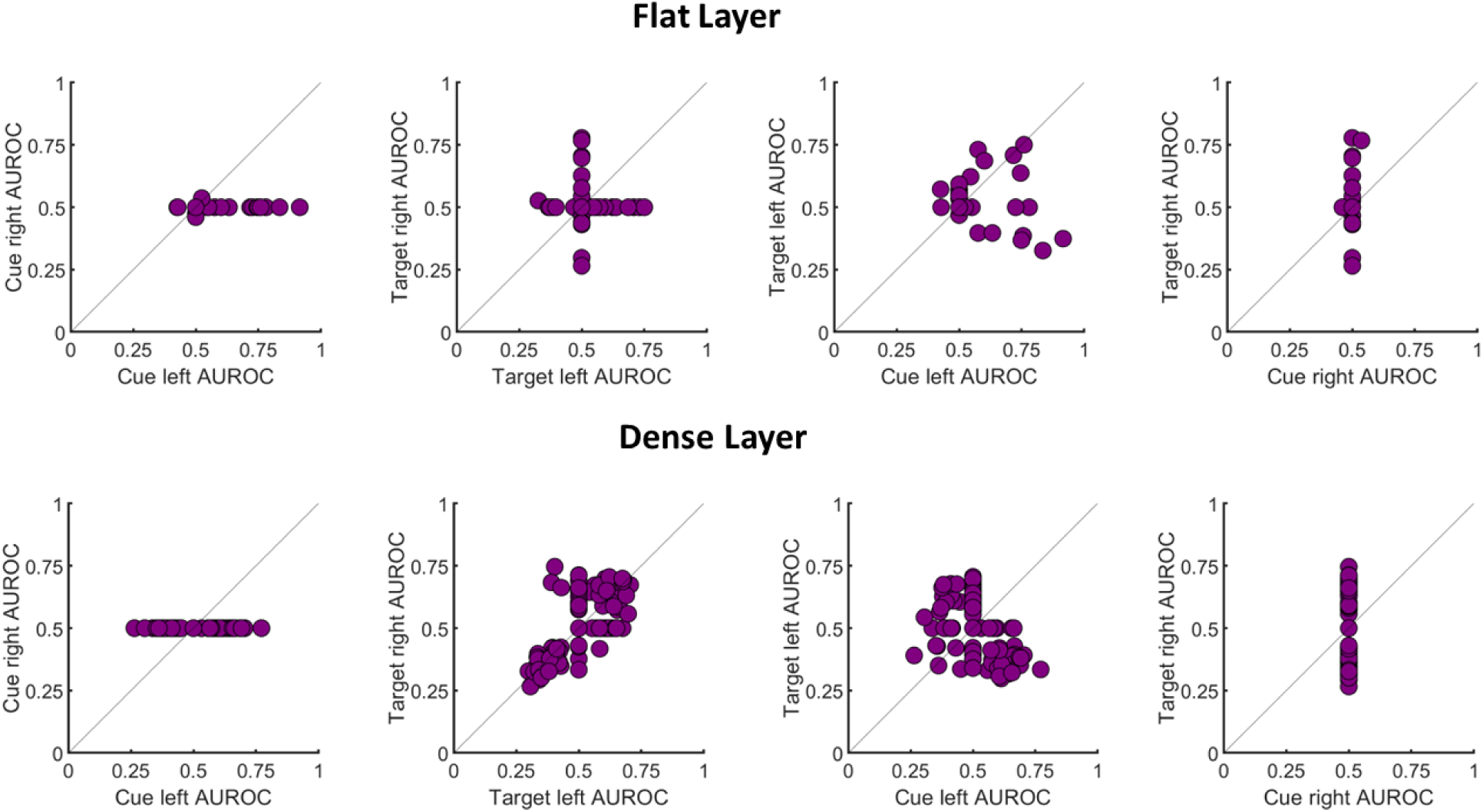
The effect of increasing the stride of convolution layers. The rest of the network architecture was kept the same as the network in the main text, but the stride of all convolutions was increased from 3 to 5. The local target-cue integration on the left location was improved in the flat layer (third column, first row. Compare with Figure 4 in the main text). However, the cue-integration and the target-cue integration on the right in the dense layer suffer, perhaps due to the loss of resolution due to downsampling (second row, first and last columns. Compare with Figures 4 and 5 in the main text).

### S.14 Effect of Increasing Kernel Size

We tried kernel size 5 besides the existing networks that have kernel size 3. The network performance increases (Proportion correct = 0.762), the amount of local target-cue integration increases in the flat layers but not the integration across locations (Supplementary Figure S16), and the output correlation with the BIO also increases (0.77).

**Supplementary Figure S16.**
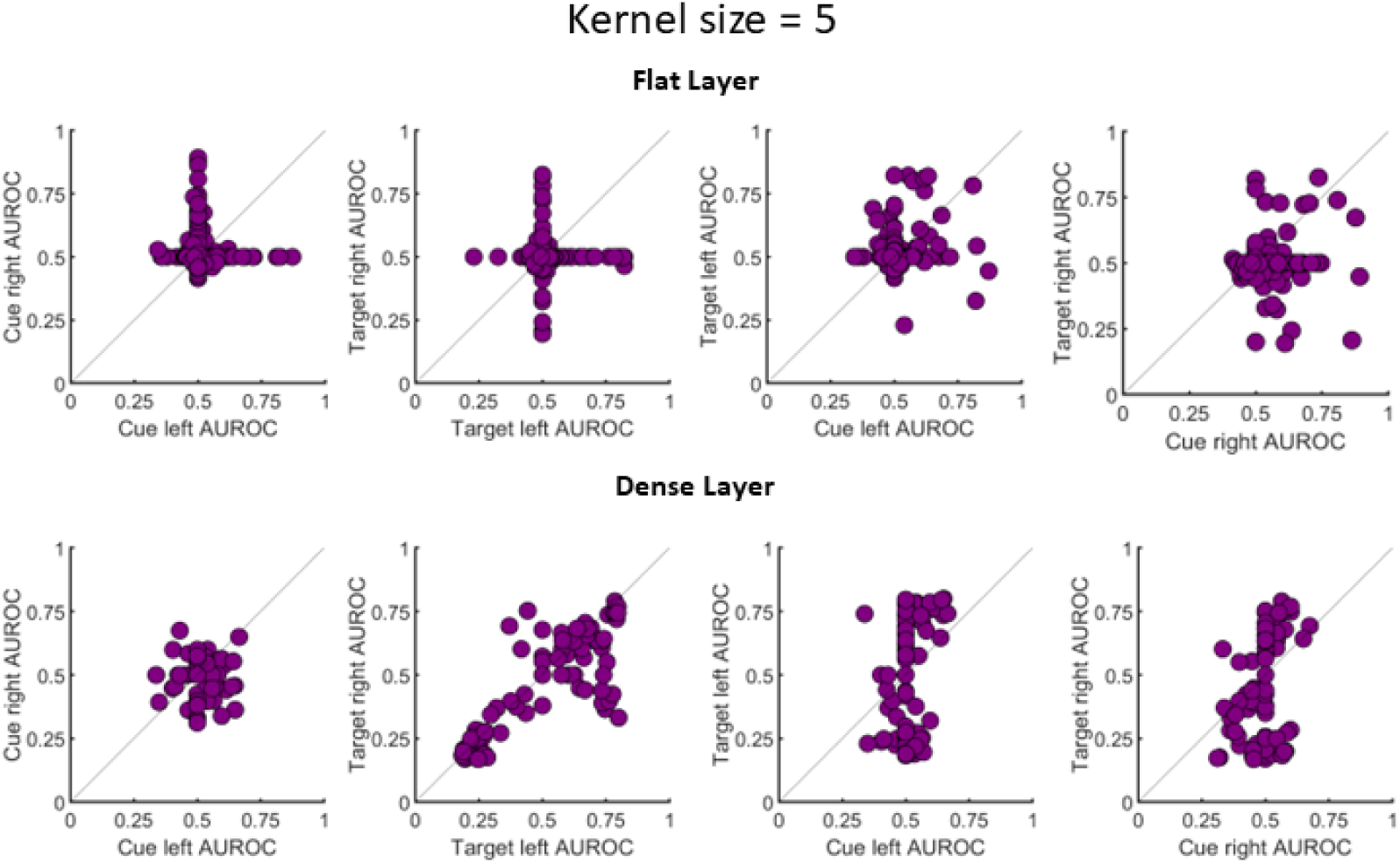
The effect of increasing the kernel size of convolution layers. The rest of the network architecture was kept the same as the network in the main text, but the kernel size of all convolutions was increased from 3 to 5. The local target-cue integration was greater in the flat layers (last two columns. Compare with Figure 4 in the main text), but not the integration across the two locations.

### S.15 Generalization across External Noise Amplitudes

While our analysis uses networks trained with noise to bring their performance in a comparable range to human subjects in ^1^, the point of this noise fitting is to make a visual comparison between humans and the models easier. The results, however, do not depend on the noise. The model still shows cueing in the absence of pixel noise (because there is noise in the target and distractor angles – the same angle noise as in the images shown to humans). When there is no noise, the model achieves an accuracy of 0.78 with a difference of 0.15 in the valid and invalid hit rates. The correlation with BIO is 0.88 at the output level, and in the dense layer, it achieves a mean absolute correlation of 0.62 and a max absolute correlation of 0.86 with the BIO. The max absolute in the flat layer is 0.55, and the mean absolute is 0.19.

### S.16 CNN Predictions with Central Cue

We trained a network where the 80 % predictive cue was a central line that pointed in the direction of the likely target location, instead of the peripheral box cue used in most of the paper. The model obtained a valid hit rate of 0.75, an invalid hit rate of 0.66, and a false positive rate of 0.22. The scatterplots for location and cue-target integration are depicted in Supplementary Figure S17.

**Supplementary Figure S17.**
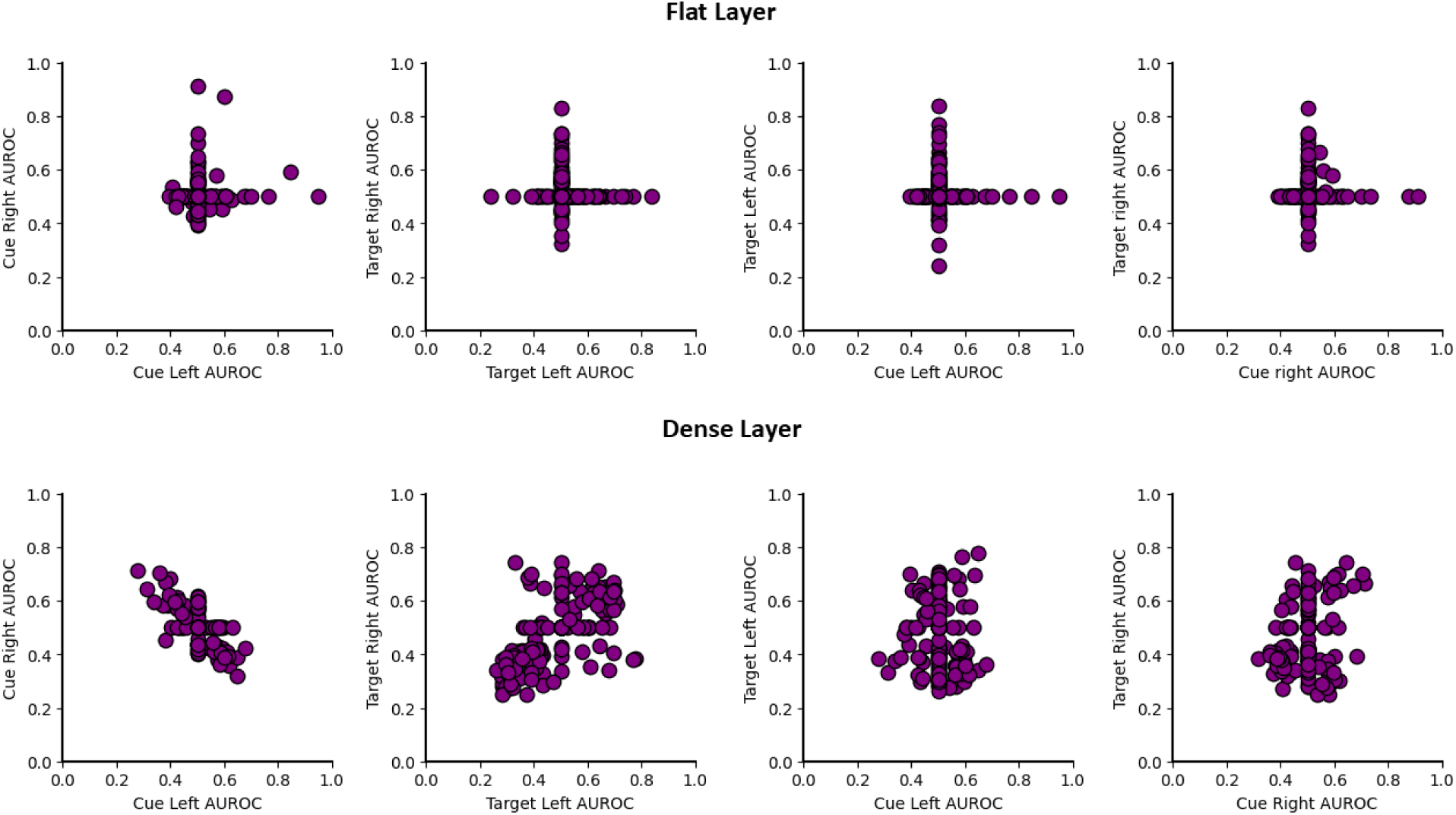
The effect of central cues. Top row: Scatter plots for cue right AUROC vs cue left AUROC, Target right AUROC vs target left AUROC, Target left AUROC vs Cue left AUROC, and Target right vs cue right AUROC for the flat layer for the model trained with central cues. Second row: Scatter plots for cue right AUROC vs cue left AUROC, Target right AUROC vs target left AUROC, Target left AUROC vs Cue left AUROC, and Target right vs cue right AUROC for the dense layer for the model trained with central cues.

### S.17 VGG-16 Pretrained on ImageNet with the Dense Layer Retrained on the Posner Cueing task

Because the visual system in the brain works on a myriad of tasks, we assessed whether a pre-trained network with more ecologically valid tasks and re-trained for the cueing task would show similar results to the smaller networks only trained for the cueing tasks. We used a VGG-16 model pretrained on ImageNet, froze its first three blocks, comprising seven convolution layers and three pooling layers, flattened the output of the last pooling layer, and trained a dense layer of size 500 followed by the output layer of size 2 on our task. The weights of the ImageNet pre-trained model were downloaded from the official implementation provided by TensorFlow. The network achieved a valid hit rate of 0.71, an invalid hit rate of 0.60, and a false positive rate of 0.19. The network resulted in similar mechanisms to the smaller network in the main text (see Figure S18 dense layer plots, Figure S19 for a table of neuron types): cue location-opponent units, summation units, and all combinations of TC neurons (T+C+, T+C-, T-C+, T-C-, last row, rightward two plots, see Fig. S18 and S19). There were some differences with the smaller network trained only on the cueing task reported in the main text. We observed that in the VGG model, even the earlier layers show a small percentage of neurons with cue-target Integration (the smaller CNN trained only the cueing task showed none). The first pooling layer has 0.06% neurons tuned jointly to the cue and the target, compared to 0.28% in the second pooling layer, 0.38% in the flat layer, and 39.2 % in the dense layer.

Additionally, as opposed to the small networks used in the main text, the dense layer in the retrained VGG model shows neurons with perfect detection of the cue (cue AUROC = 1 or 0), while the unit AUROC for the smaller network trained only on the cueing task did not exceed ∼0.8. Figure S19 shows the percentages of different neuron types for the VGG model’s dense layer.

**Supplementary Figure S18.**
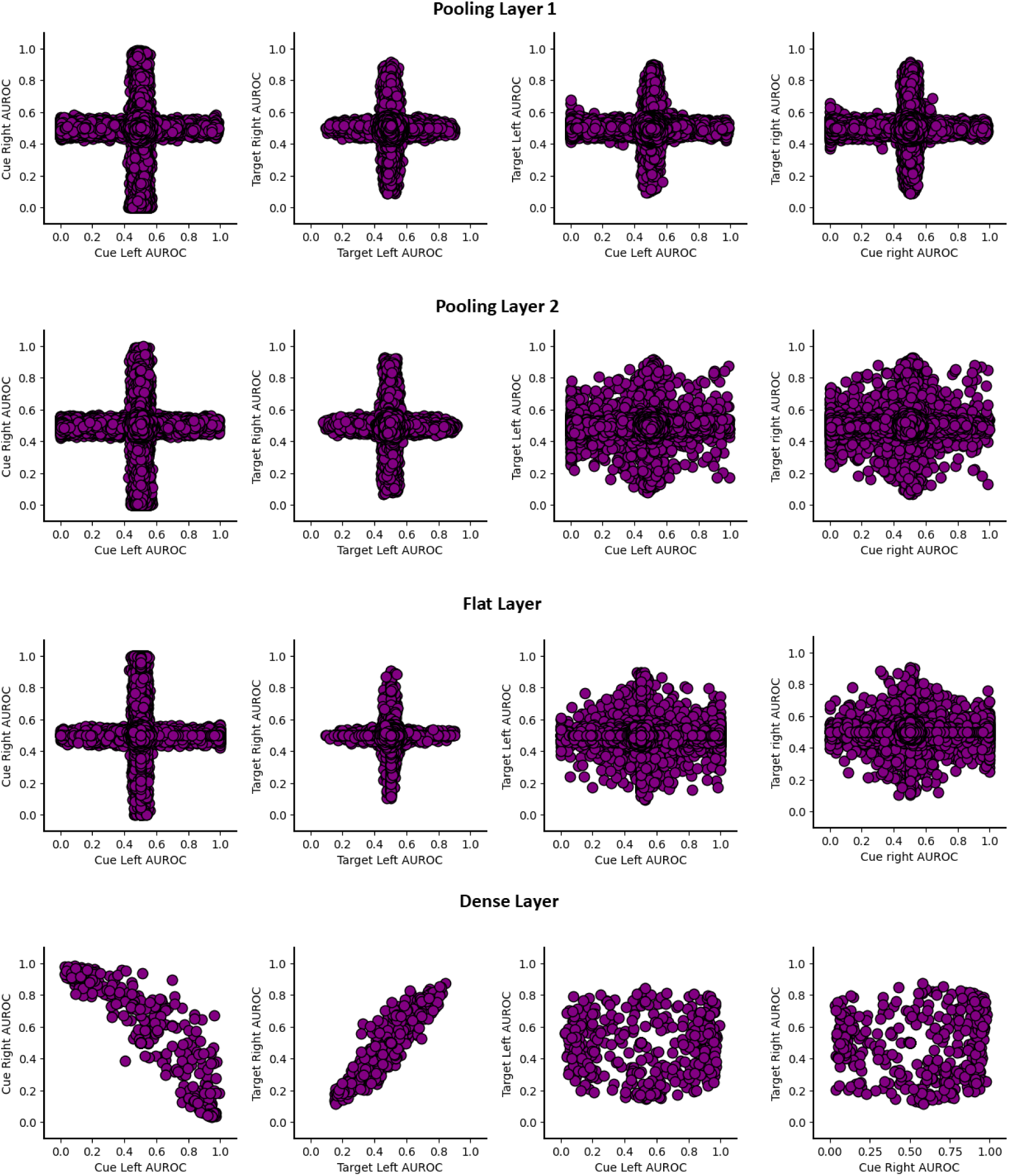
Pretrained VGG-16retrained on our task. Each row corresponds to one layer. For each of the first three blocks, we compute the AUROCs for the pooling layer of that block. The pooling layer of the third block is flattened and fully connected to the dense layer, as a result of which the third pooling layer is the flat layer in this model. First column: Scatterplots of VGG-16 layer units’ cue right AUROC vs cue left AUROC (integration of cue across locations). Second column: Scatterplots of VGG-16layer units’ target right AUROC vs target left AUROC (integration of target across locations). Third column: Scatterplots of VGG-16 layer units’ target AUROC vs cue AUROC for the left location. Fourth column: Scatterplots of VGG-16 layer units’ target AUROC vs cue AUROC for the right location.

**Supplementary Figure S19.**
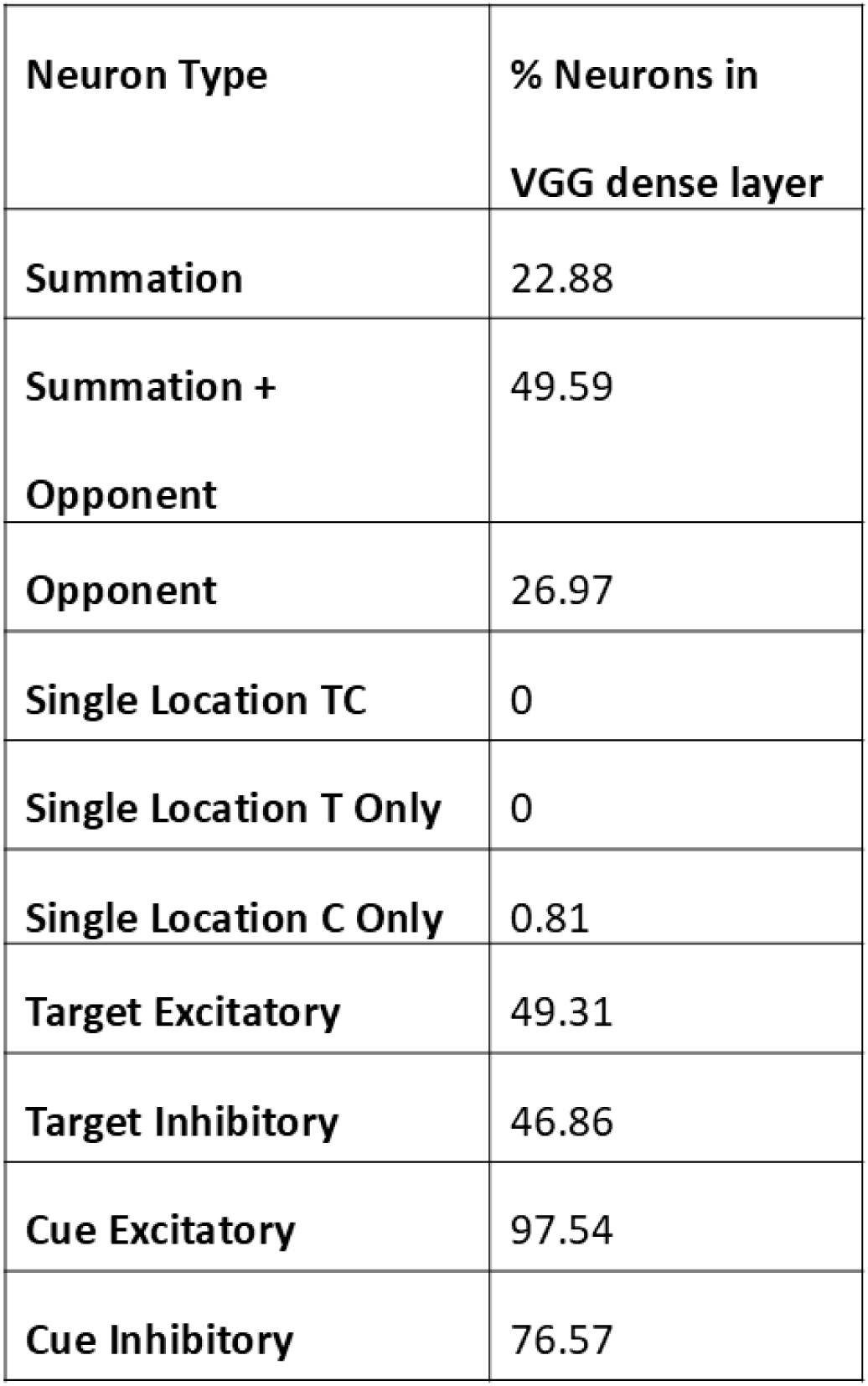
Neuron types in the dense layer of the pretrained VGG-16 retrained on our task. Excitatory and Inhibitory neurons include the ones that are excitatory at one location and inhibitory at the other and can thus add up to more than 100%.

### S.18. Target, Cue, and Location Sensitivity for Ipsilateral and Contralateral Visual Fields to Mice Superior Colliculus Cells

Here we replot the SC cells figures in terms of the stimulus locations relative to the recorded SC neuron hemifield (ipsilateral and contralateral, Figure S20). This is a common way to display neuron results in neurophysiological studies and is included for reference. The main text plots the data in terms of left and right locations to facilitate comparisons to the CNN theoretical results (the CNN does not have hemispheres that map to opposite visual fields).

**Supplementary Figure S20.**
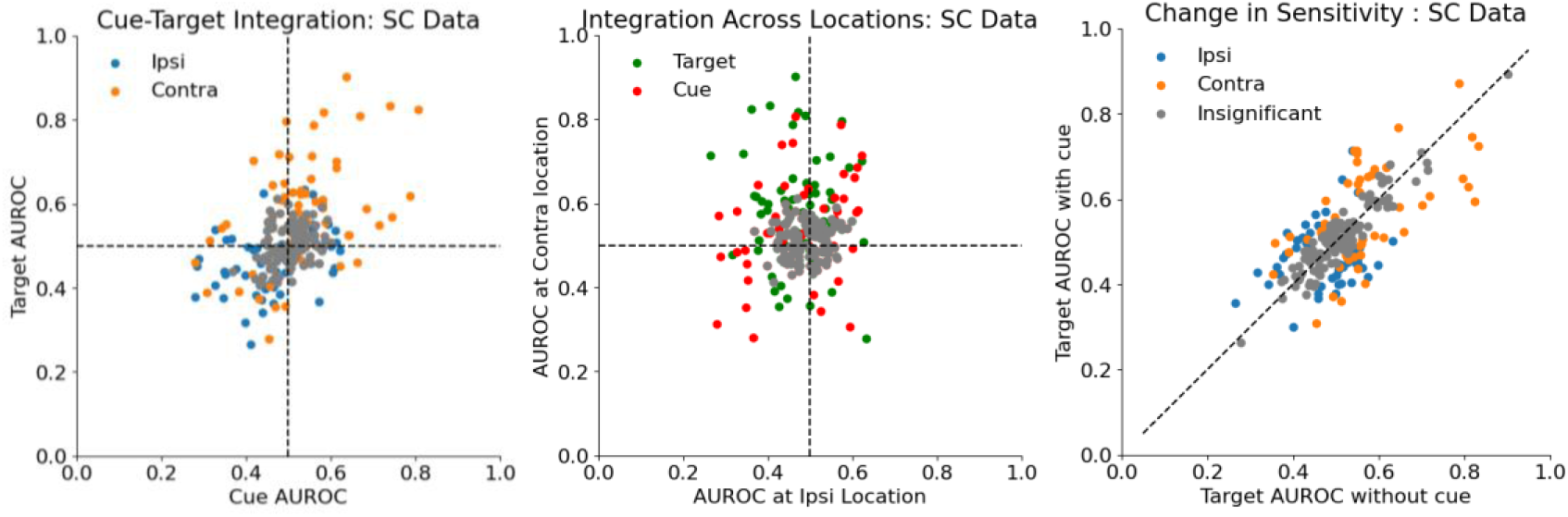
a) Cue and target sensitivity of 109 mice superior colliculus neurons during an orientation change detection task with a 100 % valid cue and another condition with no cue (Wang et al., 2022); Gray points indicate neurons with insignificant cue and target AUROCs. b) Cue and target sensitivity for each location for SC neurons; Gray points indicate neurons with insignificant AUROC at both locations. c) Target sensitivity (AUROC_t-d_) with and without cue for SC cells; In the main text (Figure 7), we report left vs right AUROCs, while here we report the stimulus locations relative to the recorded SC neuron hemifield (ipsilateral and contralateral).

### S.19 Percentage of Neuron Types for CNN and Superior Colliculus Mice Cells

Here, we show the percentage of neuron types out of the 109 SC cells analyzed and the responsive CNN units in the flat and dense layers (Figure S21).

**Supplementary Figure S21.**
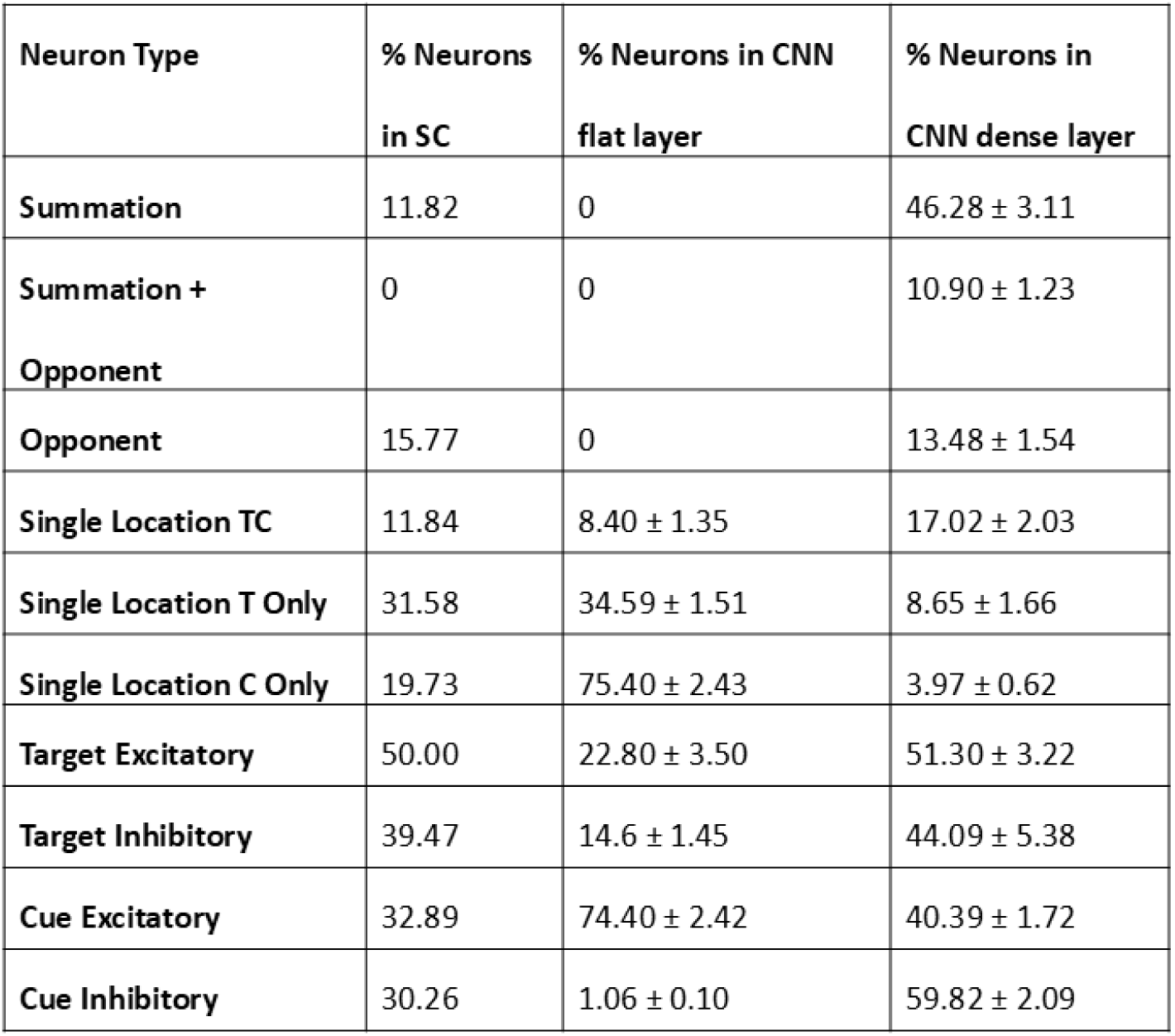
Neuron types (first column) and their prevalence in the mice SC data (second column) and in the CNN’s flat layer (third column) and the dense layer of the models (fourth column) trained on the task from Wang et al. Note that the percentages are calculated out of the responsive neurons. This is different from Figure 3e, which reports percentages out of all neurons in a layer, including non-responsive neurons. Excitatory and inhibitory neurons include the units that are excitatory at one location and inhibitory at the other, and can thus add up to more than 100%.

### S.20 The Effect of the Length of Training

Training for more epochs does not change the overall trend of the AUROC correlations. However, during later epochs, there are fewer neurons that are untuned to either the target or the cue, and thus more points in the scatterplot move away from the (0.5, 0.5) point on training longer (Figure S22).

**Supplementary Figure S22.**
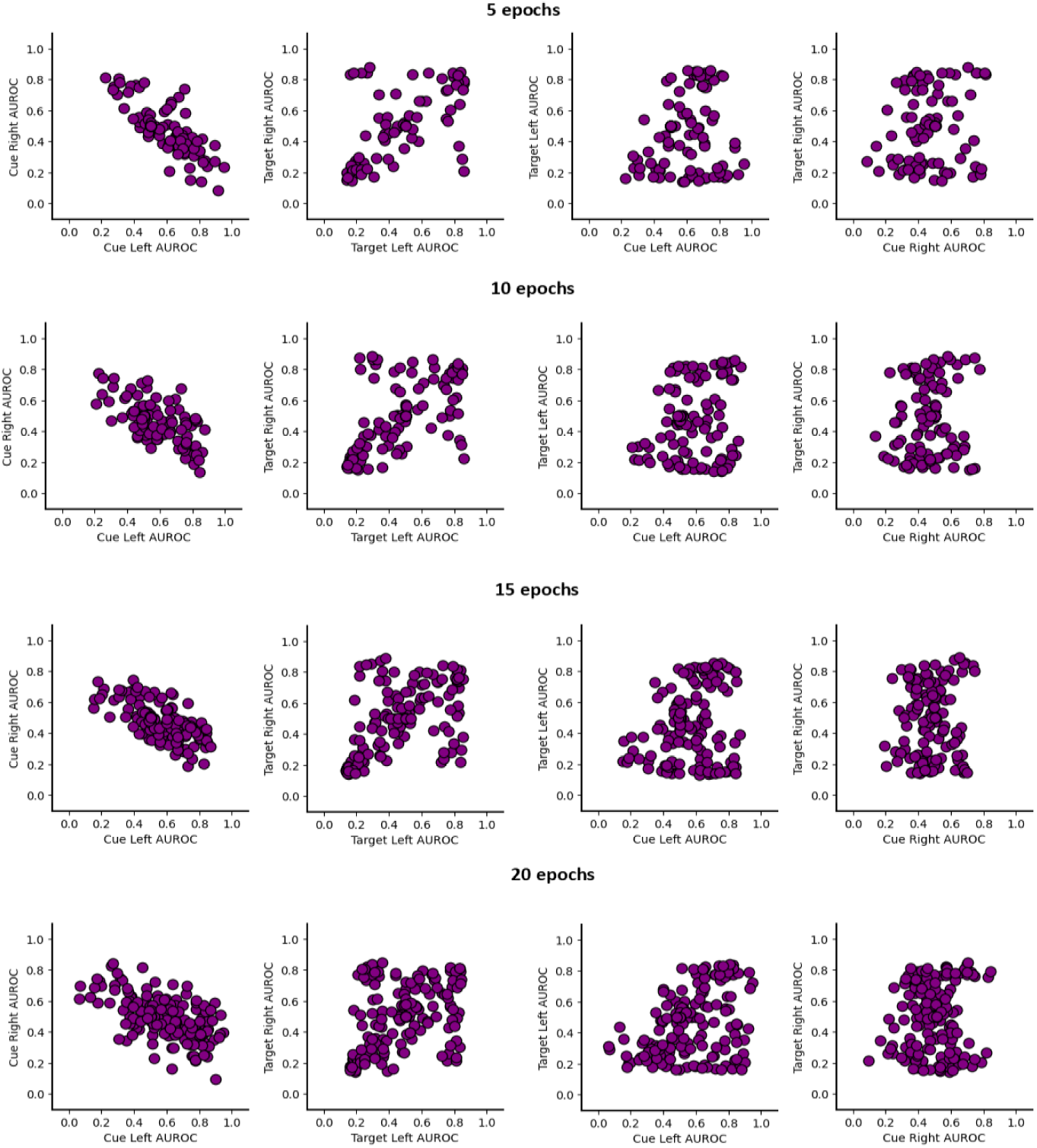
The effect of the length of training. Each row presents the dense layer AUROCs as a function of the number of epochs trained for. First column: Scatterplots of cue right AUROC vs cue left AUROC (integration of cue across locations). Second column: Scatterplots of target right AUROC vs target left AUROC (integration of target across locations). Third column: Scatterplots of target AUROC vs cue AUROC for the left location. Fourth column: Scatterplots of target AUROC vs cue AUROC for the right location.

### S.21 Visualizing the Stages of the CNN

Supplementary Figure S23 shows a summary visualization of our analyses of the properties of the neurons at the different layers in terms of their tuning to the target, cue, joint tuning, and location integration. For simplicity, the visualization shows the five activation maps in each layer (out of 8, 24, and 32 activation maps per layer) that showed the highest responsiveness. Below each layer, we indicate the corresponding functional computational stages.

**Supplementary Figure S23.**
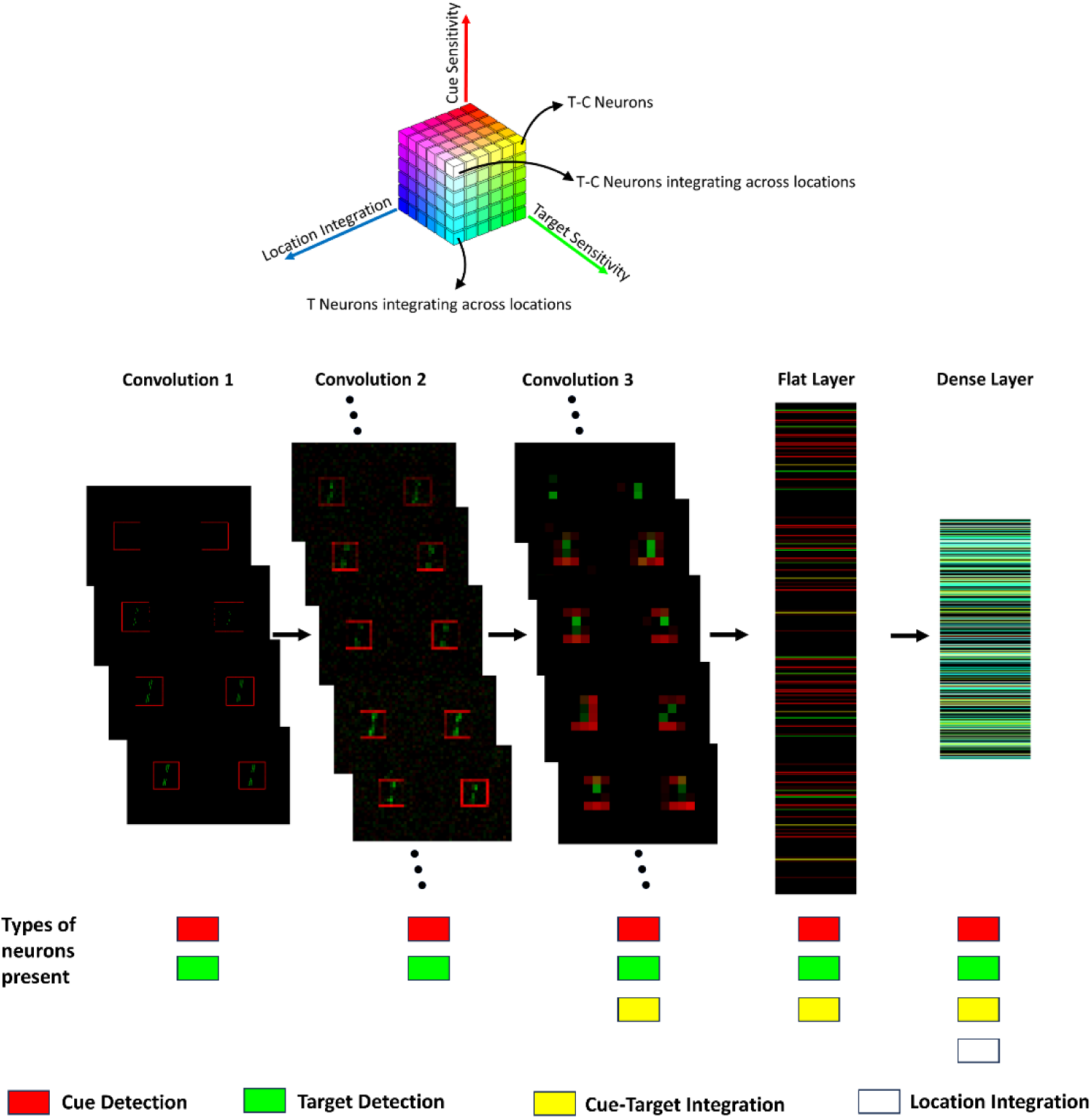
A layerwise map of neuron types present in the network. Each neuron is represented by an RGB triplet, where the red value depicts the cue sensitivity (Max{|AUROCc-nc left – 0.5|,|AUROCc-nc right – 0.5|}x 2) of the neuron, the green value depicts the target sensitivity (Max{|AUROC_t-d_ left – 0.5|,|AUROC_t-d_, right– 0.5|}x 3) and the blue value depicts the location integration of the neuron (|AUROC_t-d_ left– 0.5|x|AUROC_t-d_, right – 0.5|x 3).

### S.22 Neuron-BIO Correlations Layerwise Visualization

We extracted the log-likelihood ratio from the BIO for 1000 unseen test images (500 target absent, 400 valid, 100 invalid) and used the same images for the CNN, obtained the response of each neuron to these 1000 images, and correlated each neuron’s trial-to-trial responses with the BIO’s log-likelihood ratios for those trials. For the CNN’s output layer, we used the SoftMax response. Figure S24 shows a visualization of the BIO log-likelihood ratio and CNN correlations across the different layers.

**Supplementary Figure S24.**
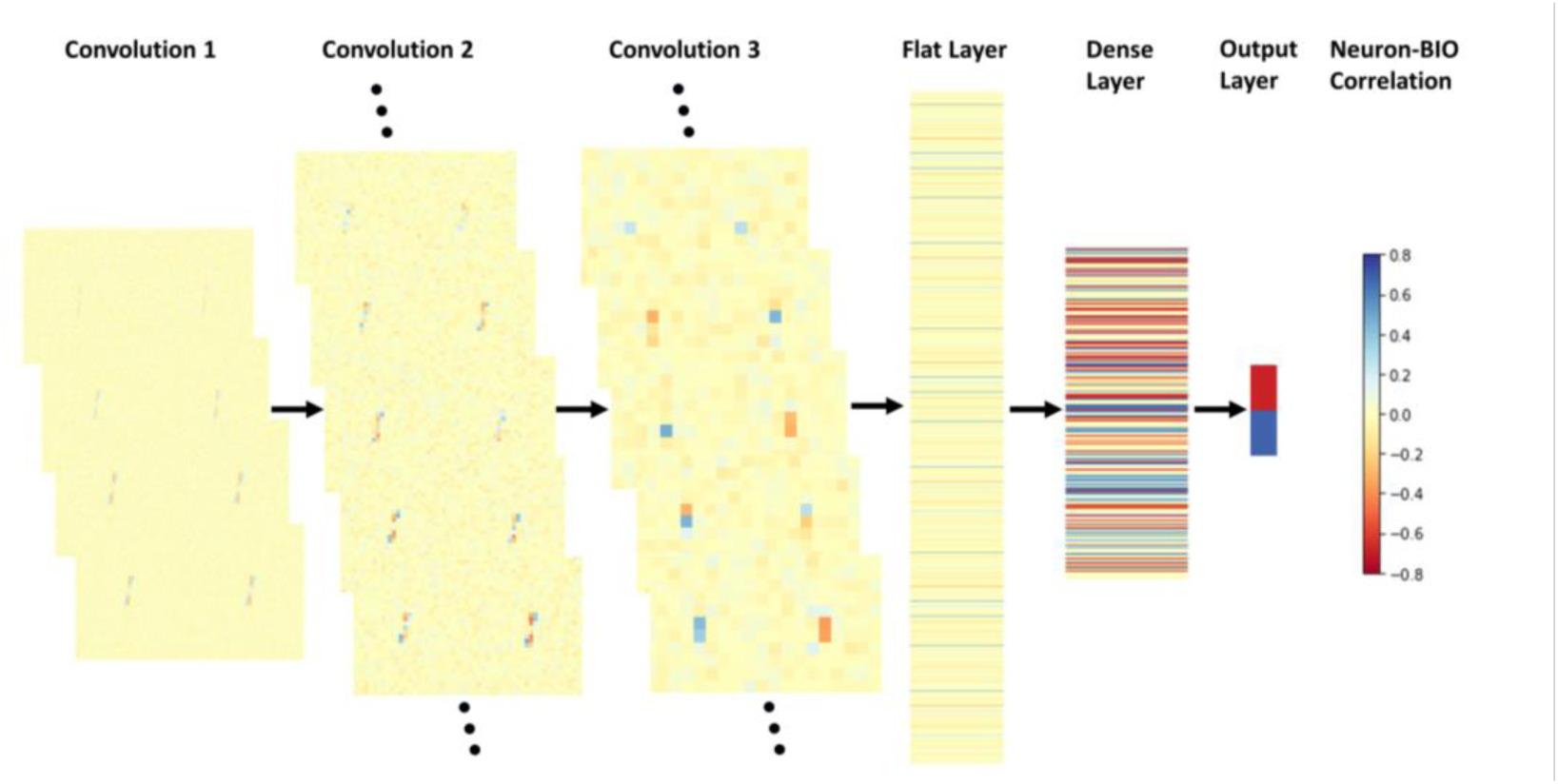
Layerwise Neuron-BIO Correlations. For each neuron, we calculated the trial-to-trial correlation with the BIO on a test set. A heatmap of these correlations is presented for each layer. For the first convolution layer, all responsive maps are shown. For the next two convolutions, the five activation maps with the highest mean absolute correlations are shown. For the flat layer, a middle strip of 800 neurons with higher correlation is shown, outside which the correlations are small in magnitude. The flat and the dense layer (256 neurons) are approximately equal in scale. The output layer only has two neurons, whose size has been amplified for visualization purposes.

### S.24 Statistical Testing for AUROCs for mice SC neurons with Non-Parametric Bootstrap

The bootstrap resampling test is considered a robust test because it does not assume parametric distributions. For our application to the SC mice data, it entails sampling with replacement a neuron’s activity from the original data. We sampled with replacement neuronal data for 40 signal-present and 40 signal-absent trials (the number of trials in the data set). We then compute the AUROC for the resampled data. This is repeated 10,000 times. The distribution of the 10,000 resampled AUROC represents a distribution of the sample mean of the AUROC, and its spread is an estimate of the error in the sample mean. To assess whether an AUROC is statistically significant, we determine whether 0.95 (p=0.05) of the resampled distribution is above or below an AUROC = 0.5. Figure S26 shows example distributions of bootstrap AUROC estimates for a T+ neuron, a T- neuron, a C+ neuron, a C- neuron, and a neuron with non-significant AUROC.

To adjust for increasing Type I errors from multiple tests, we use a False Discovery Rate (FDR) correction based on the Benjamini-Hochberg (BH) procedure^25^. The FDR procedure consists of ordering all the p-values from all neurons in ascending order and then calculating a correct p-value (q) based on the total number of tests (M) and the rank of the current test: q = p M/rank. The q values are compared to the significance value (0.05). All q < 0.05 are considered statistically significant. Our statistical testing procedure is the same test used by the original paper reporting the SC data (Wang et al., 2024), except they did not use FDR correction, making our statistical test more conservative (reducing the probability of Type I error).

**Supplementary Figure S26.**
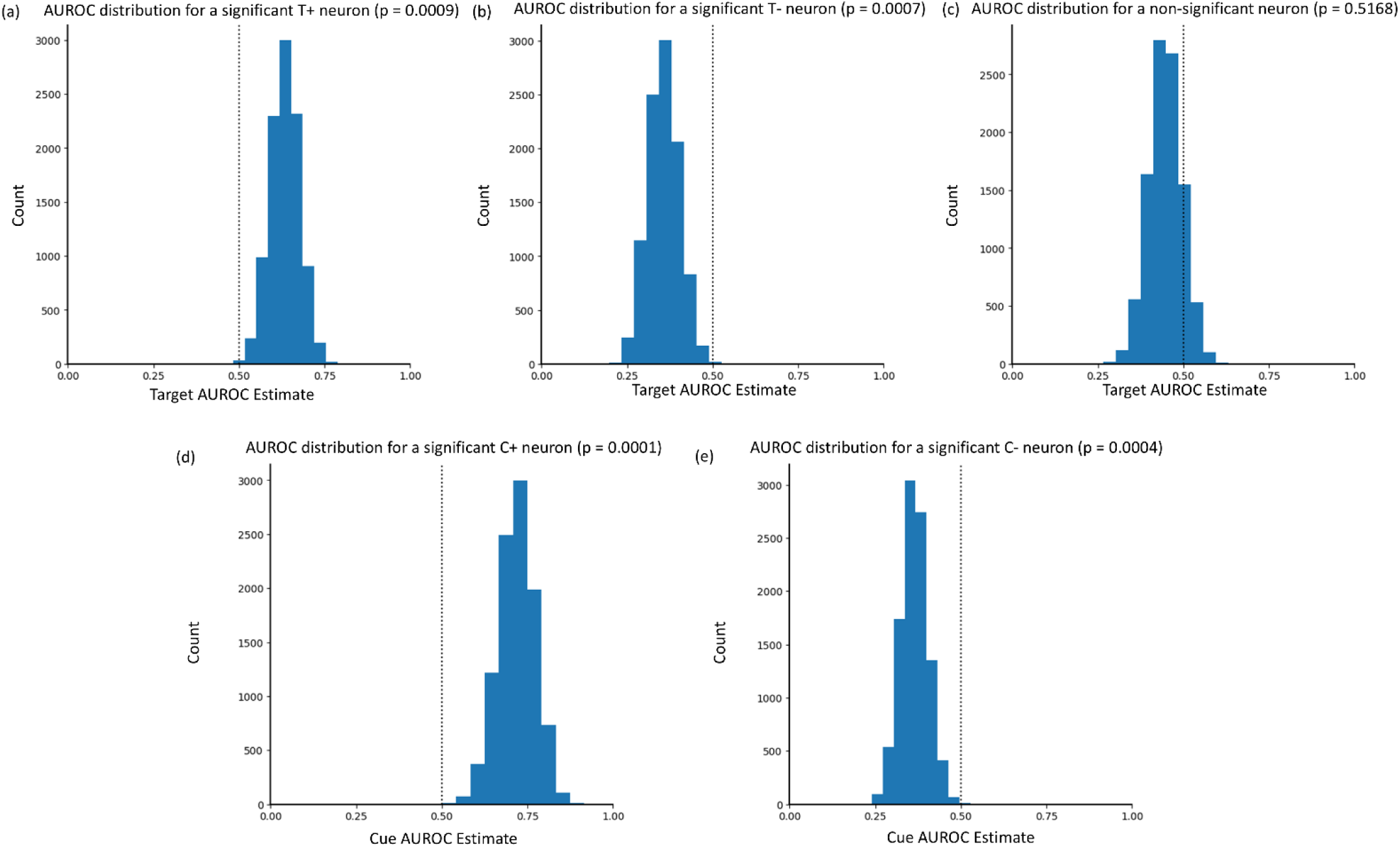
Bootstrap AUROC distributions for a (a) T+ neuron, (b) a T- neuron, (c) a non-significant neuron, (d) a C+ neuron, and (e) a C- neuron. The dotted line denotes an AUROC of 0.5. The empirical p values are corrected using FDR before being compared to p= 0.05 for statistical significance.

### S.25 Estimating Mechanism Contributions to the cueing effect using linear regression

In Figure 6k in the main text, we show a simple schematic of the three dense neuronal mechanisms, location-summation, location-opponent, and summation + opponent, that receive inputs from flat T, C, and TC neurons, and how they feed information into the output layer.

To calculate the regression from the dense mechanisms to the output neuron 1, we fit one linear regression to predict the output neuron’s response based on the responses of all dense neurons classified as one of the three mechanisms. The arrow widths depicted in Figure 6k were then calculated as the mean absolute regression weights of each dense neuron mechanism.

Separate training and test sets of responses were obtained by getting the responses of each layer on a training set of 10000 images and a testing set of 2000 images, with cue validity 80% and the target present 50% of the time.

To calculate the regression from flat to dense neurons, we fit 3 linear regressions, one for each dense mechanism. All neurons in the flat layer that were T, C, or TC neurons at either location were used as the predictor variables. The predicted variable was the average response of all dense neurons from one category type. We used the average across neurons to simplify the regression. Otherwise, our analysis would require predicting the responses of each dense neuron separately (> 1000 regressions). We thus ran three regressions to predict the average responses of location summation, location opponent, and summation + opponent. The arrow widths in Figure 6k represent the mean absolute regression weights from each flat neuron type to each dense layer mechanism.

### S.26 Approximating the CNN with a linear combination of location summation and opponent cells (and no summation + opponent cells)

An important finding is that the summation + opponent CNN cells, which had the highest regression weights to the output neuron (see Section S.25), were absent in the mice SC. The finding motivated us to assess how well a linear regression trained on just the location summation and location opponent dense layer neurons (but no summation + opponent) could account for the CNN output responses. We found that this regression model explained 86% of the variability of the output neuron and obtained a cueing effect that was 92% of that of the original CNN.

